# The K18-hACE2 mouse model of SARS-CoV-2 infection to illustrate the role and response of the vasculature in neurotropic viral infection

**DOI:** 10.1101/2025.02.07.637145

**Authors:** Simon De Neck, Rebekah Penrice-Randal, Leandro Xavier Neves, Frauke Seehusen, Parul Sharma, Barbara Helminger, Adam Kirby, Daniele Mega, Maximilian Erdmann, Martina Zanella, Michelle Reid, Edward Emmott, Giuseppe Balistreri, Alaa Othman, Udo Hetzel, James P. Stewart, Anja Kipar

## Abstract

Severe Acute Respiratory Syndrome Coronavirus 2 (SARS-CoV-2) primarily affects the respiratory tract and lungs; however, the associated disease, coronavirus disease 2019 (COVID-19), can involve the central nervous system (CNS) in both its acute and long-term (Long COVID) clinical manifestation. The pathomechanisms underlying neurological impairments in COVID-19 are not yet fully understood, hence experimental studies to clarify the direct effect of SARS-CoV-2 in the brain can provide further insight.

In the present study we used the K18-hACE2 model, intranasally challenged with SARS-CoV-2 ancestral and Delta isolates at low or medium doses, to address the hypothesis that the inflammatory response raised in the brain of infected mice is secondary to neuronal infection.

Our data confirmed that the virus reaches the brain even after low dose (10^2^ PFU/mouse, Delta isolate) infection where it targets the neurons without overt neuropathic effect, sparing the blood vessels. *In situ* investigation of the resulting inflammatory response showed the recruitment of leukocytes via postcapillary venules, with their accumulation in the perivascular space and occasional migration into the neuroparenchyma, without targeting and/or damage to the vessel wall. These changes were reflected in the brain transcriptome and proteome which showed positive enrichment of pathways and up-regulation of genes involved in the inflammatory response including the recruitment (including adhesion and migration) and activity of leukocytes. Additionally, morphological and transcriptome/proteome changes suggest minimal associated blood-brain barrier dysfunction. The brain metabolome and lipidome showed minimal changes; these were consistent with oxidative stress and inflammatory and immune/antiviral responses.

The results obtained from our model indicate that SARS-CoV-2 infection of the neurons can result in limited neuroinflammation. These data can help to understand more fully the reaction of the CNS in COVID-19 patients, and neurotropic virus infections in general.

## Introduction

SARS-CoV-2 primarily affects the respiratory tract and lungs; however, other organ systems including the CNS can also be involved in the associated disease, COVID-19 [1, 2]. Neurological and neuropsychiatric symptoms have been observed and remain a concern not only in patients with acute COVID-19 but also with Long COVID [2–6]. The pathomechanisms underlying neurological impairments are still not fully understood. Different mechanisms have been proposed including direct viral CNS infection, although this is considered a very rare event [2, 7].

Several publications report pathological changes in the brain of fatal COVID-19 cases, summarised and interpreted as “a pattern of cerebrovascular pathology and microglial-predominant inflammation” [8, 9]. Histologically, (sub)acute infarcts and hypoxic-ischaemic injury, microthrombi, acute (micro)haemorrhages, microglial activation and occasional microglial nodules as well as mild lymphocyte-dominated perivascular, parenchymal and meningeal inflammatory infiltrates have been described [8–16]. Interestingly, SARS-CoV-2 was either not detectable in the brain, or detected in minimal amounts (RNA, protein), as determined by various approaches [7–9]. However, the debate on the frequency and extent of SARS-CoV-2 infection of the brain in COVID-19 patients has not come to full conclusion.

To enhance our understanding of CNS involvement during SARS-CoV-2 infection, suitable animal models can be used. For example, several groups including ours have previously shown that intranasal challenge of K18-hACE2 mice (expressing human (h)ACE2 under the K18 promotor [17]) with SARS-CoV-2 (ancestral and variant of concern (VOC) isolates) results in rather consistent spread to the brain [18–33]. However, neurological signs and morphological features consistent with neuronal cell death were only reported with high viral doses (10^6^ PFU/mouse) [18]. Viral antigen and RNA was detected in neurons of infected brains [18, 19, 23, 26–31, 33], with astrocytes, microglia [23], endothelial cells [30] and macrophages [28] occasionally reported as infected too. Despite widespread neuronal infection, only a limited tissue reaction was observed in the brain parenchyma, consisting of mild (meningo)encephalitis, characterised by (peri)vascular leukocyte infiltrates, microgliosis and astrogliosis [18–20, 23, 27–29, 31].

Several modes of virus entry into the CNS have been discussed, including transsynaptic neuronal/retrograde axonal transport via nerves [34–36]. This is modelled in K18-hACE2 mice in which after intranasal challenge SARS-CoV-2 mainly reaches the brain via the olfactory system and spreads among neurons [18, 19, 37–39]. Therefore, and based on our previous work [19], we hypothesised that the vessel-centred inflammatory reaction in the brain of SARS-CoV-2 infected K18-hACE2 mice is a response to the neuronal infection, rather than a targeting of the blood vessels by the virus and/or leukocytes. To test this hypothesis, we used K18-hACE2 mice infected with an original SARS-CoV-2 Liverpool strain and a Delta variant to explore the role of cerebral blood vessels in brain infection, with special emphasis on leukocyte recruitment, integrity of the vascular wall and blood-brain barrier (BBB) function.

## Material and Methods

### Viruses

UK isolates of SARS-CoV-2, i.e. the original “Liverpool strain” (hCoV-2/human/Liverpool/REMRQ0001/2020) and a Delta (VOC) isolate (B.1.617.2 hCoV-19/England/SHEF-10E8F3B/2021; GISAID accession number EPI_ISL_1731019) were used for experimental infections [19, 40]. Both have previously been shown to spread to the brain with high frequency and induce a mild non-suppurative encephalitis and meningitis in a proportion of affected K18-hACE2 mice [19, 41].

### Animals and virus infection

Animal work was approved by the local University of Liverpool Animal Welfare and Ethical Review Body and performed under UK Home Office Project Licence PP4715265. It was undertaken in accordance with locally approved risk assessments and standard operating procedures. The study was undertaken on mice carrying the human ACE2 gene under the control of the keratin 18 promoter (K18-hACE2; formally B6.Cg-Tg(K18-ACE2)2Prlmn/J) purchased from Charles River Laboratories (Wilmington, Massachusetts, USA). Mice were maintained in biosafety level 3 facilities under SPF barrier conditions in individually ventilated cages.

The study was undertaken on cohorts of K18-hACE2 mice that were part of experimental infections undertaken for other projects (cohorts 1.1-1.4) or specifically recruited for the present study (cohorts 2.1-2.3) (Supplemental Table S1).

Female and male K18-hACE2 mice, aged 6-8 weeks, were intranasally infected with different doses of the SARS-CoV-2 Liverpool strain or a Delta isolate, or mock-infected, and euthanised at different time points post infection.

For infections, mice were anaesthetized lightly with isoflurane and inoculated intranasally with 50 µl sterile PBS containing the virus isolate at different doses (Supplemental Table 1). Mock-infected mice, injected intranasally with 50 µl sterile PBS, served as controls. The mice were monitored daily for clinical signs, weighed and euthanised by an overdose of pentobarbitone after reaching the clinical endpoint (maximum of 20% loss of body weight), or at the end of the experiment, at 4, 5, 6, 7 or 8 days post infection (dpi) (Supplemental Table 1). Mice were dissected immediately after death and the brain and lungs collected and sampled for downstream processing.

### Tissue processing

From all mice, the left lung was fixed in 10% neutral buffered formalin (NBF) and the right lung was frozen at −80 °C for RT-qPCR. In cohorts 1.3, 1.4 and 2.1, a sample from the frontal cortex was collected and frozen at −80 °C for RT-qPCR. The remaining brain of animals from cohort 1.3 as well as the whole brains in cohorts 1.1, 1.2 and 1.4 were fixed in 10% NBF. For all mice of cohorts 2.1-2.3, a slice of brain (approximately 1 mm thickness) was prepared by coronal sections at the level of the hippocampus and the thalamus (one section rostral, the second caudal to the pituitary ring) from which a sample (2-3 mm^3^) from the thalamic region was cut out and fixed in 5% glutaraldehyde (GA) (buffered in 0.2 M cacodylic acid buffer, pH 7.3) for transmission electron microscopy (TEM); for mice of cohorts 2.1 and 2.3, further samples were collected for bulk transcriptomic (thalamus), bulk metabolomic and bulk lipidomic (hypothalamus) and bulk proteomic (hippocampus and cerebral cortex) (Supplemental Figure S1) and frozen at −80 °C. The remaining brain tissue was fixed in 10% NBF. Information on animal cohorts and analyses undertaken is provided in Supplemental Table S1.

### Histological and immunohistological examinations

All NBF-fixed tissue samples were transferred to 70% ethanol after 24-48 h fixation, until further processing. The brain was trimmed by one sagittal or several coronal sections (cohorts 1.1-1.4), or coronal sections (cohorts 2.1-2.3). These as well as the left lungs were then routinely paraffin wax embedded. Consecutive sections (3-4 µm) were prepared and stained with haematoxylin and eosin (HE) or subjected to immunohistochemistry (IHC). All lungs and brains were stained for SARS-CoV-2 nucleocapsid protein (NP), as previously described [19]. In addition, brains from cohorts 2.1-2.3 were selectively stained to identify infiltrating leukocytes, i.e. for Iba1 (monocytes/macrophages, microglia), CD3 (T cells), CD4 (helper T cells), CD8 (cytotoxic T cells), and Ly6G (neutrophilic leukocytes), and apoptotic cells (cleaved caspase 3), as previously described [19, 42, 43]. These stains served to confirm that this set of experimental infections resulted in the same changes as those previously described by our group and in human patients [10, 19]. The brains of cohorts 2.1 and 2.3 were also stained for the expression of blood vessel/neurovascular unit (NVU) markers, i.e. platelet endothelial cell adhesion molecule 1 (PECAM-1/CD31; endothelial cells), claudin 5 (tight junction protein), platelet-derived growth factor receptor beta (PDGFR-β) and alpha smooth muscle actin (α-SMA; leiomyocytes, pericytes), and aquaporin 4 (astrocytic endfeet; AEF) [44–49]. Briefly, after deparaffination and antigen retrieval, sections were incubated with the primary antibodies diluted in Dilution buffer (Agilent Dako, Santa Clara, California, USA) at different temperatures and lengths of time. Subsequently, endogenous peroxidase was blocked using the Peroxidase Blocking Solution (Agilent Dako) for 10 min at room temperature (RT), followed by incubation with the appropriate secondary antibody and detection system and counterstaining with haematoxylin. Detailed information is provided in Supplemental Table S2. The following positive controls were used: a mouse lung previously tested positive for SARS-CoV-2 NP, and a mouse spleen for the leukocyte markers and cleaved caspase 3; for all other markers, the brain parenchyma served as internal positive controls. Negative control sections were incubated without the primary antibody in the diluent; lungs from uninfected control mice served as negative controls for SARS-CoV-2 NP.

The HE stained brain sections, together with the sections immunostained for SARS-CoV-2 NP were examined blindly and, besides a general histological examination, specifically assessed for the presence of (peri)vascular leukocyte infiltrates and for viral antigen expression. In the CNS, the “perivascular space” is a defined structural entity [50]. Therefore, in the present study, we speak about a “perivascular infiltrate” when leukocytes are located between the external layer of a vessel wall and the AEF. We apply the term “vascular infiltrate” when leukocytes are located within the vessel wall (which we and others have so far also referred to as “vasculitis” [19, 28, 31]). However, as it is often difficult to unequivocally distinguish between leukocytes located within the vascular wall and in the perivascular space in light microscopical specimens (in contrast to thin sections examined by TEM), we speak about (peri)vascular infiltrates in particular when referring to HE stained sections.

In order to allow a basic alignment of the extent of viral antigen expression and other parameters, the IHC-determined NP expression and distribution pattern was semi-quantitatively graded as a) negative (no immunostaining); b) scattered (variable number of individual positive neurons); c) clustered (variable numbers of groups of positive neurons); d) widespread (large/coalescing patches of positive neurons). The expression pattern is illustrated in Supplemental Figure S2.

### Transmission electron microscopy (TEM) including immune-EM

GA-fixed brain specimens from cohorts 2.1-2.3 were routinely embedded in epoxy resin. Semithin sections (0.95 µm) were prepared from the epoxy blocks and stained with toluidine blue to select the areas of interest (i.e. areas including blood vessels with or without (peri)vascular infiltrates) for the ultrathin sections (95 nm). The latter were prepared and contrasted with lead citrate and uranyl acetate. The ultrathin sections were viewed with a Philips CM10 microscope and digital images taken with a Gatan Orius SC1000 digital camera (Gatan Microscopical Suite, Digital Micrograph, California, USA). Detailed information on the workflow for the ultrastructural analysis is provided in the Supplemental Material (Material and Methods).

For immune-EM, ultrathin sections were prepared from the brains of cohorts 2.1 and 2.3 and incubated with citrate buffer (pH 6.0; Agilent Dako) at 60 °C for 7 h, followed by a 30 min incubation at RT in Antibody Diluent (Agilent Dako) and an overnight incubation at 4 °C with the rabbit anti-SARS-CoV-2 NP antibody used for IHC at a dilution of 1:1,000. Sections were subsequently washed with Wash Buffer 10x (Agilent Dako) and incubated with the secondary antibody (12 nm Colloidal Gold AffiniPure™ Goat Anti-Rabbit IgG (H+L), Jackson ImmunoResearch Laboratories Inc., Ely, UK), diluted 1:30 in Antibody Diluent for 2 h at RT. The sections were contrasted and examined as described above. An infected hACE2 transfected HEK-293T cell pellet (Supplemental Material) served to establish the immune-EM protocol and to define the optimal primary antibody concentration (Supplemental Figure S3). Sections of the pellet and brains incubated without the primary antibody as well as the brains of cohort 2.3 served as negative controls.

### RNA extraction and quantitative RT-PCR for viral load

RNA was extracted as previously described [41]. SARS-CoV-2 viral loads were determined in the right lung and frontal cortex tissue samples of cohorts 1.3, 1.4 and 2.1 that had been stored at −80 °C, using the GoTaq® Probe 1-Step RT-qPCR System (Promega). For quantification of SARS-COV-2 the nCOV_N1 primer/probe mix from the SARS-CoV-2 (2019-nCoV) CDC qPCR Probe Assay (IDT) and murine 18S primers were utilised as described previously [41]. The R packages used for the analysis are listed in Supplemental Table S3A.

### Multiomics analyses

Brain samples were collected and processed for bulk transcriptomic (thalamus), untargeted bulk proteomic (cerebral cortex and hippocampus), metabolomic (hypothalamus), and lipidomic (hypothalamus) analysis (Supplemental Figure S1). The code used for the analyses is available on request. The list of all used R packages is available in Supplemental Table S3B-E.

### Transcriptomic

Illumina RNA sequencing was undertaken by Novogene (Novogene (UK) Company Limited, Cambridge, UK) on RNA extracted from the coronal brain slice samples of cohorts 2.1 and 2.3, using polyA enrichment and the NovaSeq X Plus (Illumina®, San Diego, California, USA).

Sequencing reads were trimmed with fastp (v0.24.0, [51]). Trimmed paired end sequencing reads were inputted into salmon (v1.10.3, [52]) using the −l A –validateMappings –SeqBias –gcBias parameters with the “Mus_musculus.GRCm39.cdna.all.fa” reference sequence from ENSEMBL [53]. Quant files generated with salmon were imported into RStudio (RStudio: Integrated Development Environment for R, version 4.4.4, Posit Software, PBC, Boston, Massachusetts, USA) using tximport to infer gene expression (v1.32.0, [54]). Downstream analysis was conducted using edgeR utilising the filterByExpr() and calcNormFactors() functions, where differentially expressed genes were determined using the exactTest() function (v4.2.1, [55]). PCAtools (v2.16.0, [56]) was used to visualise transcriptional profiles, and functional enrichment analysis was performed by Gene Set Enrichment Analysis (GSEA)) with Gene Ontology (GO) [57, 58], in the clusterProfiler package with gseGO() (v4.12.6; [59–62]), with a Benjamini-Hochberg false discovery rate (FDR) threshold of 0.05. Details on the downstream analysis, including the packages used, are provided in the supplemental material (Supplemental Table S3B, Material and Methods).

### Proteomic

Brain tissue samples were homogenised by bead-beating (5 bursts of 15 sec; Minilys, Bertin Technologies, Montigny-le-Bretonneux, France) in 100 mM HEPES (pH 8.0) with 1% SDS, 1% NP-40, and 1× HALT protease inhibitor cocktail, and heated to 95 °C for 10 min to achieve viral inactivation. Lysates were clarified by centrifugation at 15,000 *x g*, 4 °C, for 10 min, and the supernatants sequentially treated with 4 mM dithiothreitol (DTT) at 60 °C for 10 min for cysteine reduction, and 14 mM iodoacetamide (IAM) for 30 min in the dark at RT for cysteine alkylation. Excess IAM was quenched with 3 mM DTT (7 mM total final concentration). Protein concentration was determined by Pierce BCA Protein Assay (Thermo Scientific, Rockford, USA) following the manufacturer’s microplate protocol, and 25 µg of total protein processed following a SP3-based clean-up and digestion protocol described elsewhere [63].

Liquid chromatography–coupled ion mobility mass spectrometry (LC-IM-MS) was performed using an Evosep One (Evosep, Odense, Denmark) coupled to a timsTOF-HT mass spectrometer (Bruker Daltonics, Bremen, Germany). The tryptic digests (200 ng) were loaded onto Evotips according to the manufacturer’s recommendations, and peptides resolved using a 30 SPD gradient on a PepSep C18 15 cm x 150 µm, 1.5 µm column (Bruker Daltonics) heated to 40 °C. Eluting peptides were ionised using a CaptiveSpray ESI source set to 1,600 V. The timsTOF HT was operated in positive mode performing data-dependent acquisition with parallel accumulation-serial fragmentation (PASEF) [64]).

Spectral data were analysed using FragPipe (v23.1) with MSFragger (v4.3) [65] and IonQuant v1.11.11 [66] integrated tools. Database searching and label-free quantification were performed using the LFQ-MBR workflow with the following modifications: semi-specific trypsin (P1 = K/R unless P1′ = P) and up to one missed cleavage. Database search was performed against UniProt [67] *Mus musculus* reference proteome UP000000589 (one protein per gene, 21,864 entries, assembly/annotation from ENSEMBL GCA_000001635.9) supplemented with SARS-CoV-2 UniProt reference proteome UP000464024 (17 entries, assembly/annotation from ENA/EMBL GCA_009858895.3) and common contaminants. FDR was set to 0.01 at the PSM, peptide, and protein levels. Label-free protein quantification was performed using unique and razor peptides, with FDR-controlled matching between runs.

Downstream analysis was conducted using DEP2 (v0.5.28.2 [68, 69]), utilising limma [70] to compute differential abundance, while functional enrichment analysis was performed with GSEA, using GO [57, 58] with a Benjamini-Hochberg FDR threshold of 0.05. Visualisation of proteomic profiles was done with PCAtools (v2.22.1). Details on the downstream analysis, including the packages used, are provided in the supplemental material (Supplemental Table S3C, Material and Methods).

### Metabolomic and lipidomic

Homogenised samples were collected in MeOH:IPA (1:1) and centrifuged at 11,000 rpm for 10 min at 4 °C. From the resulting supernatant, 400 µL was aliquoted for metabolomics, and 375 µL for lipidomics, stored at −80 °C.

For the <u>metabolomic</u> analysis, the remaining pellet was re-extracted using MeOH:ACN:H₂O (2:2:1) containing 2.5 µM of 13C-labeled amino acid mix (Supelco) as internal standard. Samples were vortexed, incubated at −20 °C for 1 h, and centrifuged (12,000 rpm, 10 min, 4 °C). The resulting supernatant was pooled with the initial extract and dried under nitrogen flow. Before liquid chromatography mass spectrometry (LC-MS) analysis, dried extracts were reconstituted in 90% acetonitrile, vortexed, and centrifuged (12,000 rpm, 10 min, 4 °C). A volume of 100 µL of the clarified supernatant was transferred to Total Recovery Vials (Waters) for injection. Method blanks, QC standards, and pooled samples were prepared in parallel and processed identically for quality control purposes.

Polar metabolites were acquired via LC-MS and were analysed in an untargeted fashion, as previously described [71]. For LC-MS analysis, the MS1 and MS2 resolutions were of 60,000 and 7,500, respectively. Filtering parameters used for the data analysis were: Signa/noise >3, mzCloud or mzVault match >50 ppm mass error within +/- 5 ppm, match with in-house developed MS1_RT library within +/- 10 sec, chromatographic peak and MS2 spectra quality.

The downstream analysis was undertaken with TidyMass pipeline (v2.0.10, [72]), and following best practices [73]. Briefly, for the differential analysis the computation of the fold change was done by comparing the medians, and the p-value was computed using a Wilcoxon rank sum test/Mann-Whitney U test. The metabolite profiles were visualised with PCAtools and pheatmap (v1.0.13 [74]). Details on the downstream analysis, including the packages used, are provided as supplemental material (Supplemental Table S3D, Material and Methods).

For the <u>lipidomic analysis</u>, the supernatants of all samples were spiked with EquiSPLASH (Avanti) as internal standard, vortexed and dried under nitrogen flow. Before LC-MS analysis, dried extracts were reconstituted in MeOH:IPA (1:1), vortexed, and centrifuged (12,000 rpm, 10 min, 10 °C). A volume of 90 µL of the clarified supernatant was transferred to Total Recovery Vials (Waters) for injection. Method blanks, QC standards, and pooled samples were prepared in parallel and processed identically for quality control purposes.

Lipids were acquired by LC-MS as previously described [71]. The MS1 and MS2 resolutions were of 60,000 and 30,000, respectively. The data set was evaluated in an untargeted fashion with Compound Discoverer software (Thermo Scientific). The modular data analysis workflow includes spectra selection, retention times alignment, compound detection and grouping, gap filling, and background filtering. mzVault was used to score fragmentation patterns and assign MS2-based identities to the features. A filtering process was performed, leading to the manually annotated compound table where each feature is annotated with the highest level of confidence. Filtering parameters used were the following: Signa/noise >3, mzVault match >50, chromatographic peak and MS2 spectra quality. Quality controls were run on pooled QC samples and reference compound mixtures to determine technical accuracy and stability.

Downstream analysis was undertaken using the LipidSigR pipeline (v1.0.4, [75–77]), following best practices [73]. For the differential analysis, the fold change was computed as the ratio of the means and the p-value was calculated using a Wilcoxon test. In addition to the statistical testing for individual lipid species, differential analyses were conducted for two lipid characteristics, namely: class and function. The lipid profiles were visualised with PCAtools and pheatmap. Details on the downstream analysis, including the packages used, are provided in the supplemental material (Supplemental Table S3E, Material and Methods).

## Results

### SARS-CoV-2 spread to the brain after intranasal challenge is independent of infection dose

After intranasal challenge of K18-hACE2 mice with SARS-CoV-2, viral RNA can be detected in lung and brain as soon as 1 and 2 dpi respectively, mostly with peak levels in the lungs at 3 dpi, whereas viral RNA in the brain increases to high levels from 4 to 7 dpi [18, 20, 24, 28, 31, 78]. Viral protein expression follows soon after RNA [29, 33]. In the present study we quantified SARS-CoV-2 RNA (N1 copy numbers normalised to 18S rRNA) in lungs and brain at 5, 6 or 7 dpi. Regardless of the inoculation dose, i.e. 10^3^ plaque-forming units (PFU)/mouse (moderate dose; cohort 1.3), or 10^2^ PFU/mouse (low dose; cohort 2.1) of SARS-CoV-2 Delta, we detected the virus at high copy numbers in the brain (Supplemental Figure S4) which confirmed that viral spread to the brain is independent of the inoculation dose. Interestingly, viral RNA levels in the lungs were consistently lower in mice that received the low dose than in mice that received the higher dose, whereas brain RNA levels were higher (Supplemental Figure S4). In both mouse cohorts, brain infection was associated with consistent, often widespread neuronal infection, as confirmed by viral NP expression (Supplemental Figure S2C; Supplemental Table S4).

### (Peri)vascular infiltrates occur with widespread neuronal SARS-CoV-2 infection of the brain

In K18-hACE2 mice, SARS-CoV-2 brain infection is often accompanied by a mild (meningo)encephalitis, partly with evidence of vascular infiltrates/vasculitis [18–20, 23, 27–29, 31]. To delve deeper into the associated inflammatory response, we re-examined blindly the brains of 49 intranasally-challenged mice, culled at 4-8 dpi, from several independent studies of our group (Supplemental Table S1).

Viral NP expression was detected in the brain of 85.7% (42/49) of infected mice, across all time points. At 4 dpi with 10^4^ PFU/mouse (cohort 1.2; Liverpool strain), viral antigen was detected in 4/5 brains, in clusters of infected neurons, with evidence of (peri)vascular infiltrates in one brain. Two mice (animals 1.3.1 and 1.3.2, infected with 10^3^ PFU/mouse SARS-CoV-2 Delta) had to be euthanised at 5 dpi; these showed widespread neuronal infection, accompanied by a perivascular inflammatory infiltrate in one. Almost all mice (23/26) culled at 6 dpi (infected with SARS-CoV-2 Liverpool or Delta isolates at 10^2^, 10^3^ or 10^4^ PFU/mouse; Supplemental Table S1), harboured viral antigen, in the majority (17/23) with widespread neuronal infection and generally associated with (peri)vascular infiltrates (16/17). With the “clustered” pattern of viral antigen expression (4/26), (peri)vascular infiltrates were seen in half of the cases (2/4). As inflammatory infiltrates were observed in areas of extensive SARS-CoV-2 antigen expression in neurons (Supplemental Figure S2C), they were lacking in brains with the “scattered” pattern (2 mice) or no viral antigen expression (3 mice). The findings were similar in mice culled at 7 dpi (cohorts 1.3, 2.1, and 2.2) and 8 dpi (cohort 2.2). Most brains showed the “widespread” pattern of viral NP expression (13/16), with (peri)vascular infiltrates in all but one of the positive cases. Individual animal data are provided in Supplemental Table S4.

### SARS-CoV-2 Delta infection of the brain does not target or lead to structural changes in the brain vasculature

#### (Peri)vascular infiltrates are not associated with vessel damage

We subsequently investigated the vessels in infected brains for interaction with leukocytes and pathological changes of the walls and, specifically, the endothelium. In line with our previous work [10, 19], we confirmed by IHC on selected brains of Delta infected mice (cohorts 2.1 and 2.2) that the (peri)vascular infiltrates were composed of monocytes/macrophages, T cells (CD4+, CD8+) and a few neutrophilic leukocytes. T cells and neutrophils were found to migrate further from the perivascular space into the parenchyma, however, they seemed not to target any specific cells or structures (Supplemental Figure S5).

To further this, the brains from cohorts 2.1 and 2.3 (Delta and mock-infected respectively) were IHC stained for key components of endothelium, vascular wall, and the vessel associated components of the NVU, namely PECAM-1 (endothelial cells), claudin 5 (tight junction protein), PDGFR-β and α-SMA (leiomyocytes, pericytes), and aquaporin 4 (AEF). We did not identify any differences in expression patterns and extent of these markers in vessels with and without inflammatory infiltrates in the infected brains (Figure 1), suggesting the vascular walls were intact. Staining for aquaporin 4 showed that, when present, the (peri)vascular infiltrates were mainly contained within the perivascular space, as these were delineated by a rim of AEF (Figure 1E), cell extensions that represent the outer limit of the perivascular space in the CNS [79]. Staining for PECAM-1, α-SMA, and PDGFR-β suggested the occasional presence of leukocytes within the vessel wall (Figure 1A, C, D).

**Figure 1.**
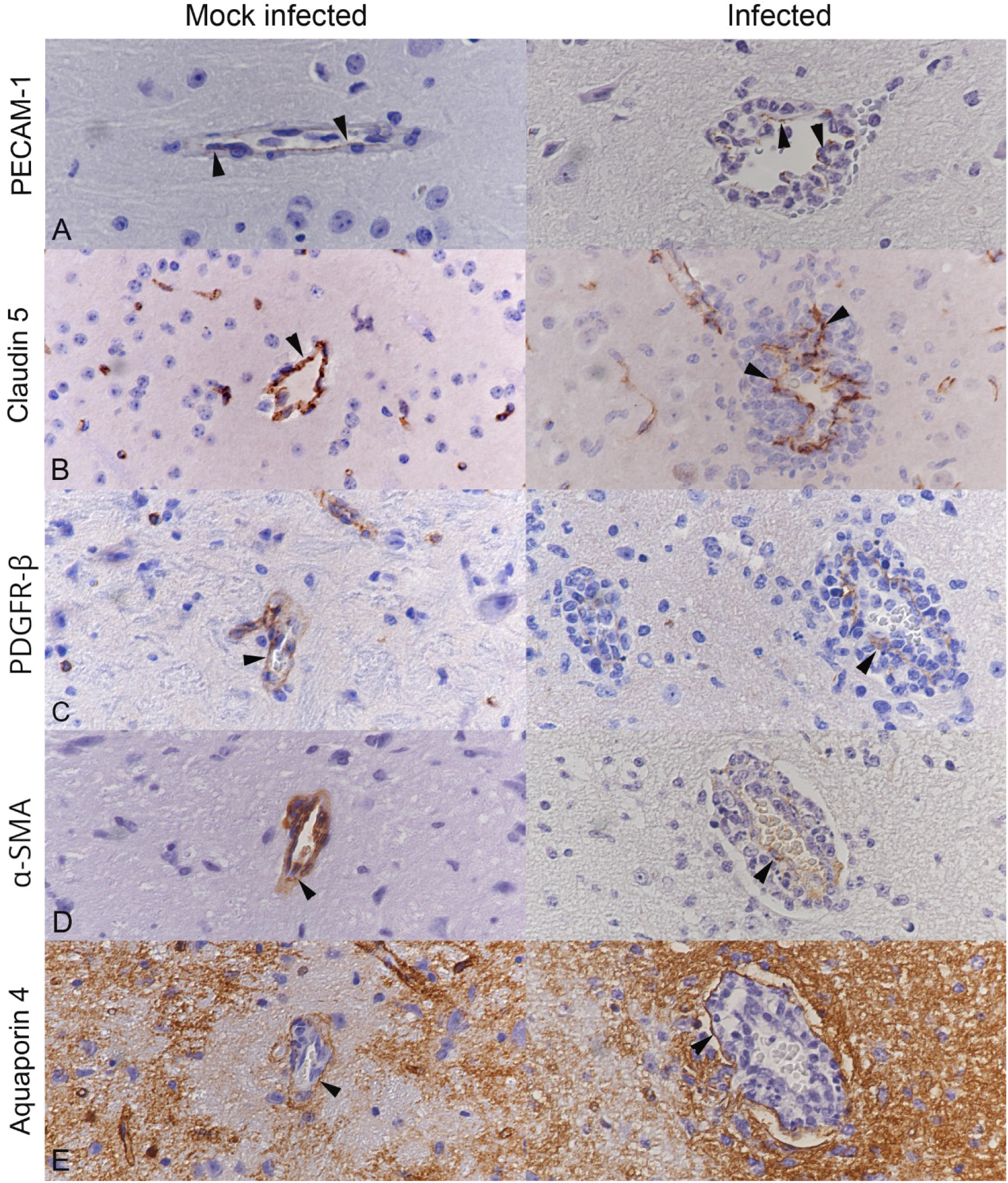
Structural key components of the blood-brain barrier/neurovascular unit are not damaged with (peri)vascular leukocyte infiltration. Brainstem of mock-infected mice (left column; cohort 2.3) or mice infected with SARS-CoV-2 Delta at 10^2^ PFU/mouse and euthanised at 7 dpi (right column; cohort 2.1). A) Endothelial cells (PECAM-1+; arrowheads) form a continuous layer (animals 2.1.4 and 2.3.2). B) Staining of the tight junction protein claudin 5 (arrowheads) confirms the integrity of the endothelial layer (animals 2.1.8 and 2.3.2). C, D) Staining of pericytes and leiomyocytes (C: PDGFR-β+; D: α-SMA+; arrowheads) highlights the integrity of the vascular wall (animals 2.1.4 and 2.3.1). E) Staining of astrocytic endfeet (aquaporin 4+) highlights that the infiltrating leukocytes accumulate in the perivascular space (animals 2.1.4 and 2.3.1). Immunohistochemistry, haematoxylin counterstain.

To gain more detail on the vascular morphology of infected brains, TEM was employed. The (peri)vascular infiltrates mainly affected post-capillary venules [50] where leukocytes, i.e. monocytes, neutrophilic leukocytes and lymphocytes, were observed in the vascular lumen and infiltrating the wall, with individual cells beneath the endothelial cell layer (Figure 2). The latter was continuous, with intact intercellular junctions between adjacent endothelial cells. Rare leukocytes were seen passing through the endothelial layer and the underlying basement membrane (Figure 2C, Supplemental Figure S6). Leukocytes were also located in the perivascular space, surrounded by a basement membrane and the AEF (Figure 2A-C, Supplemental Figure S6), or within the adjacent neuroparenchyma (Figure 2D, Supplemental Figure S6) where they were occasionally found to be apoptotic (Supplemental Figure S7A-C), in line with what was seen on light microscopy (cleaved caspase 3 IHC) and in our previous study [19].

**Figure 2.**
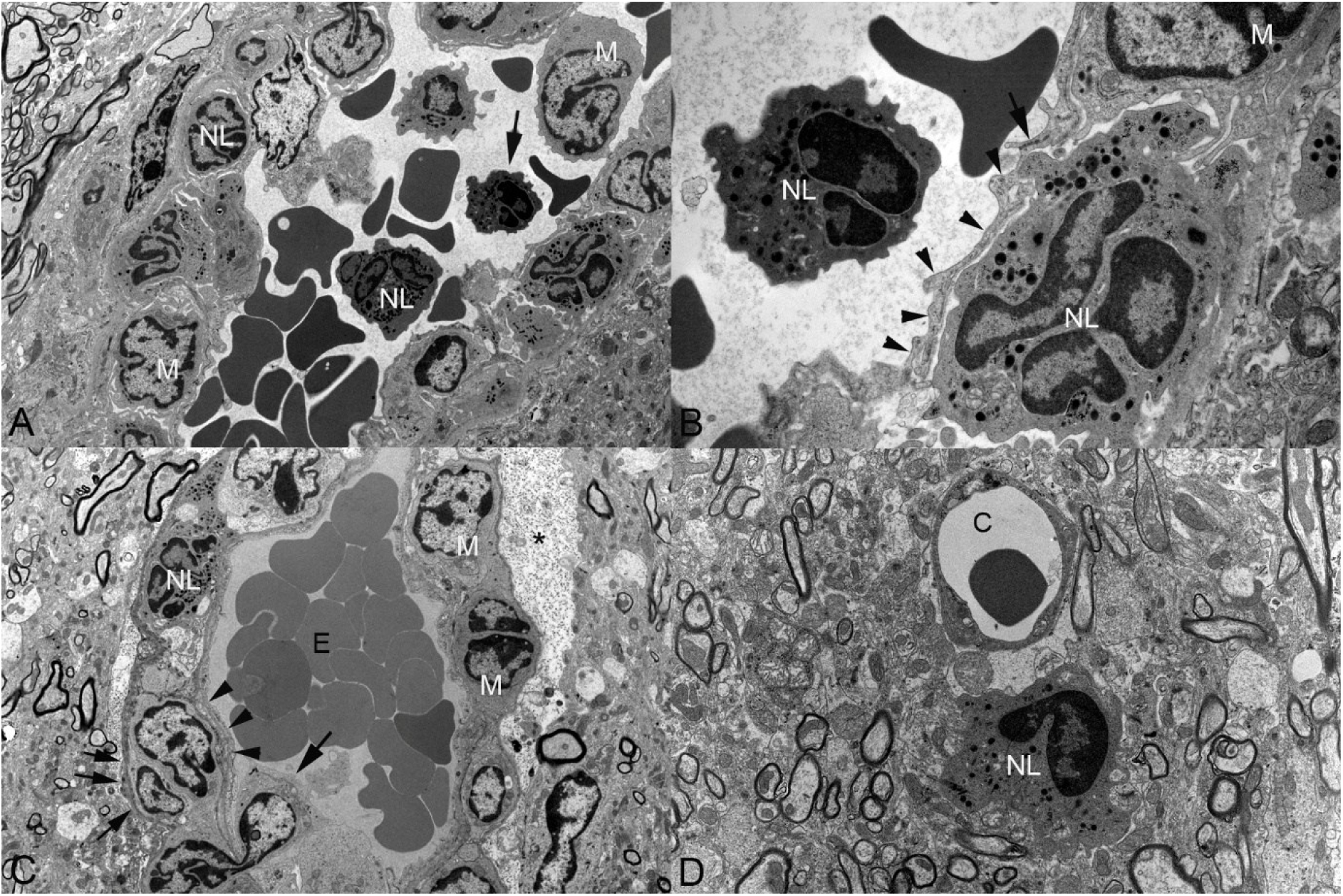
The (peri)vascular infiltrates are a consequence of leukocyte emigration without damage to the vascular wall. Brainstem, mice infected with SARS-CoV-2 Delta at 10^2^ PFU/mouse (cohort 2.1) or 10^3^ PFU/mouse (cohort 2.2) and euthanised at 6, 7, or 8 dpi. **A, B)** Animal 2.2.4. Post-capillary venule with accumulation of monocytes/macrophages (M) and neutrophilic leukocytes (NL) in the lumen and beneath the endothelial layer. The higher magnification (B; area indicated by the arrow in A) highlights a neutrophilic leukocyte and a monocyte/macrophage beneath the endothelial layer (arrowheads); the arrow points at an intact intercellular junction. **C)** Animal 2.1.4. Accumulation of monocytes/macrophages (M) and neutrophilic leukocytes (NL) in the perivascular space of a post-capillary venule. A leukocyte (likely a monocyte) is seen crossing the endothelial layer and its basement membrane (arrow). The vascular lumen is filled with erythrocytes (E). The astrocytic endfeet contain abundant finely granular electron-dense material (consistent with glycogen) (asterisk). Arrowheads: endothelial cells and associated basement membrane; small arrows: parenchymal basement membrane. **D)** Animal 2.1.1. Neutrophilic leukocyte (NL) in the neuroparenchyma immediately adjacent to a capillary (C). Transmission electron microscopy.

Regardless of the presence or absence of (peri)vascular leukocyte infiltrates, there was no evidence of morphological alterations of the vascular wall in infected brains (Figures 2-4; Supplemental Figures S6 and S7B, C).

#### The brain vasculature is spared from viral infection

Previous murine studies suggested that SARS-CoV-2 infection of the brain is limited to neurons [18, 19, 23, 26, 28–31, 33], however, some claimed that astrocytes, microglia, endothelial cells, and macrophages can also be infected [23, 28, 30]. To address this discrepancy, we employed light microscopy, TEM and immune-EM for viral NP. Strong staining for SARS-CoV-2 NP and ultrastructural elements consistent with virions (as seen in HEK293T cells; Supplemental Figure S3) were observed in neurons including many immediately adjacent to blood vessels (Figure 3); however, neither viral particles nor NP expression were detected in any structures/cells of the blood vessel wall or in surrounding non-neuronal cells, including any emigrating/infiltrating leukocytes (Figures 2 and 3; Supplemental Figures S6 and S7B, C).

**Figure 3.**
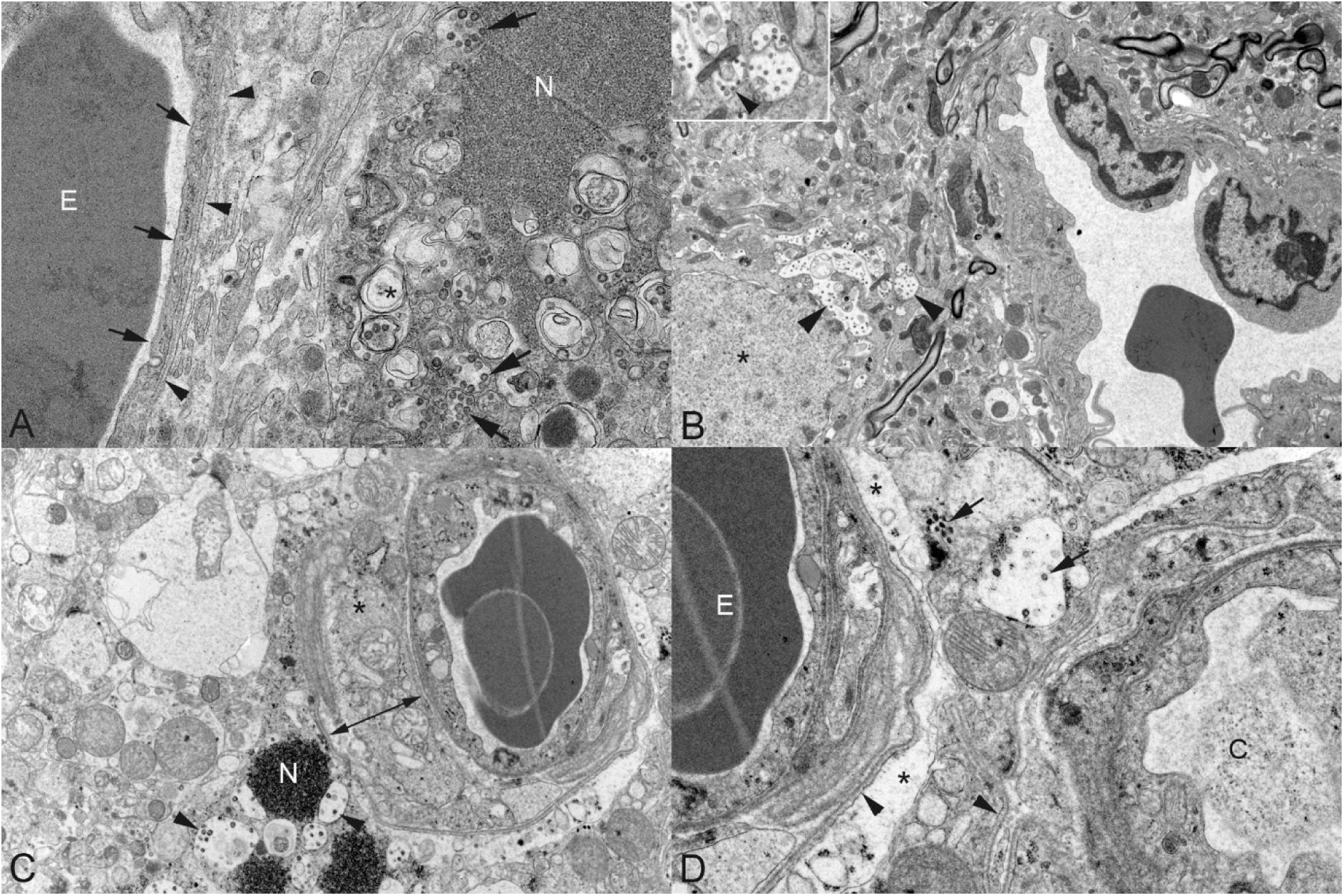
SARS-CoV-2 infection of the brain: no evidence that the virus targets the brain vasculature. Brainstem, mice infected with SARS-CoV-2 Delta at 10^2^ PFU/mouse (cohort 2.1) or 10^3^ PFU/mouse (cohort 2.2), euthanised at 6, 7, or 8 dpi. **A, B)** Transmission electron microscopy. **A)** Animal 2.1.3. Unaltered capillary (small arrows: endothelial layer, arrowheads: basement membrane; E: erythrocyte) with adjacent virus infected cell (morphology consistent with neuron) containing viral particles (arrows), replication-associated membranous structures (asterisk: double-membrane vesicles), and an electron dense aggregate consistent with cytoplasmic accumulation of nucleocapsid protein (NP) [25]. **B)** Animal 2.2.5. Unaltered (post-capillary) venule with adjacent virus infected neuron (asterisk: nucleus) with membrane-bound viral particles (arrowheads). Inset: higher magnification of viral particles in (distended) organelles (arrowhead). **C, D)** Immune-EM. Animal 2.1.5. Unaltered post-capillary venule adjacent to virus infected cells (morphology consistent with neuron). **C)** A monocyte/macrophage (asterisk) is present in the perivascular space (arrow with two tips). The infected neuron contains labelled membrane-bound viral particles (arrowheads) and strongly labelled cytoplasmic aggregates (consistent with accumulation of nucleocapsid protein (N)). **D)** Adjacent to the vessels (arrowheads: basement membrane) are astrocytic endfeet (asterisks) without evidence of viral particles or nucleocapsid protein deposition. An adjacent cell (morphology consistent with neuron) contains viral particles (arrows).

#### The AEF can exhibit subtle changes in infected brains

In the ultrastructural examination, we laid specific emphasis on the AEF, as an essential component of the NVU [80, 81]. In all examined brains, the majority of vessels were surrounded by more or less distinct AEF (Figure 4), reflecting the variable thickness of the sheet [82]; when prominent, the AEF were found to contain small amounts of finely granular electron-dense material (Figure 4) compatible with glycogen [83]. However, in some infected mice, a few vessels, both with and without (peri)vascular infiltrate, exhibited more prominent AEF containing an increased amount of the granular electron-dense material, i.e. glycogen (Figures 2C, 4). Progressive accumulation of glycogen in AEF has been interpreted as a change associated with BBB disruption in the peri-infarct brain parenchyma [83].

**Figure 4.**
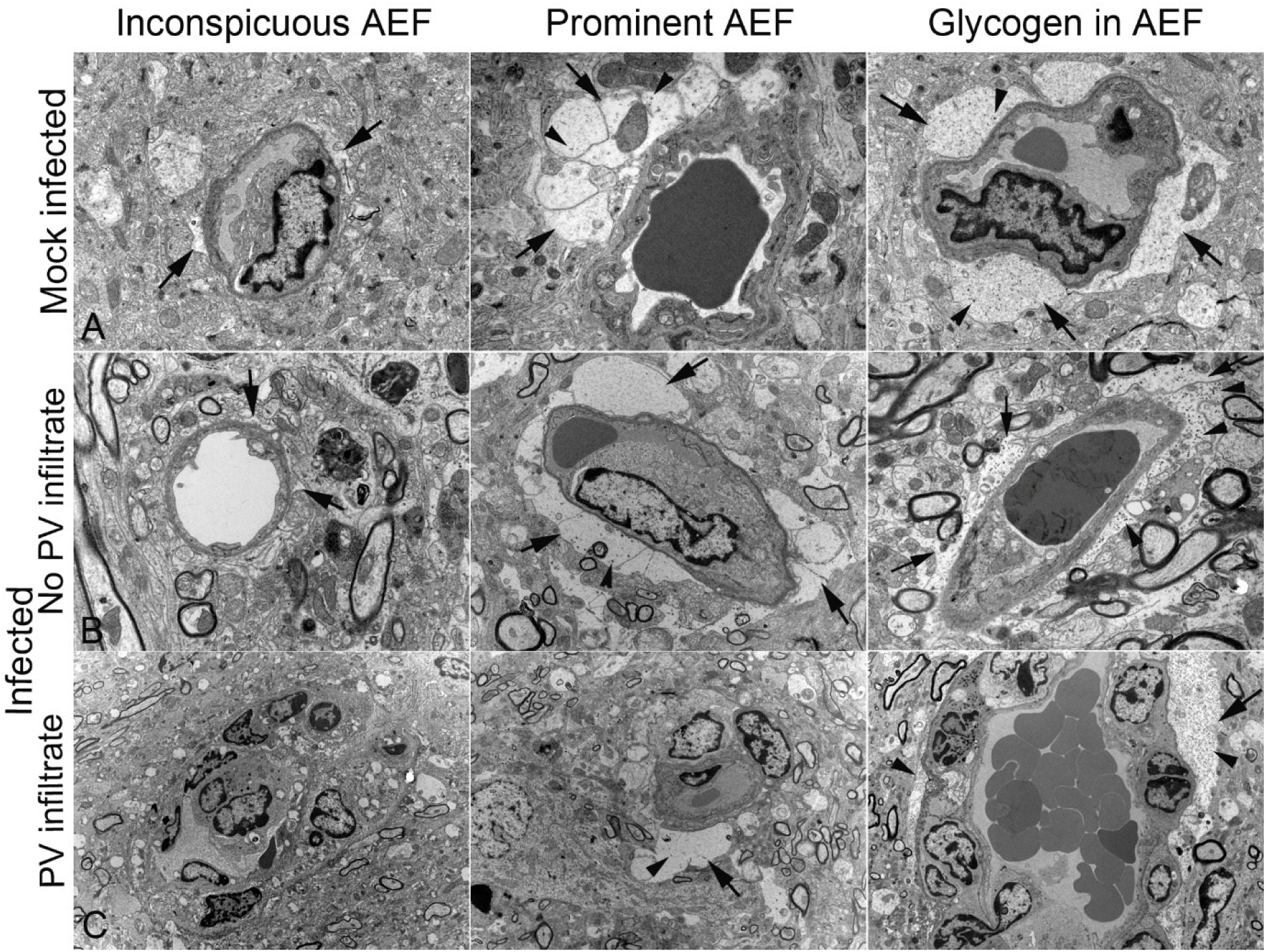
Morphological appearance of the astrocytic endfeet (AEF) of small calibre vessels. Brainstem. **A)** Mock-infected mice (animals 2.3.1 and 2.3.2). AEF (arrows) are either inconspicuous or prominent. Some of the latter contain finely granular electron-dense material compatible with glycogen granules (arrowheads). **B, C)** Mice infected with SARS-CoV-2 Delta at 10^2^ PFU/mouse (cohort 2.1; animals 2.1.1, 2.1.4, 2.1.8) or 10^3^ PFU/mouse (cohort 2.2; animal 2.2.5), euthanised at 6, 7, or 8 dpi. Regardless of the absence (B) or presence (C) of perivascular (PV) infiltrates, AEF (arrows) are either inconspicuous or prominent. They contain finely granular electron-dense material compatible with glycogen granules (B, C: right column; arrowheads) the amount of which appears higher than in the control brain. Transmission electron microscopy. The third picture on C is the same as Figure 2C.

### The transcriptome and proteome of SARS-CoV-2 Delta-infected brains reflect the *in situ* findings

To complement the *in situ* investigation, and to gain better understanding of the molecular changes in the infected brains, we performed a bulk transcriptomic and a bulk untargeted proteomic analysis on SARS-CoV-2 Delta infected and mock-infected control brains that had been examined in depth at light microscopical and ultrastructural level (cohorts 2.1 and 2.3; Supplemental Table S1), using brain samples collected adjacent to the TEM samples (Supplemental Figure S1). We aimed to 1) investigate the inflammatory environment associated with brain infection; 2) further characterise the leukocyte recruitment process; 3) determine the state of activation in recruited leukocytes; and 4) detect any changes that reflect or are associated with BBB dysfunction.

Despite exclusion of SARS-CoV-2 viral sequences and proteins from the analysis, we observed clear clustering of the Delta infected and mock infected cohorts (Figure 5A), consistent with infection-induced changes at host level. Gene products contributing to the principal component analysis (PCA) loadings highlighted inflammatory processes, in particular *Cxcl9*, *Cxcl10*, *Il1rn* (chemokines and a cytokine [84, 85]); *Lcn2* (encoding for the acute-phase protein like lipocalin 2, known to induce chemokine production and play a role in neuroinflammation [86, 87]); *Slfn4* (Schlafen 4; up-regulated during macrophage activation [88]); and *Ifi35* (interferon-induced protein 35), an interferon-stimulated gene involved in the regulation of type I interferon and cytokine production, with reported pro- or antiviral effects [89–91].

**Figure 5.**
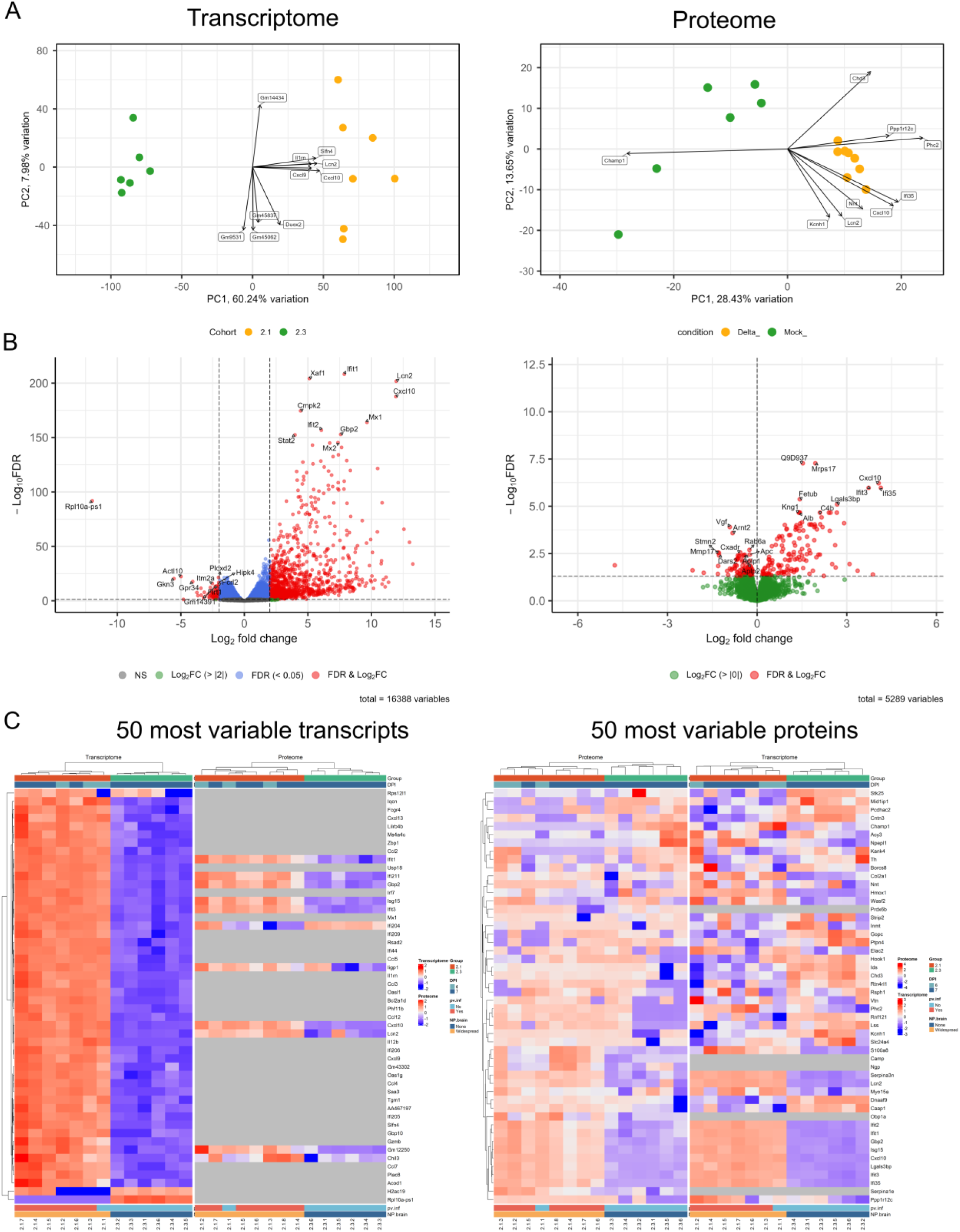
Results of the bulk transcriptomic and proteomic analyses in SARS-CoV-2 Delta-infected brains. Mice infected with SARS-CoV-2 Delta at 10^2^ PFU/mouse (cohort 2.1) and mock-infected mice (cohort 2.3). **A)** The Principal Component Analysis biplots show the clustering of the brains in their respective cohort, as well as the top loadings. **B)** Volcano plots showing the distribution of the investigated transcripts and proteins. **C)** Heatmaps showing the 50 most variable transcripts or proteins, in the transcriptome and proteome.

Differential analysis showed pronounced changes in the host transcriptome and proteome, with significant up-regulation of 1,103 genes and 141 proteins, and significant down-regulation of 80 genes and 112 proteins (Figure 5B). Of the significantly up-regulated gene products, 58 were shared between the transcriptome and proteome (Supplemental Figure S8).

Identification of the 50 most variable genes and proteins showed significant up-regulation of gene products related to inflammation and anti-viral response, such as chemokines, interleukins, interferon activated genes, and interferon-induced proteins (Figure 5C). Interestingly, while almost all of the most variable transcripts were significantly different between the two cohorts, this was the case for less than half of the most variable proteins.

Functional enrichment analysis, using GSEA with GO [57, 58], further confirmed this trend, with positive enrichment of pathways involved in response to virus and bacterium, inflammation, and immune response, whereas pathways associated with synapses/neurotransmitters, channel activity, and adult behaviour were negatively enriched in infected brains (Figure 6).

**Figure 6.**
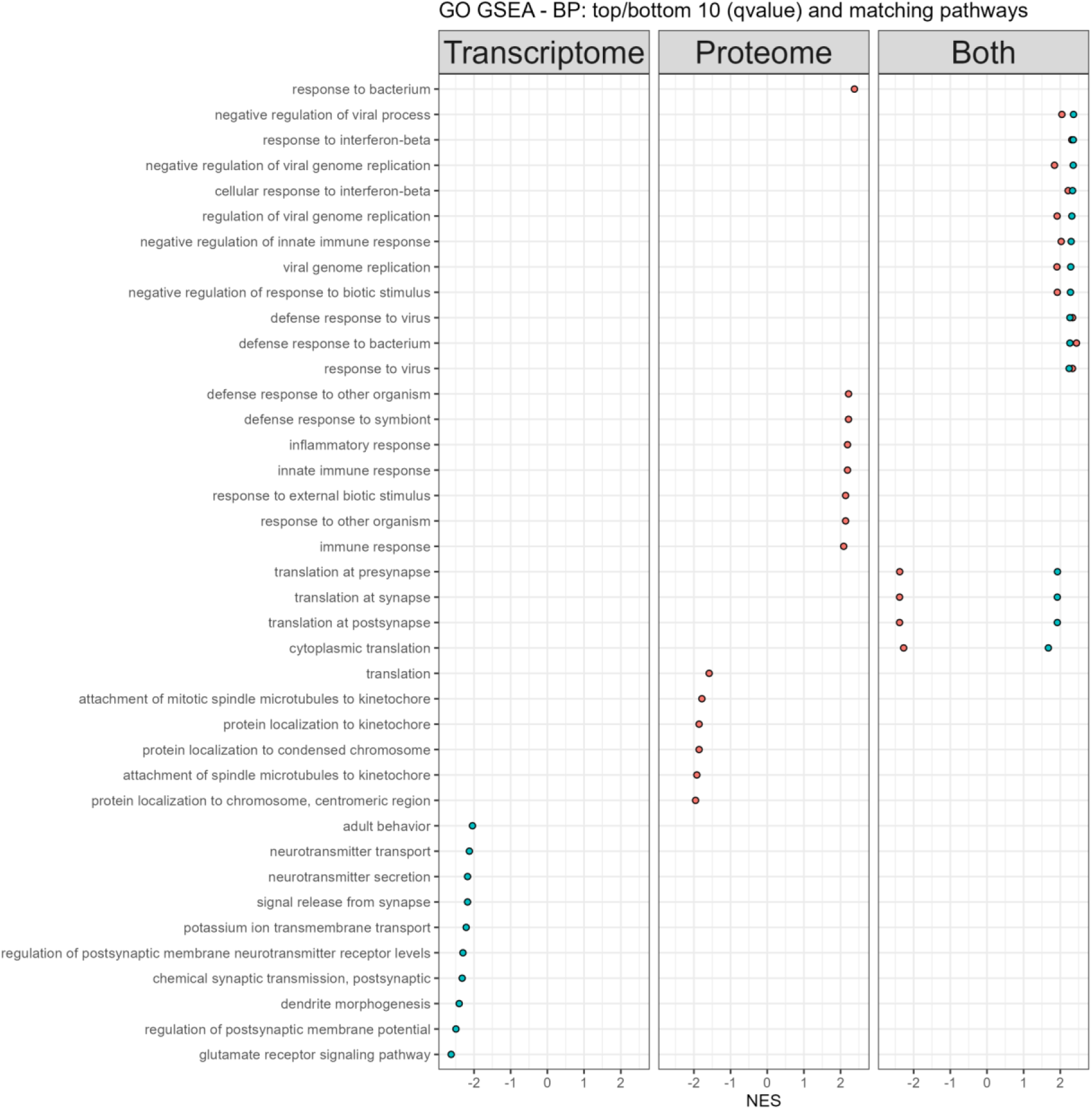
GO GSEA pathway enrichment analysis in SARS-CoV-2 Delta-infected brains. Mice infected with SARS-CoV-2 Delta at 10^2^ PFU/mouse (cohort 2.1) and mock-infected mice (cohort 2.3). Illustration of the top and bottom 10 enriched pathways (based on q-value) for the transcriptome and proteome, with matching pathways, sorted on the coalesced transcriptome and proteome normalised enrichment score (NES). BP: biological process, GO: Gene Ontology.

To understand the molecular mechanisms underpinning neuroinflammation and leukocyte recruitment/activation, and to determine potential effects on blood vessels and BBB function, we further investigated relevant sets of genes and pathways.

#### Inflammatory environment

The presence of an inflammatory environment in the infected brains was supported by positive enrichment of pathways associated with inflammatory response, interferon type I and II, cytokines, prostaglandins, and inflammasome (Figure 7A). Investigation of selected categories of inflammatory mediators (cytokines, eicosanoids, complement, vasoactive amines) confirmed this trend, with significant up-regulation of gene products from all four categories in the infected brains (Figure 7B). Two Suppressors of Cytokine Signalling (SOCS), *Socs1* and *Socs3,* were significantly up-regulated at transcript level, suggesting the simultaneous activation of inhibitory pathways [92].

**Figure 7.**
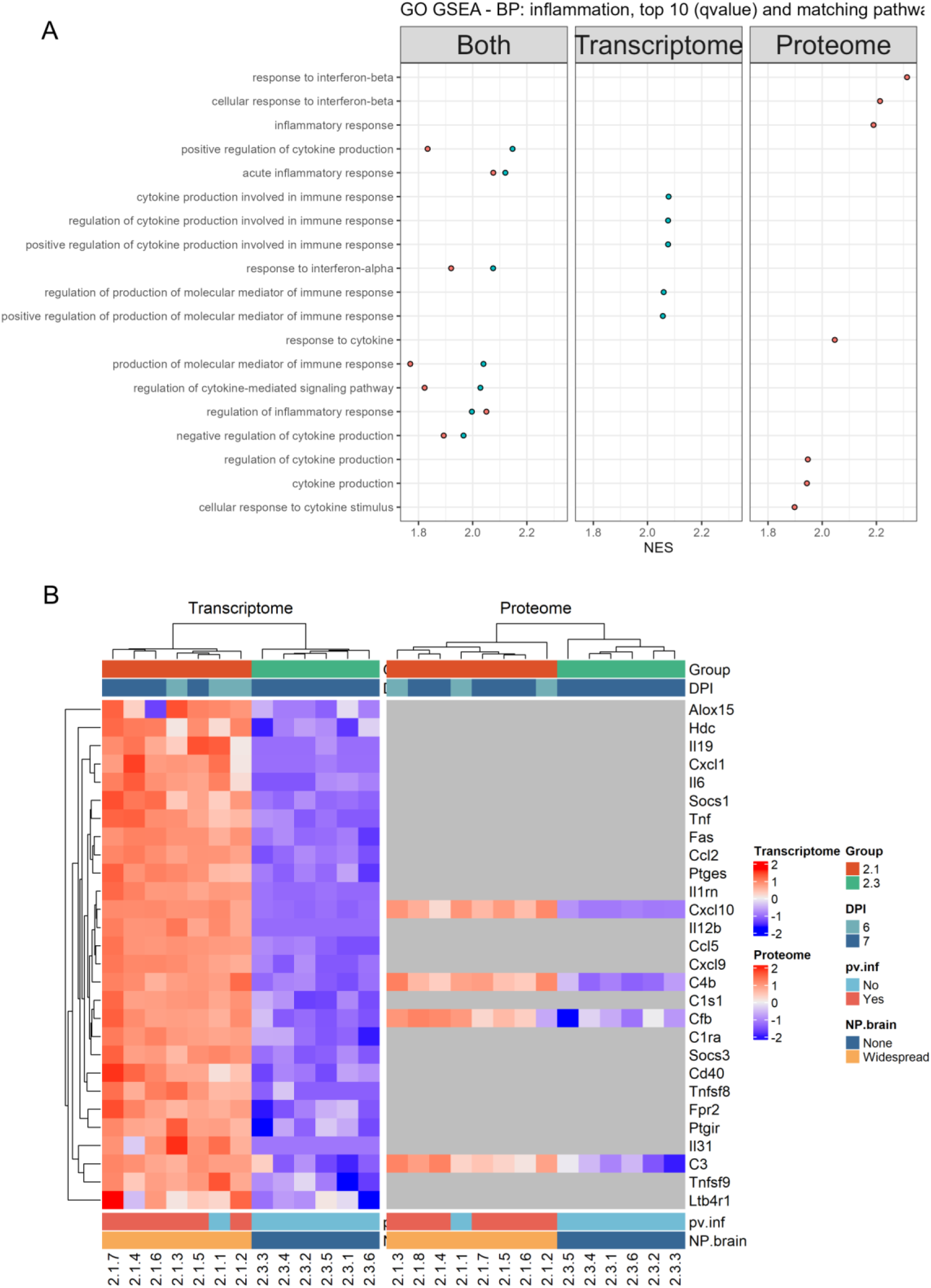
Inflammation: pathways and genes of interest. Mice infected with SARS-CoV-2 Delta at 10^2^ PFU/mouse (cohort 2.1) and mock-infected mice (cohort 2.3). **A)** Illustration of the top 10 enriched pathways (GO GSEA) (based on q-value) of the Biological Process (BP) ontology for the transcriptome and proteome, with matching pathways, sorted on the coalesced transcriptome and proteome normalised enrichment score (NES). **B)** Illustration of selected inflammation related genes from several categories/families of inflammatory mediators (cytokines – chemokines, tumour necrosis factor (TNF), interleukins – eicosanoids, complement, vasoactive amines, suppressor of cytokine signalling (SOCS)) (the top 5 gene products from each category/family based on transcripts log_2_FC were plotted).

#### Leukocyte emigration and activation

Pathways associated with leukocyte adhesion to the endothelium, migration and chemotaxis were enriched in infected brains, consistent with active recruitment of leukocytes into the brain and supporting our *in situ* findings (Supplemental Figure S9A). Several pathways related to leukocyte activity and the immune response were enriched (Figure 8A, Supplemental Figure S9B), indicating that the recruited leukocytes are part of a tissue response to virus infection. Further investigation revealed the enrichment of pathways associated with different leukocyte subtypes, such as lymphocytes (CD4+ and CD8+ T cells, B cells, natural killer cells), monocytes/macrophages, and granulocytes (including neutrophilic leukocytes) (all present in the (peri)vascular infiltrates observed in infected brains; Supplemental Figure S5); these were mostly associated with leukocyte activity, proliferation/differentiation, and migration/chemotaxis (Supplemental Figure S9). Enrichment of pathways involved in leukocyte mediated cytotoxicity, cell killing, degranulation, phagocytosis, respiratory burst and reactive oxygen species confirmed leukocyte activation (Supplemental Figure S9B) while the enrichment of several pathways associated with cell death (including lymphocyte (T/B) and neutrophilic leukocyte death) (Supplemental Figure S7D) aligns with the leukocyte apoptosis detected *in situ* (Supplemental Figure S7A-C). To further assess the interaction of leukocytes with the NVU, we investigated selected genes involved in leukocyte recruitment, emigration and activity, and found significant up-regulation of several gene products involved in different steps of this process [50, 93], such as capture (*Vcam1*), rolling (*Selp*, *Sele*), arrest and adhesion (*Icam1*, *Vcam1*), polarisation (*Icam1*), crawling against the blood flow (*Icam1*), migration across the endothelial barrier (*Cxcl10*, *Ccl2*, *Ccr7*), sojourn in the perivascular space (*Mmp19*), and migration across the parenchymal barrier (*Timp1*, *Ccl2*, *Tnfrsf1a*, *C3, Cfb*) in infected brains (Figure 8B). Significant up-regulation of genes involved in cell killing (*Gzma, Gzmb*, *Prf1*) and the generation (*Mpo*) of reactive intermediates at transcript level (Figure 8B) reflects leukocyte activity [94, 95].

**Figure 8.**
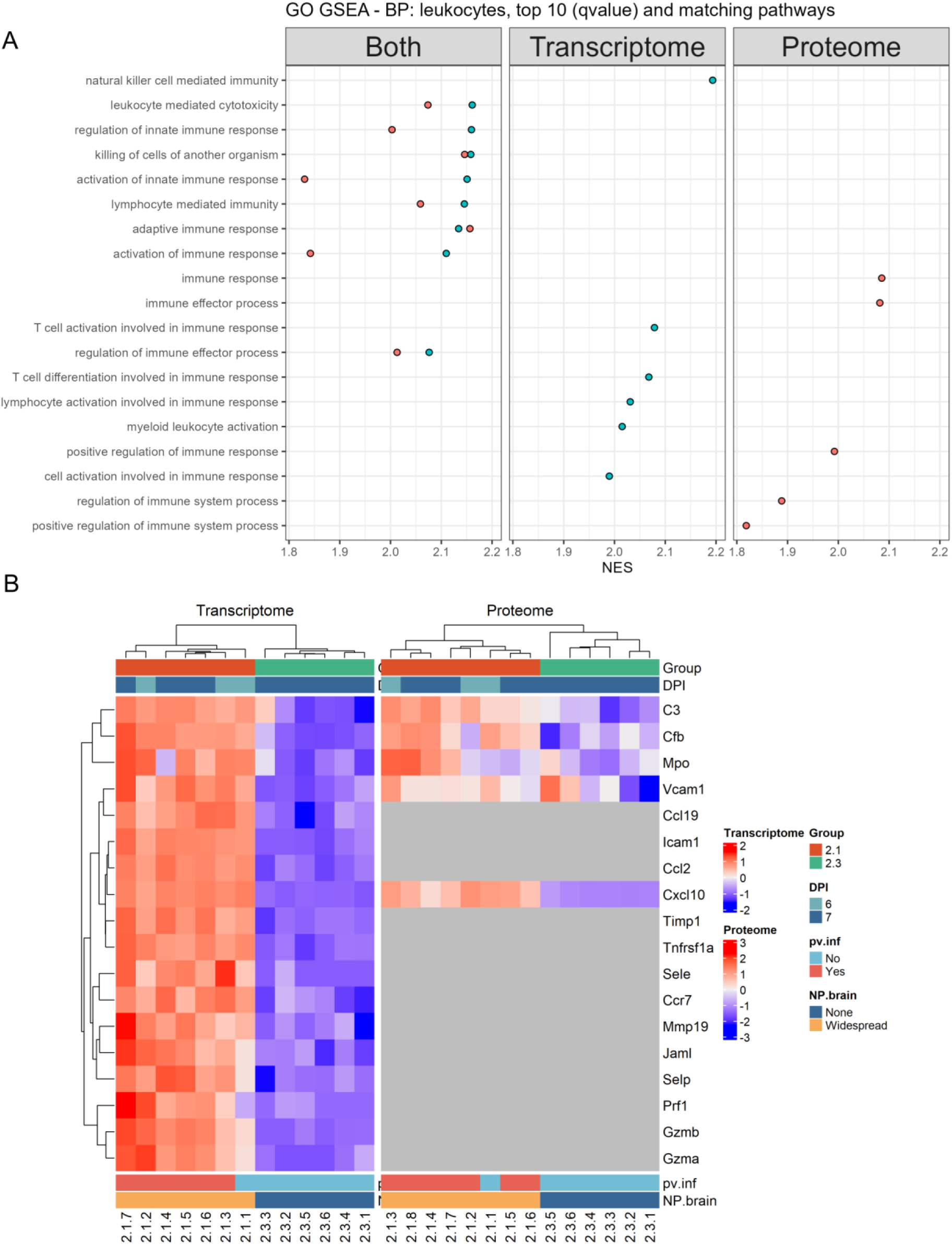
Leukocytes: pathways and genes of interest. Mice infected with SARS-CoV-2 Delta at 10^2^ PFU/mouse (cohort 2.1) and mock-infected mice (cohort 2.3). **A)** Illustration of the top 10 enriched pathways (GO GSEA) (based on q-value) of the Biological Process (BP) ontology for the transcriptome and proteome, with matching pathways, sorted on the coalesced transcriptome and proteome normalised enrichment score (NES). **B)** Illustration of the investigated genes of interest for leukocyte recruitment and activity which are significantly up-regulated in the infected brains.

#### Blood vessel, NVU, and BBB function

The morphological examination of the blood vessels did not reveal any structural alterations. However, positive enrichment of pathways associated with angiogenesis/vascular development, endothelial cell migration and apoptotic process, smooth muscle cell proliferation and apoptotic process, and tight junction assembly in infected brains was observed (Figure 9A). Further investigation of the BBB/NVU, assessing selected genes associated with its constitutive components (i.e. endothelial cells, pericytes, AEF, basement membrane), revealed differential expression of several transporters [81]: *Slc28a2*, *Slc1a5* and *Ager* were up-regulated, *Slc22a8* and *Tfrc* were down-regulated in infected brains. However, cell junctions (*Cldn1*, *Cldn3*, *Cldn5, Cldn12, Ocln, Tjp1, Tjp2, Tjp3, Cdh1, Cdh5, Pecam1)* were unaffected (Figure 9B). Investigating genes known to be associated with alterations of BBB function, we found angiopoietin 2 (*Angpt2*), involved in various vascular processes [96–98], to be significantly up-regulated in infected brains.

**Figure 9.**
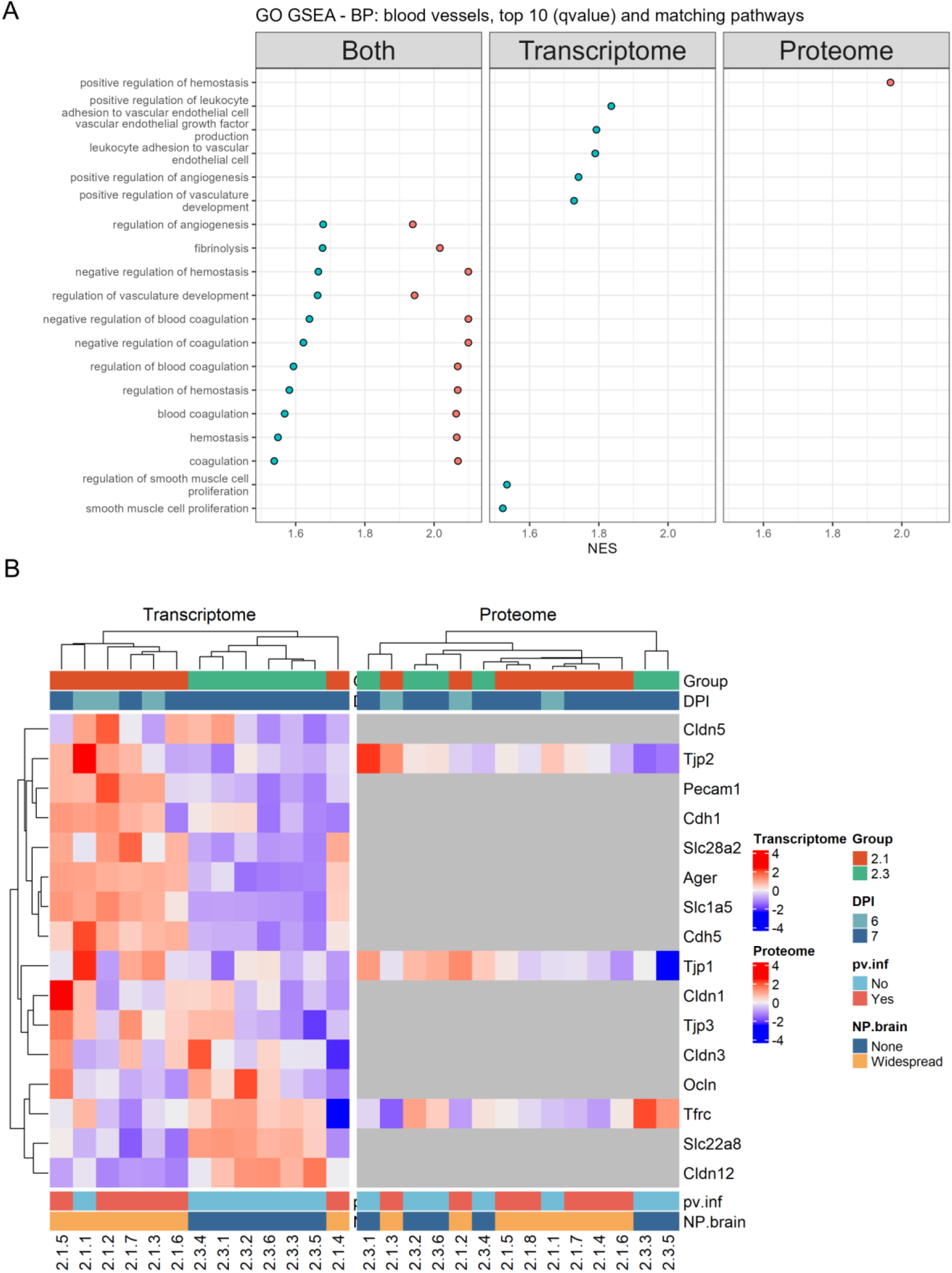
Blood vessels and blood-brain barrierneurovascular unit: pathways and genes of interest. Mice infected with SARS-CoV-2 Delta at 10^2^ PFU/mouse (cohort 2.1) and mock-infected mice (cohort 2.3). **A)** Illustration of the top 10 enriched pathways (GO GSEA) (based on q-value) of the Biological Process (BP) ontology for the transcriptome and proteome, with matching pathways, sorted on the coalesced transcriptome and proteome normalised enrichment score (NES). **B)** Illustration of gene products associated with tight junctions, as well as gene products of transporters which are significantly different.

To broaden our analysis, we ran our two datasets against published transcriptome and proteome data of the murine BBB [99–101] (Supplemental Figure S10). This revealed significant up-and down-regulation of additional gene products (Figure 10A, Supplemental Table S5); amongst these, *Aoc3* (endothelial adhesion protein involved in leukocyte binding and migration [102–104]), *Apod* (apolipoprotein associated with maintaining BBB integrity [105]), *Emp1* (a tight junction-associated protein [106]) and *Slc43a3* (purine-selective nucleobase transporter [107, 108]) were significantly up-regulated, while *Abcc6* (ATP-dependent transporter the exact function of which remains to be elucidated [109, 110]) and *Slc40a1* (ferroportin, iron-exporting protein that plays a role in iron entry into the CNS [111, 112]) were significantly down-regulated.

**Figure 10.**
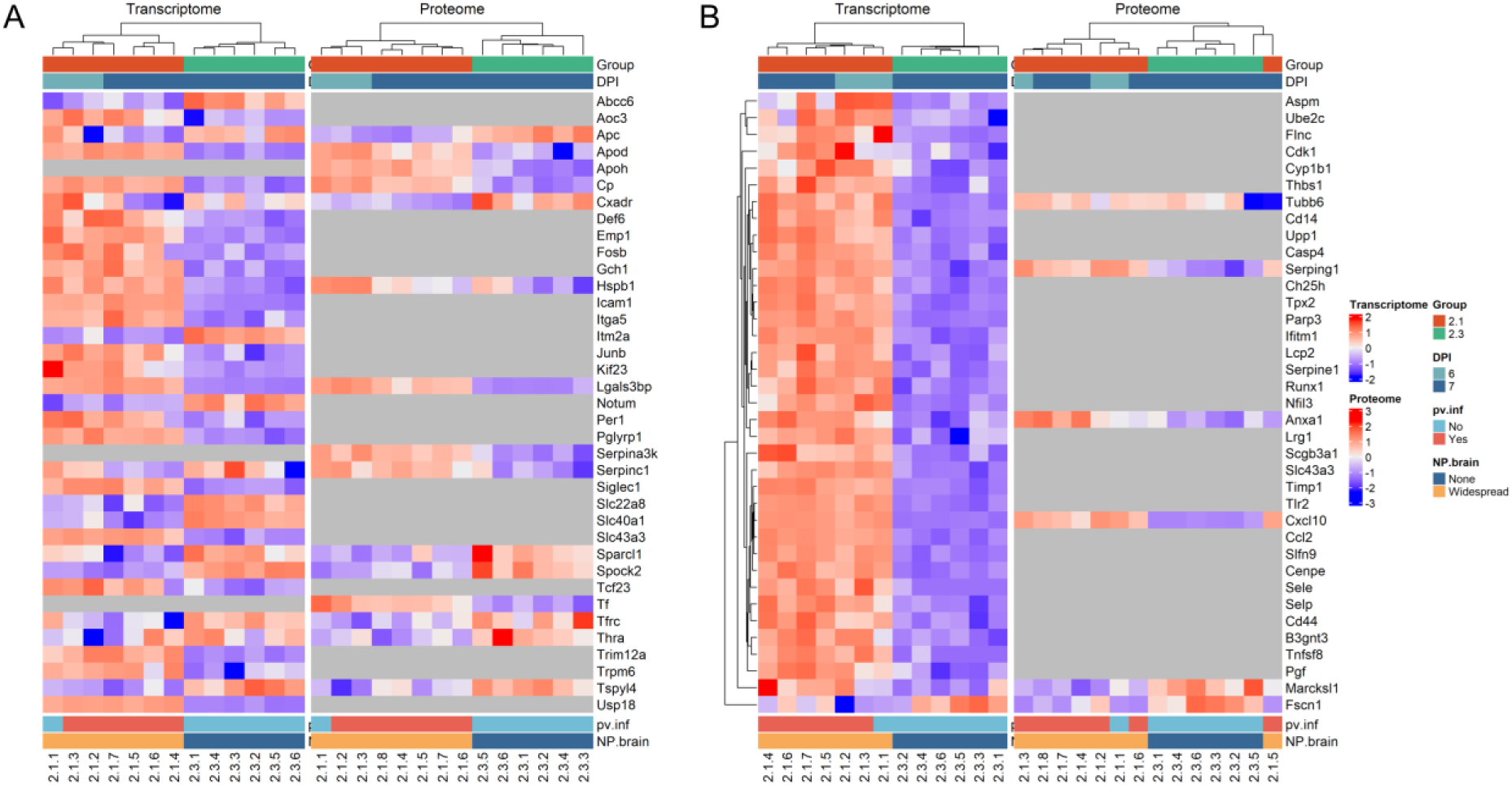
Blood vessels and blood-brain barrierneurovascular unit: comparison with published murine datasets. Mice infected with SARS-CoV-2 Delta at 10^2^ PFU/mouse (cohort 2.1) and mock-infected mice (cohort 2.3). **A)** Illustration of the significantly different transcripts and/or proteins after comparison with Tremblay et al., 2024 (proteome of the BBB); Daneman et al., 2010 (transcriptome of the BBB); and Munji et al., 2019 (transcriptome of BBB-enriched genes and transcriptome of tight junction proteins expressed in brain endothelial cells). **B)** Illustration of the significantly different transcripts and/or proteins after comparison with Munji et al., 2019, BBB dysfunction module.

In addition, we probed our data using the “BBB dysfunction module” [101]; here, we retrieved 130 of the 136 genes that compose this module from our datasets. Of these, 35 were significantly up-regulated (some of these were already highlighted as part of the neuroinflammation and leukocyte recruitment), and two (*Marcksl1* and *Fscn1*) were significantly down-regulated (Figure 10B).

These findings are in line with the *in situ* analysis that did neither detect structural alterations of the components comprising the BBB/NVU, nor morphological evidence of effective BBB leakage. Hence, the observed gene product alterations, affecting components of the BBB/NVU and a known marker of BBB alteration, could suggest minimal BBB dysfunction; the ultrastructural evidence of increased glycogen in the AEF could be the consequence of the latter.

### SARS-CoV-2 Delta infection with mild inflammatory response is associated with minimal metabolomic and lipidomic disturbance in the brain

The brain metabolome and lipidome of SARS-CoV-2 Delta infected mice showed minimal changes (Figures 11 and 12) in metabolites and lipids (fatty acyls) involved in oxidative stress, inflammation and immune/antiviral response (Figure 11C, D), all in line with the transcriptome and proteome results: Citraconic acid (a reported immunomodulator exerting anti-inflammatory, anti-oxidant, and anti-viral effects and acting via *Acod1* inhibition – a mitochondrial enzyme involved in the synthesis of itaconic acid [113, 114]) was significantly less abundant in the infected brains; interestingly though, *Acod1* was significantly up-regulated and itaconic acid (a key immunomodulator [113, 114]) was unchanged in the infected brains. This suggests activation of immunomodulator pathways in the latter. S-adenosylhomocysteine (an inhibitor of cell methyltransferases with roles in inflammation via NF-κB, also involved in SARS-CoV-2 replication and immune evasion [115–117]) was also significantly less abundant in the infected brains. The enrichment of pathways associated with reactive oxygen species and the significant up-regulation of *Mpo* and *Nos2* (nitric oxide synthase 2, inducible) transcripts are consistent with oxidative stress in the infected brains; interestingly, two known anti-oxidants, quinic acid [118] and uric acid (a derivate of purine metabolism [119]), were significantly elevated as well.

**Figure 11.**
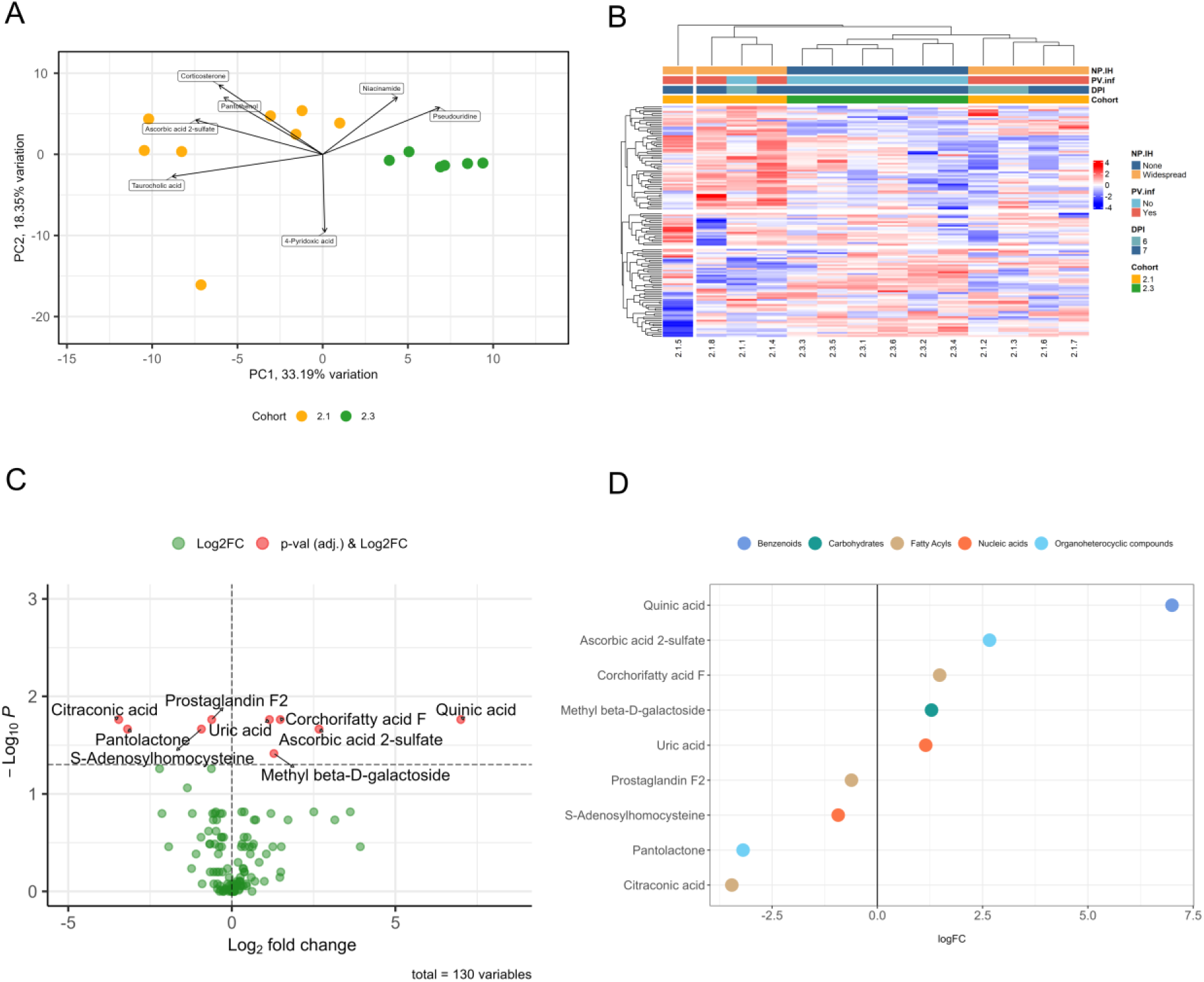
Results of the bulk metabolomic analysis in SARS-CoV-2 Delta-infected brains (hypothalamus). Mice infected with SARS-CoV-2 Delta at 10^2^ PFU/mouse (cohort 2.1) and mock-infected mice (cohort 2.3). **A)** The Principal Component Analysis biplot shows the clustering of the samples in their respective cohort, as well as the top loadings. **B)** Hierarchical clustering of the samples based on all 130 metabolites. **C)** Volcano plot showing significantly different metabolites between the two cohorts. **D)** The significantly different metabolites are shown with their respective super class.

**Figure 12.**
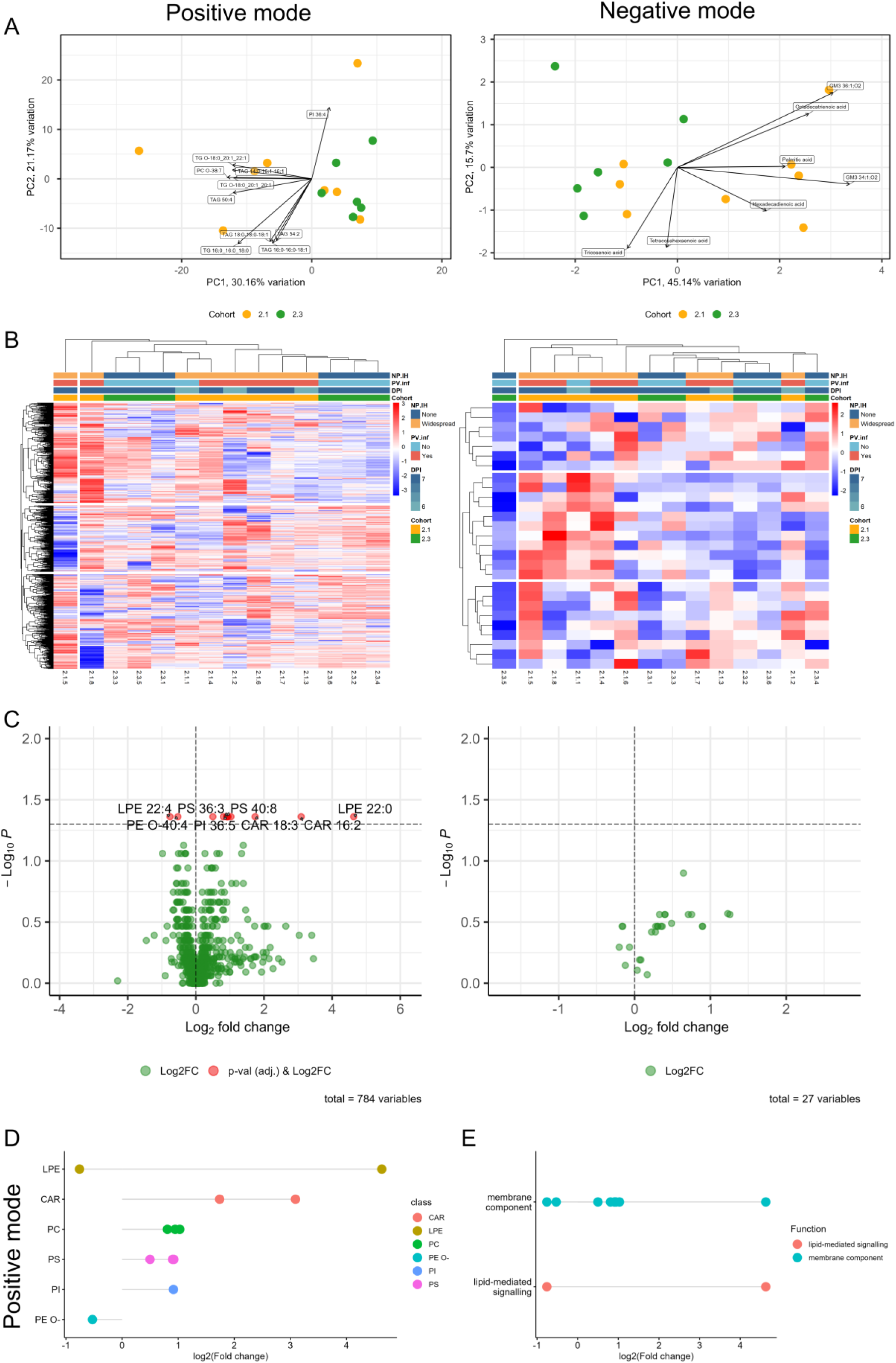
Results of the bulk lipidomic analysis (positive and negative modes) in SARS-CoV-2 Delta-infected brains (hypothalamus). Mice infected with SARS-CoV-2 Delta at 10^2^ PFU/mouse (cohort 2.1) and mock-infected mice (cohort 2.3). **A)** The Principal Component Analysis biplots show the clustering of the samples, as well as the top loadings. **B)** Hierarchical clustering of the samples based on all lipids for each mode. **C)** Volcano plot showing significantly different lipids between the two cohorts. **D)** The significantly different lipids are shown with their respective class. **E)** Function associated with the significantly different lipids.

Further differential analysis of individual lipid species revealed significant differences in only a few additional lipids, namely: CAR 16:2, CAR 18:3, LPE 22:0, LPE 22:4, PC18:1-18:2, PC 36:5, PC38:6, PE O-40:4, PI 36:5, PS 36:3, PS 40:8, PS 44:12. These lipids represent several lipid classes; some are associated with membrane components and lipid-mediated signalling, for others the function is unknown (Figure 12).

## Discussion

The present study focused on the response of the brain to extensive but exclusively neuronal infection with SARS-CoV-2, as a virus without a major neuropathic effect. Intranasal challenge of K18-hACE-2 mice with SARS-CoV-2 VOC proceeds to widespread infection of the brain in a large proportion of infected animals, most likely via the olfactory route, and takes places within the first 4 days post challenge [18–20, 28, 33, 120]. Infection is often associated with a mild vessel-centred inflammatory response (i.e. (peri)vascular leukocyte infiltrates) and microgliosis [18–20, 28, 29, 31]. Here we showed that also a very low dose of SARS-CoV-2 Delta (10^2^ PFU) leads to widespread neuronal infection and mild encephalitis in areas of intense neuronal infection by 6 to 7 dpi after intranasal challenge.

In-depth *in situ* investigation showed that, different from early assumptions made in fatal COVID-19 cases (reviewed in [7, 121]), the (peri)vascular leukocyte infiltration is not a consequence of the virus targeting endothelial cells or other components of the vascular wall. Indeed, our combined morphological and multiomics investigations clearly show that in K18-hACE2 mice it is a consequence of recruitment of leukocytes from the blood into the brain in areas of intense neuronal viral infection. The emigration and perivascular accumulation process is not associated with destruction of the vascular wall beyond the transient structural alterations that mediate transmigration. Indeed, the transcriptome and proteome of affected mouse brains reflect this observation and confirm the presence of a pro-inflammatory environment (neuroinflammation).

Studies in fatal COVID-19 cases and in mouse models have reported evidence of BBB impairment [7, 83, 122–127]. Such dramatic changes were not observed in the present study where morphologically, the ultrastructural evidence of an increased amount of glycogen in AEF was the only hint towards, albeit low key, BBB dysfunction. In one publication, the authors provided a definition of the processes and mechanisms they consider as components of BBB disruption; besides loss of tight junction integrity and increased transcytosis (we did not find any evidence of either with any of our methodological approaches), they listed alterations in transport properties and an increased expression of leukocyte adhesion molecules as components of BBB disruption [101]. When we investigated the BBB/NVU in more detail with our bulk-omics approach, we revealed a significantly altered expression of genes known to be associated with the function of endothelial cells, such as leukocyte adhesion molecules (*Vcam1*, *Icam1*, *Selp*, *Sele, Aoc3*) and transporters for nucleotides and nucleobases (*Slc28a2, Slc43a3*), organic anions (*Slc22a8*), neutral amino acids (*Slc1a5*), iron (*Slc40a1*) as well as glycosylated proteins (*Ager*), transferrin (*Tfrc*), and an ATP-dependent transporter (*Abcc6*) [81, 93, 102–104, 107–112, 128] at least at transcriptome level, which could suggest some minimal functional alterations of the BBB. Increased amounts of glycogen in AEF have been linked to BBB disruption by one group [83]. Interestingly, while we found ultrastructural evidence of this in the SARS-CoV-2 Delta infected brains, it was seen independent of the presence of (peri)vascular infiltrates. As SARS-CoV-2 has been reported to manipulate the glucose metabolism [129, 130] and astrocytes are known to store glycogen to support neuronal function [131], we hypothesise that the widespread neuronal infection may have resulted in a higher glucose demand, leading to increased capture of glucose from the blood and storage in the astrocytes as glycogen to help facilitate maintenance of neuron function [132]. This would be supported by the observed moderate (although below our cutoff) but significant increase of the (endothelial) glucose transporter *Slc5a1*. Overall, our data suggest that extensive neuronal SARS-CoV-2 Delta infection in K18-hACE2 mice induces minimal “non-disruptive” BBB dysfunction [133].

In line with our other observations, the metabolome and lipidome of the infected brains exhibited only minimal alterations, restricted to a few compounds associated with oxidative stress, inflammation and immune/antiviral response.

Taken together, our results clearly separate SARS-CoV-2 from truly neuropathic viruses such as Japanese encephalitis virus (JEV), since infection with the latter induces neuronal damage and severe encephalitis associated with alterations of tight junction proteins that represent truly disruptive changes of the BBB, based on structural damage of the bloods vessels, as well as more pronounced metabolic changes in the brain [133–135].

Further studies are warranted that fully elucidate the direct effects of the virus on neurons and to pinpoint the primary insults that trigger the observed processes in SARS-CoV-2 infection of the brain.

## Conclusions

In the present study we showed that intranasal challenge of K18-hACE2 mice with SARS-CoV-2 Delta can lead to widespread neuronal infection and mild neuroinflammation also at a very low infectious dose. The brain reaction is reflected by subtle but distinct changes at molecular level, highlighting limited neuroinflammation with leukocyte recruitment and activation and, possible, minimal BBB dysfunction. We confirmed that SARS-CoV-2 does not target the brain vasculature and hence hypothesise that the observed neuroinflammatory processes are initiated by the infected neurons and mediated by activated glial cells. These findings help to understand the general brain reaction to neuronal infection with viruses that are not overtly cytopathic.

## List of abbreviations

ACE2: angiotensin-converting enzyme 2
AEF: astrocytic endfeet
α-SMA: alpha smooth muscle actin
BBB: blood-brain barrier
BP: biological process
CNS: central nervous system
COVID-19: coronavirus disease 2019
DPI: days post infection
DTT: dithiothreitol
FDR: false discovery rate
GA: glutaraldehyde
GO: Gene Ontology
GSEA: gene set enrichment analysis
H: hour
hACE2: human angiotensin-converting enzyme 2
HE: haematoxylin and eosin
IAM: iodoacetamide
Iba1: ionized calcium binding adaptor molecule 1
IHC: immunohistochemistry
Immune-EM: immune electron microscopy
JEV: Japanese encephalitis virus
K18-hACE2: human (h)ACE2 under the K18 promotor
LC-IM-MS: Liquid chromatography–coupled ion mobility mass spectrometry
LC-MS: Liquid chromatography mass spectrometry
Ly6G: lymphocyte antigen 6 family member G
Min: minutes
NES: normalised enrichment score
NBF: neutral buffered formalin
NP: nucleocapsid protein
NVU: neurovascular unit
PBS: phosphate buffered saline
PCA: principal component analysis
PDGFR-β: platelet-derived growth factor receptor beta
PECAM-1: platelet endothelial cell adhesion molecule 1
PFU: plaque-forming unit
QC: quality control
ppm: parts per million
rpm: revolutions per minute
RT: room temperature
RT-qPCR: reverse transcription quantitative real-time
PCR SARS-CoV-2: severe acute respiratory syndrome coronavirus 2
Sec: seconds
SPF: specific-pathogen free
TEM: transmission electron microscopy
VOC: variants of concern

## Declarations

### Ethics approval and consent to participate

Animal work was approved by the local University of Liverpool Animal Welfare and Ethical Review Body and performed under UK Home Office Project Licence PP4715265. It was undertaken in accordance with locally approved risk assessments and standard operating procedures.

### Consent for publication

Not applicable.

### Competing interests

The authors declare that they have no competing interests.

## Funding

This work was funded by the European Union’s Horizon Europe Research and Innovation Programme under grant agreement No 101057553 and the Swiss State Secretariat for Education, Research and Innovation (SERI) under contract number 22.00094.

## Authors’ contribution

AK (Anja Kipar), JPS, FS, UH, GB: conception and design of the project. SDN, BH, RPR, AK (Adam Kirby), PS, LN, ME, MZ, MR: methodology. PS, AK (Adam Kirby), DM: in vivo experimental work. GB: in vitro infections. SDN, RPR, PS, BH, AK (Adam Kirby), LN, ME, MZ, MR: data collection. SDN, RPR, LN, AK (Anja Kipar), UH, MZ, MR, AO: data analysis. SDN, AK (Anja Kipar): writing of the original draft. SDN, AK (Anja Kipar), RPR, LN, ME, EE, JPS, GB, UH, FS, AO: review and editing. AK (Anja Kipar), JPS, GB, EE: funding acquisition. AK (Anja Kipar): supervision of the project. All authors read and approved the manuscript.

## Acknowledgements

We are grateful to the technical staff in the Histology Laboratory, Institute of Veterinary Pathology, Vetsuisse Faculty, University of Zurich (IVPZ), for excellent technical support. We would like to thank Professor Wei-Chung Cheng and Dr. Pei-Chun Shen at Program for Cancer Molecular Biology and Drug Discovery, College of Medicine, China Medical University, for their help in applying their R package, LipidSigR, to our analysis.

## Notes

### Competing Interest Statement

The authors have declared no competing interest.

### Summary of Updates

The study has been extended by including proteomic, metabolomic and lipidomic analyses of the brains. All sections (Materials and Methods, Results, Discussion) including Figures and Supplemental material files have been updated to incorporate this additional work.

## References

1. Milross L, Majo J, Cooper N, Kaye PM, Bayraktar O, Filby A, Fisher AJ. Post-mortem lung tissue: the fossil record of the pathophysiology and immunopathology of severe COVID-19. The Lancet Respiratory Medicine. 2022;10:95–106. doi:10.1016/S2213-2600(21)00408-2.

2. Monje M, Iwasaki A. The neurobiology of long COVID. Neuron 2022. doi:10.1016/j.neuron.2022.10.006.

3. Mehandru S, Merad M. Pathological sequelae of long-haul COVID. Nat Immunol. 2022;23:194– 202. doi:10.1038/s41590-021-01104-y.

4. Chou SH-Y, Beghi E, Helbok R, Moro E, Sampson J, Altamirano V, et al. Global incidence of neurological manifestations among patients hospitalized with COVID-19-A report for the GCS-NeuroCOVID Consortium and the ENERGY Consortium. JAMA Netw Open. 2021;4:e2112131. doi:10.1001/jamanetworkopen.2021.12131.

5. Ross Russell AL, Hardwick M, Jeyanantham A, White LM, Deb S, Burnside G, et al. Spectrum, risk factors and outcomes of neurological and psychiatric complications of COVID-19: a UK-wide cross-sectional surveillance study. Brain Commun. 2021;3:fcab168. doi:10.1093/braincomms/fcab168.

6. Khedr EM, Abo-Elfetoh N, Deaf E, Hassan HM, Amin MT, Soliman RK, et al. Surveillance study of acute neurological manifestations among 439 Egyptian patients with COVID-19 in Assiut and Aswan university hospitals. Neuroepidemiology. 2021;55:109–18. doi:10.1159/000513647.

7. Meinhardt J, Streit S, Dittmayer C, Manitius RV, Radbruch H, Heppner FL. The neurobiology of SARS-CoV-2 infection. Nat Rev Neurosci. 2024;25:30–42. doi:10.1038/s41583-023-00769-8.

8. Mukerji SS, Solomon IH. What can we learn from brain autopsies in COVID-19? Neurosci Lett. 2021;742:135528. doi:10.1016/j.neulet.2020.135528.

9. Solomon IH, Singh A, Folkerth RD, Mukerji SS. What can we still learn from brain autopsies in COVID-19? Semin Neurol. 2023;43:195–204. doi:10.1055/s-0043-1767716.

10. Matschke J, Lütgehetmann M, Hagel C, Sperhake JP, Schröder AS, Edler C, et al. Neuropathology of patients with COVID-19 in Germany: a post-mortem case series. The Lancet Neurology. 2020;19:919–29. doi:10.1016/S1474-4422(20)30308-2.

11. Bocci M, Oudenaarden C, Sàenz-Sardà X, Simrén J, Edén A, Sjölund J, et al. Infection of brain pericytes underlying neuropathology of COVID-19 patients. Int J Mol Sci 2021. doi:10.3390/ijms222111622.

12. Agrawal S, Farfel JM, Arfanakis K, Al-Harthi L, Shull T, Teppen TL, et al. Brain autopsies of critically ill COVID-19 patients demonstrate heterogeneous profile of acute vascular injury, inflammation and age-linked chronic brain diseases. Acta Neuropathol Commun. 2022;10:186. doi:10.1186/s40478-022-01493-7.

13. Puelles VG, Lütgehetmann M, Lindenmeyer MT, Sperhake JP, Wong MN, Allweiss L, et al. Multiorgan and renal tropism of SARS-CoV-2. N Engl J Med. 2020;383:590–2. doi:10.1056/NEJMc2011400.

14. Deigendesch N, Sironi L, Kutza M, Wischnewski S, Fuchs V, Hench J, et al. Correlates of critical illness-related encephalopathy predominate postmortem COVID-19 neuropathology. Acta Neuropathol. 2020;140:583–6. doi:10.1007/s00401-020-02213-y.

15. Meinhardt J, Radke J, Dittmayer C, Franz J, Thomas C, Mothes R, et al. Olfactory transmucosal SARS-CoV-2 invasion as a port of central nervous system entry in individuals with COVID-19. Nat Neurosci. 2021;24:168–75. doi:10.1038/s41593-020-00758-5.

16. Fabbri VP, Riefolo M, Lazzarotto T, Gabrielli L, Cenacchi G, Gallo C, et al. COVID-19 and the brain: The neuropathological Italian experience on 33 adult autopsies. Biomolecules 2022. doi:10.3390/biom12050629.

17. McCray PB, Pewe L, Wohlford-Lenane C, Hickey M, Manzel L, Shi L, et al. Lethal infection of K18-hACE2 mice infected with severe acute respiratory syndrome coronavirus. J Virol. 2007;81:813–21. doi:10.1128/JVI.02012-06.

18. Carossino M, Kenney D, O’Connell AK, Montanaro P, Tseng AE, Gertje HP, et al. Fatal neurodissemination and SARS-CoV-2 tropism in K18-hACE2 mice is only partially dependent on hACE2 expression. Viruses 2022. doi:10.3390/v14030535.

19. Seehusen F, Clark JJ, Sharma P, Bentley EG, Kirby A, Subramaniam K, et al. Neuroinvasion and neurotropism by SARS-CoV-2 variants in the K18-hACE2 mouse. Viruses 2022. doi:10.3390/v14051020.

20. Kumari P, Rothan HA, Natekar JP, Stone S, Pathak H, Strate PG, et al. Neuroinvasion and encephalitis following intranasal inoculation of SARS-CoV-2 in K18-hACE2 mice. Viruses 2021. doi:10.3390/v13010132.

21. Song E, Zhang C, Israelow B, Lu-Culligan A, Prado AV, Skriabine S, et al. Neuroinvasion of SARS-CoV-2 in human and mouse brain. J Exp Med 2021. doi:10.1084/jem.20202135.

22. Hassan AO, Case JB, Winkler ES, Thackray LB, Kafai NM, Bailey AL, et al. A SARS-CoV-2 infection model in mice demonstrates protection by neutralizing antibodies. Cell. 2020;182:744–753.e4. doi:10.1016/j.cell.2020.06.011.

23. Fumagalli V, Ravà M, Marotta D, Di Lucia P, Laura C, Sala E, et al. Administration of aerosolized SARS-CoV-2 to K18-hACE2 mice uncouples respiratory infection from fatal neuroinvasion. Sci Immunol. 2022;7:eabl9929. doi:10.1126/sciimmunol.abl9929.

24. Winkler ES, Bailey AL, Kafai NM, Nair S, McCune BT, Yu J, et al. SARS-CoV-2 infection of human ACE2-transgenic mice causes severe lung inflammation and impaired function. Nat Immunol. 2020;21:1327–35. doi:10.1038/s41590-020-0778-2.

25. Gan ES, Syenina A, Linster M, Ng B, Zhang SL, Watanabe S, et al. A mouse model of lethal respiratory dysfunction for SARS-CoV-2 infection. Antiviral Res. 2021;193:105138. doi:10.1016/j.antiviral.2021.105138.

26. Olivarria GM, Cheng Y, Furman S, Pachow C, Hohsfield LA, Smith-Geater C, et al. Microglia do not restrict SARS-CoV-2 replication following infection of the central nervous system of K18-human ACE2 transgenic mice. J Virol. 2022;96:e0196921. doi:10.1128/jvi.01969-21.

27. Tarrés-Freixas F, Trinité B, Pons-Grífols A, Romero-Durana M, Riveira-Muñoz E, Ávila-Nieto C, et al. Heterogeneous infectivity and pathogenesis of SARS-CoV-2 variants Beta, Delta and Omicron in transgenic K18-hACE2 and wildtype mice. Front Microbiol. 2022;13:840757. doi:10.3389/fmicb.2022.840757.

28. Vidal E, López-Figueroa C, Rodon J, Pérez M, Brustolin M, Cantero G, et al. Chronological brain lesions after SARS-CoV-2 infection in hACE2-transgenic mice. Vet Pathol. 2022;59:613–26. doi:10.1177/03009858211066841.

29. Yinda CK, Port JR, Bushmaker T, Offei Owusu I, Purushotham JN, Avanzato VA, et al. K18-hACE2 mice develop respiratory disease resembling severe COVID-19. PLoS Pathog. 2021;17:e1009195. doi:10.1371/journal.ppat.1009195.

30. Zhang L, Zhou L, Bao L, Liu J, Zhu H, Lv Q, et al. SARS-CoV-2 crosses the blood-brain barrier accompanied with basement membrane disruption without tight junctions alteration. Signal Transduct Target Ther. 2021;6:337. doi:10.1038/s41392-021-00719-9.

31. Oladunni FS, Park J-G, Pino PA, Gonzalez O, Akhter A, Allué-Guardia A, et al. Lethality of SARS-CoV-2 infection in K18 human angiotensin-converting enzyme 2 transgenic mice. Nat Commun. 2020;11:6122. doi:10.1038/s41467-020-19891-7.

32. Lee KS, Wong TY, Russ BP, Horspool AM, Miller OA, Rader NA, et al. SARS-CoV-2 Delta variant induces enhanced pathology and inflammatory responses in K18-hACE2 mice. PLoS One. 2022;17:e0273430. doi:10.1371/journal.pone.0273430.

33. Yang J-H, Yang M-S, Kim D-M, Kim B, Tark D, Kang S-M, Lee G-H. Delta (B1.617.2) variant of SARS-CoV-2 induces severe neurotropic patterns in K18-hACE2 mice. Scientific reports. 2023;13:3303. doi:10.1038/s41598-023-29909-x.

34. Sieracka J, Sieracki P, Kozera G, Szurowska E, Gulczyński J, Sobolewski P, et al. COVID-19 - neuropathological point of view, pathobiology, and dilemmas after the first year of the pandemic struggle. Folia Neuropathol. 2021;59:1–16. doi:10.5114/fn.2021.105128.

35. Chakravarty D, Das Sarma J. Murine-β-coronavirus-induced neuropathogenesis sheds light on CNS pathobiology of SARS-CoV2. J Neurovirol. 2021;27:197–216. doi:10.1007/s13365-021-00945-5.

36. Butowt R, Meunier N, Bryche B, von Bartheld CS. The olfactory nerve is not a likely route to brain infection in COVID-19: a critical review of data from humans and animal models. Acta Neuropathol. 2021;141:809–22. doi:10.1007/s00401-021-02314-2.

37. Martínez-Mármol R, Giordano-Santini R, Kaulich E, Cho A-N, Przybyla M, Riyadh MA, et al. SARS-CoV-2 infection and viral fusogens cause neuronal and glial fusion that compromises neuronal activity. Sci Adv. 2023;9:eadg2248. doi:10.1126/sciadv.adg2248.

38. Pinja K, Janika R, Tania Q, Ravi O, Saber SH, Shaker M, et al. SARS-CoV-2 infection in hiPSC-derived neurons is cathepsin-dependent and causes accumulation of HIF1ɑ and phosphorylated tau; 2024.

39. Jeon D, Kim S-H, Kim J, Jeong H, Uhm C, Oh H, et al. Discovery of a new long COVID mouse model via systemic histopathological comparison of SARS-CoV-2 intranasal and inhalation infection. Biochim Biophys Acta Mol Basis Dis. 2024;1870:167347. doi:10.1016/j.bbadis.2024.167347.

40. Patterson EI, Prince T, Anderson ER, Casas-Sanchez A, Smith SL, Cansado-Utrilla C, et al. Methods of inactivation of SARS-CoV-2 for downstream biological assays. J Infect Dis. 2020;222:1462–7. doi:10.1093/infdis/jiaa507.

41. Clark JJ, Penrice-Randal R, Sharma P, Dong X, Pennington SH, Marriott AE, et al. Sequential infection with Influenza A virus followed by Severe Acute Respiratory Syndrome Coronavirus 2 (SARS-CoV-2) leads to more severe disease and encephalitis in a mouse model of COVID-19. Viruses 2024. doi:10.3390/v16060863.

42. Cabrera LE, Jokiranta ST, Mäki S, Miettinen S, Kant R, Kareinen L, et al. The assembly of neutrophil inflammasomes during COVID-19 is mediated by type I interferons. PLoS Pathog. 2024;20:e1012368. doi:10.1371/journal.ppat.1012368.

43. Kang Y, Hepojoki J, Maldonado RS, Mito T, Terzioglu M, Manninen T, et al. Ancestral allele of DNA polymerase gamma modifies antiviral tolerance. Nature. 2024;628:844–53. doi:10.1038/s41586-024-07260-z.

44. Winkler EA, Bell RD, Zlokovic BV. Pericyte-specific expression of PDGF beta receptor in mouse models with normal and deficient PDGF beta receptor signaling. Mol Neurodegener. 2010;5:32. doi:10.1186/1750-1326-5-32.

45. Hubbard JA, Hsu MS, Seldin MM, Binder DK. Expression of the astrocyte water channel aquaporin-4 in the mouse brain. ASN Neuro 2015. doi:10.1177/1759091415605486.

46. Crouch EE, Doetsch F. FACS isolation of endothelial cells and pericytes from mouse brain microregions. Nat Protoc. 2018;13:738–51. doi:10.1038/nprot.2017.158.

47. Grant RI, Hartmann DA, Underly RG, Berthiaume A-A, Bhat NR, Shih AY. Organizational hierarchy and structural diversity of microvascular pericytes in adult mouse cortex. Journal of Cerebral Blood Flow & Metabolism. 2019;39:411–25. doi:10.1177/0271678X17732229.

48. Greene C, Hanley N, Campbell M. Claudin-5: gatekeeper of neurological function. Fluids Barriers CNS. 2019;16:3. doi:10.1186/s12987-019-0123-z.

49. Gautam J, Zhang X, Yao Y. The role of pericytic laminin in blood brain barrier integrity maintenance. Scientific reports. 2016;6:36450. doi:10.1038/srep36450.

50. Ineichen BV, Okar SV, Proulx ST, Engelhardt B, Lassmann H, Reich DS. Perivascular spaces and their role in neuroinflammation. Neuron. 2022;110:3566–81. doi:10.1016/j.neuron.2022.10.024.

51. Chen S, Zhou Y, Chen Y, Gu J. fastp: an ultra-fast all-in-one FASTQ preprocessor. Bioinformatics. 2018;34:i884–i890. doi:10.1093/bioinformatics/bty560.

52. Patro R, Duggal G, Love MI, Irizarry RA, Kingsford C. Salmon provides fast and bias-aware quantification of transcript expression. Nat Methods. 2017;14:417–9. doi:10.1038/nmeth.4197.

53. Dyer SC, Austine-Orimoloye O, Azov AG, Barba M, Barnes I, Barrera-Enriquez VP, et al. Ensembl 2025. Nucleic Acids Res. 2025;53:D948–D957. doi:10.1093/nar/gkae1071.

54. Soneson C, Love MI, Robinson MD. Differential analyses for RNA-seq: transcript-level estimates improve gene-level inferences. F1000Res. 2015;4:1521. doi:10.12688/f1000research.7563.2.

55. Chen Y, Chen L, Lun ATL, Baldoni PL, Smyth GK. edgeR v4: powerful differential analysis of sequencing data with expanded functionality and improved support for small counts and larger datasets. Nucleic Acids Res 2025. doi:10.1093/nar/gkaf018.

56. Blighe Kevin, Lun Aaron. PCAtools: PCAtools: Everything Principal Components Analysis; 2024.

57. The Gene Ontology Consortium. The Gene Ontology knowledgebase in 2026. Nucleic Acids Res. 2026;54:D1779–D1792. doi:10.1093/nar/gkaf1292.

58. Ashburner M, Ball CA, Blake JA, Botstein D, Butler H, Cherry JM, et al. Gene ontology: tool for the unification of biology. The Gene Ontology Consortium. Nat Genet. 2000;25:25–9. doi:10.1038/75556.

59. Xu S, Hu E, Cai Y, Xie Z, Luo X, Zhan L, et al. Using clusterProfiler to characterize multiomics data. Nat Protoc. 2024;19:3292–320. doi:10.1038/s41596-024-01020-z.

60. Yu G. Thirteen years of clusterProfiler. Innovation (Camb). 2024;5:100722. doi:10.1016/j.xinn.2024.100722.

61. Wu T, Hu E, Xu S, Chen M, Guo P, Dai Z, et al. clusterProfiler 4.0: A universal enrichment tool for interpreting omics data. Innovation (Camb). 2021;2:100141. doi:10.1016/j.xinn.2021.100141.

62. Yu G, Wang L-G, Han Y, He Q-Y. clusterProfiler: an R package for comparing biological themes among gene clusters. OMICS. 2012;16:284–7. doi:10.1089/omi.2011.0118.

63. Hughes CS, Moggridge S, Müller T, Sorensen PH, Morin GB, Krijgsveld J. Single-pot, solid-phase-enhanced sample preparation for proteomics experiments. Nat Protoc. 2019;14:68–85. doi:10.1038/s41596-018-0082-x.

64. Meier F, Brunner A-D, Koch S, Koch H, Lubeck M, Krause M, et al. Online Parallel Accumulation-Serial Fragmentation (PASEF) with a Novel Trapped Ion Mobility Mass Spectrometer. Mol Cell Proteomics. 2018;17:2534–45. doi:10.1074/mcp.TIR118.000900.

65. Kong AT, Leprevost FV, Avtonomov DM, Mellacheruvu D, Nesvizhskii AI. MSFragger: ultrafast and comprehensive peptide identification in mass spectrometry-based proteomics. Nat Methods. 2017;14:513–20. doi:10.1038/nmeth.4256.

66. Yu F, Haynes SE, Nesvizhskii AI. IonQuant Enables Accurate and Sensitive Label-Free Quantification With FDR-Controlled Match-Between-Runs. Mol Cell Proteomics. 2021;20:100077. doi:10.1016/j.mcpro.2021.100077.

67. UniProt Consortium. UniProt: the Universal Protein Knowledgebase in 2025. Nucleic Acids Res. 2025;53:D609–D617. doi:10.1093/nar/gkae1010.

68. Feng Z, Fang P, Zheng H, Zhang X. DEP2: an upgraded comprehensive analysis toolkit for quantitative proteomics data. Bioinformatics 2023. doi:10.1093/bioinformatics/btad526.

69. Zhang X, Smits AH, van Tilburg GB, Ovaa H, Huber W, Vermeulen M. Proteome-wide identification of ubiquitin interactions using UbIA-MS. Nat Protoc. 2018;13:530–50. doi:10.1038/nprot.2017.147.

70. Ritchie ME, Phipson B, Di Wu, Hu Y, Law CW, Shi W, Smyth GK. limma powers differential expression analyses for RNA-sequencing and microarray studies. Nucleic Acids Res. 2015;43:e47. doi:10.1093/nar/gkv007.

71. Sharma AK, Khandelwal R, Zurkovic J, Long F, Wang T, Dewal RS, et al. DGAT-driven futile lipid cycling has a pronounced, yet concealed, thermogenic function. Cell Metab 2026. doi:10.1016/j.cmet.2025.12.009.

72. Shen X, Yan H, Wang C, Gao P, Johnson CH, Snyder MP. TidyMass an object-oriented reproducible analysis framework for LC-MS data. Nat Commun. 2022;13:4365. doi:10.1038/s41467-022-32155-w.

73. Idkowiak J, Dehairs J, Schwarzerová J, Olešová D, Truong JXM, Kvasnička A, et al. Best practices and tools in R and Python for statistical processing and visualization of lipidomics and metabolomics data. Nat Commun. 2025;16:8714. doi:10.1038/s41467-025-63751-1.

74. Kolde Raivo. pheatmap: Pretty heatmaps; 2025.

75. Lin W-J, Shen P-C, Liu H-C, Cho Y-C, Hsu M-K, Lin I-C, et al. LipidSig: a web-based tool for lipidomic data analysis. Nucleic Acids Res. 2021;49:W336–W345. doi:10.1093/nar/gkab419.

76. Liu C-H, Shen P-C, Lin W-J, Liu H-C, Tsai M-H, Huang T-Y, et al. LipidSig 2.0: integrating lipid characteristic insights into advanced lipidomics data analysis. Nucleic Acids Res. 2024;52:W390–W397. doi:10.1093/nar/gkae335.

77. Liu C-H, Shen P-C, Tsai M-H, Liu H-C, Lin W-J, Lai Y-L, et al. LipidSigR: a R-based solution for integrated lipidomics data analysis and visualization. Bioinform Adv. 2025;5:vbaf047. doi:10.1093/bioadv/vbaf047.

78. Rosiak M, Schreiner T, Beythien G, Leitzen E, Ulianytska A, Allnoch L, et al. Spatiotemporal Characterization of Changes in the Respiratory Tract and the Nervous System, Including the Eyes in SARS-CoV-2-Infected K18-hACE2 Mice. Viruses 2025. doi:10.3390/v17070963.

79. Gouveia-Freitas K, Bastos-Leite AJ. Perivascular spaces and brain waste clearance systems: relevance for neurodegenerative and cerebrovascular pathology. Neuroradiology. 2021;63:1581–97. doi:10.1007/s00234-021-02718-7.

80. Iadecola C. The Neurovascular Unit Coming of Age: A Journey through Neurovascular Coupling in Health and Disease. Neuron. 2017;96:17–42. doi:10.1016/j.neuron.2017.07.030.

81. Zhao Z, Nelson AR, Betsholtz C, Zlokovic BV. Establishment and dysfunction of the blood-brain barrier. Cell. 2015;163:1064–78. doi:10.1016/j.cell.2015.10.067.

82. Mathiisen TM, Lehre KP, Danbolt NC, Ottersen OP. The perivascular astroglial sheath provides a complete covering of the brain microvessels: an electron microscopic 3D reconstruction. Glia. 2010;58:1094–103. doi:10.1002/glia.20990.

83. Nahirney PC, Reeson P, Brown CE. Ultrastructural analysis of blood-brain barrier breakdown in the peri-infarct zone in young adult and aged mice. J Cereb Blood Flow Metab. 2016;36:413–25. doi:10.1177/0271678X15608396.

84. Cui A, Huang T, Li S, Ma A, Pérez JL, Sander C, et al. Dictionary of immune responses to cytokines at single-cell resolution. Nature. 2024;625:377–84. doi:10.1038/s41586-023-06816-9.

85. Hughes CE, Nibbs RJB. A guide to chemokines and their receptors. FEBS J. 2018;285:2944–71. doi:10.1111/febs.14466.

86. Li J, Xu P, Hong Y, Xie Y, Peng M, Sun R, et al. Lipocalin-2-mediated astrocyte pyroptosis promotes neuroinflammatory injury via NLRP3 inflammasome activation in cerebral ischemia/reperfusion injury. J Neuroinflammation. 2023;20:148. doi:10.1186/s12974-023-02819-5.

87. Jha MK, Lee S, Park DH, Kook H, Park K-G, Lee I-K, Suk K. Diverse functional roles of lipocalin-2 in the central nervous system. Neurosci Biobehav Rev. 2015;49:135–56. doi:10.1016/j.neubiorev.2014.12.006.

88. van Zuylen WJ, Garceau V, Idris A, Schroder K, Irvine KM, Lattin JE, et al. Macrophage activation and differentiation signals regulate schlafen-4 gene expression: evidence for Schlafen-4 as a modulator of myelopoiesis. PLoS One. 2011;6:e15723. doi:10.1371/journal.pone.0015723.

89. Das A, Dinh PX, Panda D, Pattnaik AK. Interferon-inducible protein IFI35 negatively regulates RIG-I antiviral signaling and supports vesicular stomatitis virus replication. J Virol. 2014;88:3103–13. doi:10.1128/JVI.03202-13.

90. Yang H, Winkler W, Wu X. Interferon Inducer IFI35 regulates RIG-I-mediated innate antiviral response through mutual antagonism with Influenza protein NS1. J Virol 2021. doi:10.1128/JVI.00283-21.

91. Yu Y, Xu N, Cheng Q, Deng F, Liu M, Zhu A, et al. IFP35 as a promising biomarker and therapeutic target for the syndromes induced by SARS-CoV-2 or influenza virus. Cell Rep. 2021;37:110126. doi:10.1016/j.celrep.2021.110126.

92. Yan M, Sun Z, Zhang S, Yang G, Jiang X, Wang G, et al. SOCS modulates JAK-STAT pathway as a novel target to mediate the occurrence of neuroinflammation: Molecular details and treatment options. Brain Res Bull. 2024;213:110988. doi:10.1016/j.brainresbull.2024.110988.

93. Marchetti L, Engelhardt B. Immune cell trafficking across the blood-brain barrier in the absence and presence of neuroinflammation. Vasc Biol. 2020;2:H1–H18. doi:10.1530/VB-19-0033.

94. Bouwman AC, van Daalen KR, Crnko S, ten Broeke T, Bovenschen N. Intracellular and Extracellular Roles of Granzyme K. Front Immunol. 2021;12:677707. doi:10.3389/fimmu.2021.677707.

95. Odobasic D, Kitching AR, Yang Y, O’Sullivan KM, Muljadi RCM, Edgtton KL, et al. Neutrophil myeloperoxidase regulates T-cell-driven tissue inflammation in mice by inhibiting dendritic cell function. Blood. 2013;121:4195–204. doi:10.1182/blood-2012-09-456483.

96. Nourhaghighi N, Teichert-Kuliszewska K, Davis J, Stewart DJ, Nag S. Altered expression of angiopoietins during blood-brain barrier breakdown and angiogenesis. Lab Invest. 2003;83:1211–22. doi:10.1097/01.lab.0000082383.40635.fe.

97. Nag S, Papneja T, Venugopalan R, Stewart DJ. Increased angiopoietin2 expression is associated with endothelial apoptosis and blood-brain barrier breakdown. Lab Invest. 2005;85:1189–98. doi:10.1038/labinvest.3700325.

98. Gurnik S, Devraj K, Macas J, Yamaji M, Starke J, Scholz A, et al. Angiopoietin-2-induced blood-brain barrier compromise and increased stroke size are rescued by VE-PTP-dependent restoration of Tie2 signaling. Acta Neuropathol. 2016;131:753–73. doi:10.1007/s00401-016-1551-3.

99. Tremblay T-L, Alata W, Slinn J, Baumann E, Delaney CE, Moreno M, et al. The proteome of the blood-brain barrier in rat and mouse: highly specific identification of proteins on the luminal surface of brain microvessels by in vivo glycocapture. Fluids Barriers CNS. 2024;21:23. doi:10.1186/s12987-024-00523-x.

100. Daneman R, Zhou L, Agalliu D, Cahoy JD, Kaushal A, Barres BA. The mouse blood-brain barrier transcriptome: a new resource for understanding the development and function of brain endothelial cells. PLoS One. 2010;5:e13741. doi:10.1371/journal.pone.0013741.

101. Munji RN, Soung AL, Weiner GA, Sohet F, Semple BD, Trivedi A, et al. Profiling the mouse brain endothelial transcriptome in health and disease models reveals a core blood-brain barrier dysfunction module. Nat Neurosci. 2019;22:1892–902. doi:10.1038/s41593-019-0497-x.

102. Boyer DS, Rippmann JF, Ehrlich MS, Bakker RA, Chong V, Nguyen QD. Amine oxidase copper-containing 3 (AOC3) inhibition: a potential novel target for the management of diabetic retinopathy. Int J Retina Vitreous. 2021;7:30. doi:10.1186/s40942-021-00288-7.

103. Unzeta M, Hernàndez-Guillamon M, Sun P, Solé M. SSAO/VAP-1 in Cerebrovascular Disorders: A Potential Therapeutic Target for Stroke and Alzheimer’s Disease. Int J Mol Sci 2021. doi:10.3390/ijms22073365.

104. Noda K, Nakao S, Zandi S, Engelstädter V, Mashima Y, Hafezi-Moghadam A. Vascular adhesion protein-1 regulates leukocyte transmigration rate in the retina during diabetes. Exp Eye Res. 2009;89:774–81. doi:10.1016/j.exer.2009.07.010.

105. Gong C-X, Shi P-X, Huang Y-J, Dai Y, Hu L-L, Cheng X-F, et al. Apolipoprotein D, a Novel Ligand for CD36, Is Essential for Blood-Brain Barrier Integrity. Circulation 2026. doi:10.1161/CIRCULATIONAHA.125.077356.

106. Bangsow T, Baumann E, Bangsow C, Jaeger MH, Pelzer B, Gruhn P, et al. The epithelial membrane protein 1 is a novel tight junction protein of the blood-brain barrier. J Cereb Blood Flow Metab. 2008;28:1249–60. doi:10.1038/jcbfm.2008.19.

107. Furukawa J, Inoue K, Maeda J, Yasujima T, Ohta K, Kanai Y, et al. Functional identification of SLC43A3 as an equilibrative nucleobase transporter involved in purine salvage in mammals. Scientific reports. 2015;5:15057. doi:10.1038/srep15057.

108. Wallgard E, Larsson E, He L, Hellström M, Armulik A, Nisancioglu MH, et al. Identification of a core set of 58 gene transcripts with broad and specific expression in the microvasculature. Arterioscler Thromb Vasc Biol. 2008;28:1469–76. doi:10.1161/ATVBAHA.108.165738.

109. Warren MS, Zerangue N, Woodford K, Roberts LM, Tate EH, Feng B, et al. Comparative gene expression profiles of ABC transporters in brain microvessel endothelial cells and brain in five species including human. Pharmacol Res. 2009;59:404–13. doi:10.1016/j.phrs.2009.02.007.

110. de Vilder EYG, Cardoen S, Hosen MJ, Le Saux O, de Zaeytijd J, Leroy BP, et al. Pathogenic variants in the ABCC6 gene are associated with an increased risk for ischemic stroke. Brain Pathol. 2018;28:822–31. doi:10.1111/bpa.12620.

111. Zheng H, Guo X, Kang S, Li Z, Tian T, Li J, et al. Cdh5-mediated Fpn1 deletion exerts neuroprotective effects during the acute phase and inhibitory effects during the recovery phase of ischemic stroke. Cell Death Dis. 2023;14:161. doi:10.1038/s41419-023-05688-1.

112. Donovan A, Lima CA, Pinkus JL, Pinkus GS, Zon LI, Robine S, Andrews NC. The iron exporter ferroportin/Slc40a1 is essential for iron homeostasis. Cell Metab. 2005;1:191–200. doi:10.1016/j.cmet.2005.01.003.

113. Chen F, Elgaher WAM, Winterhoff M, Büssow K, Waqas FH, Graner E, et al. Citraconate inhibits ACOD1 (IRG1) catalysis, reduces interferon responses and oxidative stress, and modulates inflammation and cell metabolism. Nat Metab. 2022;4:534–46. doi:10.1038/s42255-022-00577-x.

114. McGettrick AF, O’Neill LA. Two for the price of one: itaconate and its derivatives as an anti-infective and anti-inflammatory immunometabolite. Curr Opin Immunol. 2023;80:102268. doi:10.1016/j.coi.2022.102268.

115. Steiner S, Kratzel A, Barut GT, Lang RM, Aguiar Moreira E, Thomann L, et al. SARS-CoV-2 biology and host interactions. Nat Rev Microbiol. 2024;22:206–25. doi:10.1038/s41579-023-01003-z.

116. Barroso M, Kao D, Blom HJ, Tavares de Almeida I, Castro R, Loscalzo J, Handy DE. S-adenosylhomocysteine induces inflammation through NFkB: A possible role for EZH2 in endothelial cell activation. Biochim Biophys Acta. 2016;1862:82–92. doi:10.1016/j.bbadis.2015.10.019.

117. Srivastava R, Panda SK, Sen Gupta PS, Chaudhary A, Naaz F, Yadav AK, et al. In silico evaluation of S-adenosyl-L-homocysteine analogs as inhibitors of nsp14-viral cap N7 methyltranferase and PLpro of SARS-CoV-2: synthesis, molecular docking, physicochemical data, ADMET and molecular dynamics simulations studies. J Biomol Struct Dyn. 2025;43:3258–75. doi:10.1080/07391102.2023.2297005.

118. Park Y, Paing YMM, Cho N, Kim C, Yoo J, Choi JW, Lee SH. Quinic Acid Alleviates Behavior Impairment by Reducing Neuroinflammation and MAPK Activation in LPS-Treated Mice. Biomol Ther (Seoul). 2024;32:309–18. doi:10.4062/biomolther.2023.184.

119. Shin YJ, Kim YJ, Lee JE, Kim YS, Lee JW, Kim H, et al. Uric acid regulates α-synuclein transmission in Parkinsonian models. Front Aging Neurosci. 2023;15:1117491. doi:10.3389/fnagi.2023.1117491.

120. Zheng J, Wong L-YR, Li K, Verma AK, Ortiz ME, Wohlford-Lenane C, et al. COVID-19 treatments and pathogenesis including anosmia in K18-hACE2 mice. Nature. 2021;589:603–7. doi:10.1038/s41586-020-2943-z.

121. Nicosia RF, Ligresti G, Caporarello N, Akilesh S, Ribatti D. COVID-19 Vasculopathy: Mounting Evidence for an Indirect Mechanism of Endothelial Injury. Am J Pathol. 2021;191:1374–84. doi:10.1016/j.ajpath.2021.05.007.

122. Bonetto V, Pasetto L, Lisi I, Carbonara M, Zangari R, Ferrari E, et al. Markers of blood-brain barrier disruption increase early and persistently in COVID-19 patients with neurological manifestations. Front Immunol. 2022;13:1070379. doi:10.3389/fimmu.2022.1070379.

123. Etter MM, Martins TA, Kulsvehagen L, Pössnecker E, Duchemin W, Hogan S, et al. Severe Neuro-COVID is associated with peripheral immune signatures, autoimmunity and neurodegeneration: a prospective cross-sectional study. Nat Commun. 2022;13:6777. doi:10.1038/s41467-022-34068-0.

124. Greene C, Connolly R, Brennan D, Laffan A, O’Keeffe E, Zaporojan L, et al. Blood-brain barrier disruption and sustained systemic inflammation in individuals with long COVID-associated cognitive impairment. Nat Neurosci. 2024;27:421–32. doi:10.1038/s41593-024-01576-9.

125. Trevino TN, Almousawi AA, Robinson KF, Fogel AB, Class J, Minshall RD, et al. Caveolin-1 mediates blood-brain barrier permeability, neuroinflammation, and cognitive impairment in SARS-CoV-2 infection. J Neuroimmunol. 2024;388:578309. doi:10.1016/j.jneuroim.2024.578309.

126. Trevino TN, Fogel AB, Otkiran G, Niladhuri SB, Sanborn MA, Class J, et al. Engineered Wnt7a ligands rescue blood-brain barrier and cognitive deficits in a COVID-19 mouse model. Brain. 2024;147:1636–43. doi:10.1093/brain/awae031.

127. Song H, Tomasevich A, Acheampong KK, Schaff DL, Shaffer SM, Dolle J-P, et al. Detection of blood-brain barrier disruption in brains of patients with COVID-19, but no evidence of brain penetration by SARS-CoV-2. Acta Neuropathol. 2023;146:771–5. doi:10.1007/s00401-023-02624-7.

128. Hashimoto R, Takahashi J, Shirakura K, Funatsu R, Kosugi K, Deguchi S, et al. SARS-CoV-2 disrupts respiratory vascular barriers by suppressing Claudin-5 expression. Sci Adv. 2022;8:eabo6783. doi:10.1126/sciadv.abo6783.

129. Liu X, Chen L, Niu H, Chen Y, Chen P, Liu L, Wu R. The bittersweet link between glucose metabolism, cellular microenvironment and viral infection. Virulence. 2025;16:2554302. doi:10.1080/21505594.2025.2554302.

130. Darweesh M, Mohammadi S, Rahmati M, Al-Hamadani M, Al-Harrasi A. Metabolic reprogramming in viral infections: the interplay of glucose metabolism and immune responses. Front Immunol. 2025;16:1578202. doi:10.3389/fimmu.2025.1578202.

131. Hill SA, Bravo-Ferrer I, Čiulkinytė A, Pérez Ramos N, Rossetti I, Colvin C, et al. Molecular profiling of brain endothelial cell to astrocyte endfoot communication in mouse and human. Nat Commun. 2025;16:9750. doi:10.1038/s41467-025-65487-4.

132. Huang S-F, Ogunshola OO. Metabolomic profiling provides new insights into blood-brain barrier regulation. Neural Regen Res. 2021;16:1786–7. doi:10.4103/1673-5374.306077.

133. Varatharaj A, Galea I. The blood-brain barrier in systemic inflammation. Brain Behav Immun. 2017;60:1–12. doi:10.1016/j.bbi.2016.03.010.

134. Su Z, Sun N, Yin C, Zheng X. An untargeted metabolomics analysis in feces and brain of Orthoflaviviruses-infected mice. BMC Microbiol. 2025;25:452. doi:10.1186/s12866-025-04192-0.

135. Li F, Wang Y, Yu L, Cao S, Wang K, Yuan J, et al. Viral Infection of the Central Nervous System and Neuroinflammation Precede Blood-Brain Barrier Disruption during Japanese Encephalitis Virus Infection. J Virol. 2015;89:5602–14. doi:10.1128/JVI.00143-15.

136. Katz K, Shutov O, Lapoint R, Kimelman M, Brister JR, O’Sullivan C. The Sequence Read Archive: a decade more of explosive growth. Nucleic Acids Res. 2022;50:D387–D390. doi:10.1093/nar/gkab1053.

137. Perez-Riverol Y, Bandla C, Kundu DJ, Kamatchinathan S, Bai J, Hewapathirana S, et al. The PRIDE database at 20 years: 2025 update. Nucleic Acids Res. 2025;53:D543–D553. doi:10.1093/nar/gkae1011.

