## Supplemental Material for "The K18-hACE2 mouse model of SARS-CoV-2 infection to illustrate the role and response of the vasculature in neurotropic viral infection"

**Supplemental Tables**

**Supplemental Table S1**. Study cohorts and investigations undertaken.

| **Cohorts** | **Virus** | **PFU/mouse** | **DPI: animal no (sex)** | **Investigations** |
| --- | --- | --- | --- | --- |
| 1.1 | Liverpool | 10^3^ | **6**: 1.1.1 - 1.1.6 (F) | H, IHa |
| 1.2 | Liverpool | 10^4^ | **4**: 1.2.1, 1.2.2 (M); 1.2.3 - 1.2.5 (F)  **6**: 1.2.6 - 1.2.9 (M); 1.2.10 - 1.2.13 (F) | H, IHa |
| 1.3 | Delta | 10^3^ | **5**: 1.3.1 (M); 1.3.2 (F)  **6**: 1.3.3, 1.3.4 (M); 1.3.5 - 1.3.11 (F)  **7**: 1.3.12, 1.3.13 (M); 1.3.14 - 1.3.17 (F) | H, IHa, IHb; RT-qPCR |
| 1.4 | Mock | - | **7**: 1.4.1, 1.4.2 (M); 1.4.3, 1.4.4 (F) | H, IHa; RT-qPCR |
| 2.1 | Delta | 10^2^ | **6**: 2.1.1 - 2.1.3 (F)  **7**: 2.1.4 - 2.1.8 (F) | H, IHa, IHb, IHc; RT-qPCR; multiomics; TEM, immune-EM |
| 2.2 | Delta | 10^3^ | **7**: 2.2.1 (F)  **8**: 2.2.2 - 2.2.5 (F) | H, IHa, IHb; TEM |
| 2.3 | Mock | - | **7**: 2.3.1 - 2.3.6 (F) | H, IHa, IHc; multiomics; TEM, immune-EM |

**Abbreviations**: Animal no: animal number for identification of individual mice; DPI: days post infection (day of euthanasia); Delta: SARS-CoV-2 Delta isolate; F: female; H: histological examination of the brain (HE-stained section); IHa: immunohistochemistry for SARS-CoV-2 nucleocapsid protein (NP), as previously described [1], performed on sections of the brain; IHb: immunohistochemistry for leukocyte markers (Iba1, CD3, CD4, CD8, Ly6G) and apoptotic cells (cleaved caspase 3), as previously described [1–3], performed on sections of the brain; IHc: immunohistochemistry for markers of blood vessels/blood-brain-barrier (PECAM-1/CD31, Claudin 5, PDGFR-β, α-SMA, AQP4), performed on sections of the brain; immune-EM: immune electron microscopy for SARS-CoV-2 nucleocapsid protein performed on the brain; Liverpool: Liverpool strain of SARS-CoV-2; PFU: plaque-forming units; M: male; Mock: mock-infected; multiomics: bulk transcriptomic, proteomic, metabolomic, and lipidomic performed on the brain; RT-qPCR: reverse transcriptase-quantitative PCR for SARS-CoV-2 N1 [4] performed on lung and brain tissue; TEM: transmission electron microscopy performed on the brain.

**Supplemental Table S2**. List of antibodies used for immunohistochemistry, including pretreatment and detection methods.

| **Antigen** | **Antibody (clone)** | **Dilution, incubation** | **Antigen retrieval** | **Detection system** |
| --- | --- | --- | --- | --- |
| SARS-CoV NP | Rb pAb^a^ | 1:6000, 4°C, ON | CB^b^ pH 6.0, 98°C, 20 min | EnVision+ (HRP, Rb)^b,1^ |
| CD3 | Rb mAb (SP7)^c^ | 1:350, 37°C, 1 h | CC1^d^ pH 9.0, 37° C, 20 min | DiscoveryOmniMap anti-Rb HRP, Discovery Chromo-Map^d,2^ |
| CD4 | Rb mAb^e^ | 1:170, 37°C, 1 h | CC1^d^ pH 9.0, 37° C, 20 min | Donkey anti-Rb Ab^f^;  Discovery DAB Map^d,2^ |
| CD8 (CD8α) | Rb mAb (D4W2Z)^g^ | 1:200, RT, 1 h | CB^b^ pH 6.0, 98°C, 20 min | EnVision+ (HRP, Rb)^b,1^ |
| Iba1 | Rb pAb^h^ | 1:1000, RT, 1 h | CB^b^ pH 6.0, 98°C, 20 min | EnVision+ (HRP, Rb)^b,1^ |
| Ly6G | Rat mAb (1A8)^i^ | 1:200, RT, 1 h | CB^b^ pH 6.0, 98°C, 20 min | Rb anti-rat Ab^j^;  EnVision+ (HRP, Rb)^b,1^ |
| Cleaved caspase 3 | Rb mAb (D175)^g^ | 1:200, 4°C, ON | TE^b^ pH 9.0, 98°C, 20 min | EnVision+ (HRP, Rb)^b,1^ |
| AQP4 | Rb pAb^k^ | 1:1000, 4°C, ON | None | EnVision+ (HRP, Rb)^b,1^ |
| Claudin 5 | Rb pAb^l^ | 1:400, 37°C, 1 h | CC1^d^ pH 9.0, 37° C, 20 min | DiscoveryOmniMap anti-Rb HRP, Discovery Chromo-Map^d,2^ |
| PDGFR-β | Rb mAb (28E1)^g^ | 1:20, 4°C, ON | TE^b^ pH 9.0, 98°C, 20 min | EnVision+ (HRP, Rb)^b,1^ |
| PECAM-1 (CD31) | Rb pAb^c^ | 1:1000, RT, 1 h | TE^b^ pH 9.0, 98°C, 20 min | EnVision+ (HRP, Rb)^b,1^ |
| α-SMA | Mouse mAb (1A4)^b^ | 1:400, RT, 1 h | None | MACH 4^d,1^ |

**Abbreviations:** AQP4: aquaporin 4; CB: citrate buffer; CC1: Cell Conditioning Solution; Iba1: ionized calcium binding adaptor molecule 1; Ly6G: lymphocyte antigen 6 family member G; mAb: monoclonal antibody; ON: overnight; pAb: polyclonal antibody; PDGFR-β: platelet-derived growth factor receptor beta; PECAM-1: platelet endothelial cell adhesion molecule 1; Rb: rabbit; RT: room temperature; α-SMA: alpha smooth muscle actin; TE: Tris-EDTA.

^1^Autostainer Link 48 (Agilent Dako), ^2^Discovery XT (Roche/Ventana, Basel, Switzerland)

**List of suppliers:**

^a^Rockland Immunochemicals Inc., Limerick, Pennsylvania, USA

^b^Agilent Dako, Santa Clara, California, USA

^c^Abcam, Cambridge, UK

^d^Roche/Ventana, Basel, Switzerland

^e^SinoBiological, Beijing, China

^f^Jackson ImmunoResearch Laboratories, Inc., Ely, UK

^g^Cell Signaling Technology, Danvers, Massachusetts, USA

^h^Wako Pure Chemical Corporation, Osaka, Japan

^i^BioLegend, San Diego, California, USA

^j^Vector Laboratories, Newark, California, USA

^k^Merck (Sigma-Aldrich), Burlington, Massachusetts, USA

^l^Invitrogen, ThermoFisher Scientific, Waltham, Massachusetts, USA

**Supplemental Table S3.** List of R packages used for the -omic analyses and the visualisation of the qRT-PCR results.

A) qRT-PCR results

| **Package** | **Version** | **References** |
| --- | --- | --- |
| ggplot2 | 3.5.1 | [5] |
| tidyr | 1.3.1 | [6] |
| dplyr | 1.1.4 | [7] |
| patchwork | 1.3.0 | [8] |

| **Package** | **Version** | **References** |
| --- | --- | --- |
| tximport | 1.32.0 | [9] |
| edgeR | 4.2.1 | [10] |
| PCAtools | 2.16.0 | [11] |
| EnhancedVolcano | 1.22.0 | [12] |
| clusterProfiler | 4.12.6 | [13–16] |
| fgsea | 1.30.0 | [17] |
| AnnotationDbi | 1.66.0 | [18] |
| tidyr | 1.3.1 | [6] |
| dplyr | 1.1.4 | [7] |
| forcats | 1.0.0 | [19] |
| stringr | 1.5.1 | [20] |
| ggplot2 | 3.5.1 | [5] |
| enrichplot | 1.24.4 | [21] |
| limma | 3.60.4 | [22] |
| org.Mm.eg.db | 3.19.1 | [23] |

B) Transcriptomic analysis

C) Proteomic analysis

| **Package** | **Version** | **References** |
| --- | --- | --- |
| DEP2 | 0.5.28.2 | [24, 25] |
| AnnotationDbi | 1.72.0 | [26] |
| ComplexHeatmap | 2.26.0 | [27, 28] |
| cowplot | 1.2.0 | [29] |
| dplyr | 1.1.4 | [7] |
| enrichplot | 1.30.4 | [30] |
| ggridges | 0.5.7 | [31] |
| ggplot2 | 4.0.1 | [5] |
| ggVennDiagram | 1.5.7 | [32] |
| limma | 3.66.0 | [22] |
| org.Mm.eg.db | 3.22.0 | [33] |
| tidyr | 1.3.2 | [34] |
| PCAtools | 2.22.1 | [35] |
| stringr | 1.6.0 | [36] |
| EnhancedVolcano | 1.28.2 | [37] |
| forcats | 1.0.1 | [38] |
| UpSetR | 1.4.0 | [39] |

D) Metabolomic analysis

| **Package** | **Version** | **References** |
| --- | --- | --- |
| tidymass | 2.0.10 | [40] |
| massdataset | 0.99.0 | [40] |
| masscleaner | 1.0.12 | [40] |
| massqc | 1.0.8 | [40] |
| massstat | 1.0.6 | [40] |
| ggplot2 | 4.0.1 | [5] |
| rstatix | 0.7.3 | [41] |
| EnhancedVolcano | 1.28.2 | [37] |
| ggrepel | 0.9.6 | [42] |
| dplyr | 1.1.4 | [7] |
| tidyr | 1.3.2 | [34] |
| stringr | 1.6.0 | [36] |
| patchwork | 1.3.2 | [8] |
| RefMet | 1.0.0 | [43] |
| pheatmap | 1.0.13 | [44] |

E) Lipidomic analysis

| **Package** | **Version** | **References** |
| --- | --- | --- |
| LipidSigR | 1.0.4 | [45–47] |
| rgoslin | 1.14.0 | [48, 49] |
| RefMet | 1.0.0 | [43] |
| SummarizedExperiment | 1.40.0 | [50] |
| PCAtools | 2.22.1 | [35] |
| dplyr | 1.1.4 | [7] |
| stringr | 1.6.0 | [36] |
| tidyr | 1.3.2 | [34] |
| EnhancedVolcano | 1.28.2 | [37] |
| ggplot2 | 4.0.1 | [5] |
| patchwork | 1.3.2 | [8] |
| rstatix | 0.7.3 | [41] |
| pheatmap | 1.0.13 | [44] |
| tibble | 3.3.1 | [51] |

**Supplemental Table S4**. This spreadsheet (separate file) shows the results of the SARS-CoV-2 N1 RT-qPCR, haematoxylin and eosin ((peri)vascular infiltrates) and immunohistochemistry (nucleocapsid protein pattern) examination in the brain.

**Supplemental Table S5.** Overview of the intersection of the transcriptome and proteome datasets with published datasets related to the murine blood-brain barrier (BBB), highlighting the significantly up- and down-regulated gene products (FDR < 0.05 and log_2_FC > |2| or > |0|).

| **Dataset**  (*type*) | **Size** | **Transcriptome** | | | **Proteome** | | |
| --- | --- | --- | --- | --- | --- | --- | --- |
|  |  | **Intersect** | **Up (> 2)** | **Down (< -2)** | **Intersect** | **Up (> 0)** | **Down (< 0)** |
| **Daneman et al., 2010 [52]**  (*transcriptome*) | 409^1^ | 293  (71.6%) | 8 | 3 | 122  (29.8%) | 1 | 4 |
| **Tremblay et al., 2024 [53]**  (*proteome*) | 224 | 122  (54.5%) | 8 | 1 | 68  (30.4%) | 6 | 1 |
| **Munji et al., 2019 – S3^2^ [54]**  (*transcriptome*) | 518 | 407  (78.6%) | 9 | 4 | 121  (23.4%) | 1 | 4 |
| **Munji et al., 2019 – S4^3^ [54]**  (*transcriptome*) | 70 | 64  (91.4%) | 0 | 0 | 32  (45.7%) | 0 | 2 |
| **Munji et al., 2019 – S6^4^ [54]**  (*transcriptome*) | 136 | 130  (95.6%) | 35 | 0 | 29  (21.3%) | 2 | 2 |

Gene products were considered as significant when the FDR was less than 0.05.

^1^	The original size of the dataset (as published) was 541; it dropped to 409 after removing duplicated entries and entries without gene symbol.

^2^	File S3: BBB-enriched genes.

^3^	File S4: tight junction proteins expressed in brain endothelial cells.

^4^	File S6: BBB dysfunction module.

**Supplemental Material & Methods**

**Transmission Electron Microscopy (TEM)**

The ultrathin sections were subjected to extensive screening of the blood vessels to determine any potential pathological changes, such as the presence of (peri)vascular leukocytes, changes at the level of the astrocytic endfeet (AEF), and morphological alterations of the blood vessel wall. The number of blood vessels chosen for further assessment and imaging ranged from 12 to 59 per brain (depending on the number of vessels in the sections). Images were taken from vessels primarily based on the presence of the following features: (peri)vascular leukocytes, evidence of AEF at section level, viral particles in cells adjacent to the vessels and/or changes suggestive of cell degeneration/death. In addition, blood vessels without apparent changes were included. For the subsequent in-depth assessment of the vessels, a two-step process was applied. First, the normal appearance of the blood vessels/BBB in the K18-hACE2 mice was determined, using literature as a reference [55–63]. Here, the vessels in the brains of the mock-infected control mice (2.3) served for the comparative assessment of vessels in the Delta-infected (cohorts 2.1 and 2.2) brains. In a second step, a qualitative review of all examined vessels was undertaken, focusing on the following morphological features: alteration of the vascular wall (cell degeneration/death, alteration of the endothelial tight junctions), changes at the level of the AEF (swelling, glycogen accumulation), presence of leukocytes (in the vascular wall, the perivascular space, or the adjacent neuroparenchyma), presence of viral particles (at any level of the vessel wall, in cells adjacent to the vessel).

A glutaraldehyde-fixed, epoxy resin embedded pellet of human embryonic kidney (HEK-293T) cells transduced to stably express the human angiotensin-converting enzyme 2 (ACE2) receptor, and infected with SARS-CoV-2 Omicron Xbb.1.5 at a dose of 5 MOI (multiplicity of infection, PFU/cell) and collected at 72 h post infection [64], served as positive control, i.e. as reference for viral structures at different stages of the viral cycle in cells in the brains (Supplemental Figure S3).

**Multiomic analyses**

***Transcriptomic*.** After import of the matrix of genes counts in RStudio (v2024.09.1 Build 394), R (v4.4.0 [65]) was used to perform all the transcriptomic analyses. The filtering of genes with low or no counts and the normalisation of the libraries size was done with the edgeR package (v4.2.1, [10]), using the filterByExpr() and calcNormFactors() functions. Visualisation and clustering of the samples was investigated with a principal component analysis (PCA), on log_2_-transformed counts per million (log_2_ CPM), using the PCAtools package (v2.16.0, [11]). Testing for differential gene expression was carried out using the exactTest() function from the edgeR package. Genes were considered as differentially expressed when showing at least a 4-fold difference (log_2_FC) from the mock-infected group, with a false discovery rate (FDR) below 0.05. Functional enrichment analysis was performed on a gene list sorted by decreasing values of log_2_FC (Gene Set Enrichment Analysis (GSEA)) with Gene Ontology (GO) [66, 67], using the gseGO() function from the clusterProfiler package (v4.12.6; [13–16]) (using fgsea algorithm; v1.30.0 [17]), with a Benjamini-Hochberg FDR threshold equaling 0.05. A list of the packages used is available in Supplemental Table S3B.

***Proteomic*.** Differential protein abundance analysis was undertaken in RStudio (v2026.01.0 build 392) with the R language (v4.5.2) [68], using DEP2 (v0.5.28.2 [24, 25]). The detected proteins were filtered to include only the host proteome (“*Mus musculus*”), after which missing values were filtered out (using a threshold of 30% of missing values per row). The intensities were log_2_-transformed and normalised using variance stabilising transformation. Imputation of the missing values was computed with the MinDet method, using DEP2::impute(). Visualisation and clustering of the samples was performed with a PCA, on log_2_-transformed intensities, using the PCAtools package (v2.22.1). Differential abundance was tested with DEP2::test_diff(), using limma [22] for the statistical testing and Benjamini-Hochberg multiple testing correction to control the FDR. Proteins with a FDR less than 0.05 were considered as significantly different. Functional enrichment analysis was performed with GSEA (DEP2::test_GSEA()), using GO [66, 67] with a Benjamini-Hochberg FDR threshold equaling 0.05. A list of the packages used is available in Supplemental Table S3C.

***Transcriptome-proteome integration***. First the overlap between the transcriptome and proteome spaces was investigated with Venn diagrammes, using ggVennDiagram package (v1.5.7, [32]), by considering all gene products, or only significantly up-regulated gene products, or only significantly down-regulated gene products. Second, the overall results of both analyses were compared with heatmaps, using the ComplexHeatmap R package (v2.26.0 [27, 28]). For this, using log_2-_transformed and row scaled counts and intensities, the 50 most variable transcripts and proteins were computed (using rowVars() from matrixStats v1.5.0 [69]) and were investigated in the transcriptome and proteome space. Third, for the functional analysis, the significant (FDR less than 0.05) GO [66, 67] enriched pathways were filtered on the BP ontology, and the top 10 (normalised enrichment score (NES) > 0) and bottom 10 (NES < 0) (based on the q-value) pathways from the transcriptome and proteome enrichment were extracted; using this list of GO ID, all corresponding significant pathways from the transcriptome and proteome were ordered based on the coalesced transcriptome and proteome NES (dplyr::coalesce()) and plotted using ggplot2 (v4.0.1).

In the subsequent analysis, we focused on pathways and specific genes that were of interest in the context of inflammation and the recruitment of leukocytes into the brain, and on those that would acknowledge differences in blood vessel and BBB function. For the significant pathways, the results of the GO GSEA analysis were subjected to a manual review and potential pathways of interest were marked. From this selection, pathways of the biological process (BP) ontology were extracted in RStudio and further classified into three categories, using manually curated lists of keywords, namely: 1) inflammatory mediators; 2) leukocytes; 3) blood vessels and BBB/neurovascular unit (NVU).

Genes of interest were derived from the literature and were organised in three categories, namely: 1) inflammatory mediators [70–76]; 2) leukocyte recruitment factors and activation [77–83]; 3) BBB/NVU key components/factors and genes known to be associated with BBB dysfunction [79, 84–97]. Using these gene lists, the differentially expressed gene products (inflammatory mediators, leukocyte recruitment factors and activity), or all the gene products (BBB/NVU key components/factors, genes associated with alterations of BBB function), were parsed, and the corresponding log_2_ transformed and row scaled counts and intensities, were visualised using ComplexHeatmap for gene products of interest. The BBB and NVU were further investigated using published results of the murine BBB proteome and transcriptome [52–54]; the results were also visualised using ComplexHeatmap. A list of the packages used is available in Supplemental Table S3C.

***Metabolomic***. The manually curated table of compound intensities (n = 130) was imported in RStudio (v2026.01.0 build 392) for further analysis with the R language (v4.5.2) [68], using the TidyMass pipeline (v2.0.10, [40]), and following best practices [98]. For the analysis, the information about the samples and the compound intensities were combined using the massdataset (v0.99.0, [40]) create_mass_dataset() function. Data exploration was performed, before and after normalisation on the median, via density plots (ggplot2 and rstatix (get_summary_stats() function) v0.7.3, [41]), and boxplots (massqc (v1.0.8, [40]) massqc_sample_boxplot() function; ggplot2). The data were normalised to the median using the masscleaner (v1.0.12, [40]) normalize_data() function. Data exploration was performed on log_2_ transformed data with PCA, using PCAtools (v2.22.1), and heatmap (data were row scaled using the scale() function), using pheatmap (v1.0.13, [44]). Normality of the data was investigated with the Shapiro-Wilk test implemented in rstatix, using the shapiro_test() function; this was done on normalised data, with and without log_10_ transformation. For the univariate analysis, as for both cohorts there were metabolite intensities which were not following a normal distribution even after log_10_ transformation, the statistical analysis was conducted on the median normalised non-log-transformed data ; the computation of the fold change was done by comparing the medians, and the p-value was computed using a non-parametric test (Wilcoxon rank sum test/Mann-Whitney U test) (massstat (v1.0.6, [40]); mutate_fc() and mutate_p_value respectively). The correction for multiple testing was done using the Benjamini-Hochberg method; the metabolites for which the adjusted p-value was <0.05 were considered as significantly different. The results of the univariate analysis were visualised with a volcano plot, using EnhancedVolcano (v1.28.2), and the significantly different metabolites were plotted with their standardised name, using RefMet (v1.0.0 [43]) for the mapping (as Prostaglandin F2 could not be successfully mapped it was plotted without a standard name). A list of the packages used is available in Supplemental Table S3D.

***Lipidomic***. The manually curated tables of compound intensities (positive (n = 963) and negative (n = 53) modes) were imported in RStudio (v2026.01.0 build 392) for further analysis with the R language (v4.5.2) [68], using the LipidSigR pipeline (v1.0.4, [45–47]), and following best practices [98]. The detected lipids were mapped using rgoslin (v1.14.0, [48, 49]) as well as RefMet (v1.0.0) to ensure the use of standardised names. When isomers or duplicates were present, the lipid intensities were summed for each sample, so that only one lipid entry for that name remained (aggregate() function from the stats package). Ultimately, 784 lipids from the positive mode and 27 lipids from the negative mode were available for analysis; all subsequent analyses were performed on each lipid dataset (positive and negative mode) separately. Lipid intensities were normalised to the median using LipidSigR data_process() function. Data exploration was performed on log_2_-transformed data with PCA, using PCAtools (v2.22.1), and heatmaps (data were row scaled using the scale() function), using pheatmap (v1.0.13). Normality of the data was investigated with the Shapiro-Wilk test implemented in rstatix, using the shapiro_test() function; this was done on normalised data, with and without log_10_ transformation. As several lipid intensities in both cohorts did not follow a normal distribution, the differential analysis was performed with a non-parametric test. This was done using the LipidSigR deSp_twoGroup() function, with a Wilcoxon test, and the correction for multiple testing was done using the Benjamini-Hochberg method; the fold change was computed as the ratio of the means. The lipids for which the adjusted p-value was <0.05 were considered as significantly different. The results of the differential analysis were visualised with a volcano plot, using EnhancedVolcano (v1.28.2). In addition to the statistical testing for individual lipid species, differential analyses were conducted for two lipid characteristics on the positive mode dataset (as no significant lipids were detected from the negative datasets), using the lipid_profiling() function from LipidSigR, namely: class and function. A list of the packages used is available in Supplemental Table S3E.

**Supplemental Figures**

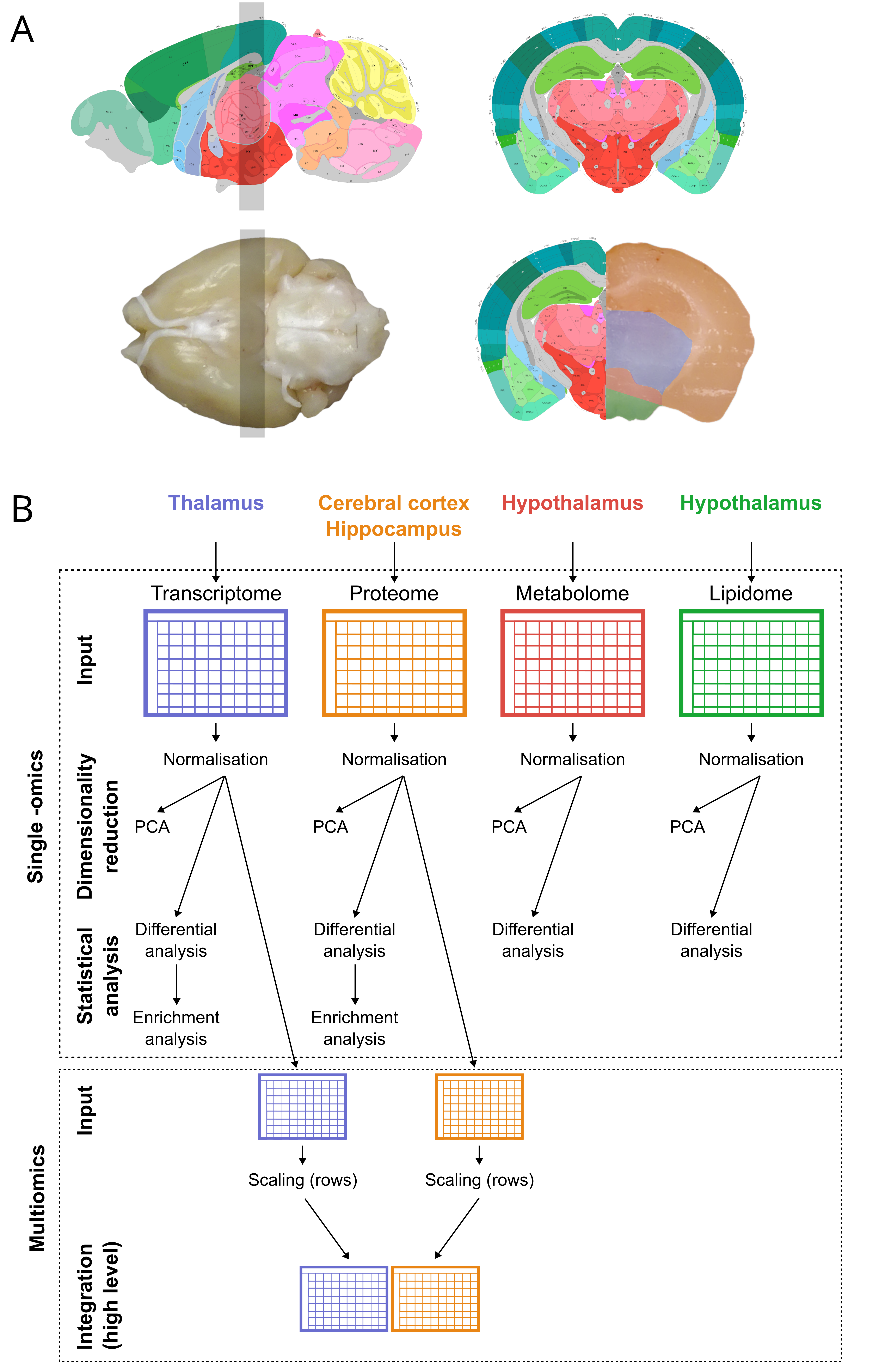

**Supplemental Figure S1**. Overview of the multiomics pipeline. **A)** Illustration of the tissue slice prepared by coronal sections (grey rectangle) to provide the samples for transmission electron microscopy and for the bulk transcriptomic, proteomic, metabolomic, and lipidomic analyses. Pictures of an example brain are mapped to Allen Mouse Brain Atlas (Allen Reference Atlas – Adult Mouse Coronal Sections (image 75), Mouse, P56 Sagittal (image 21) - <https://atlas.brain-map.org/>) [99–102]. **B)** Overview of the workflow used for the single -omics and the multiomics analyses. PCA: principal component analysis.

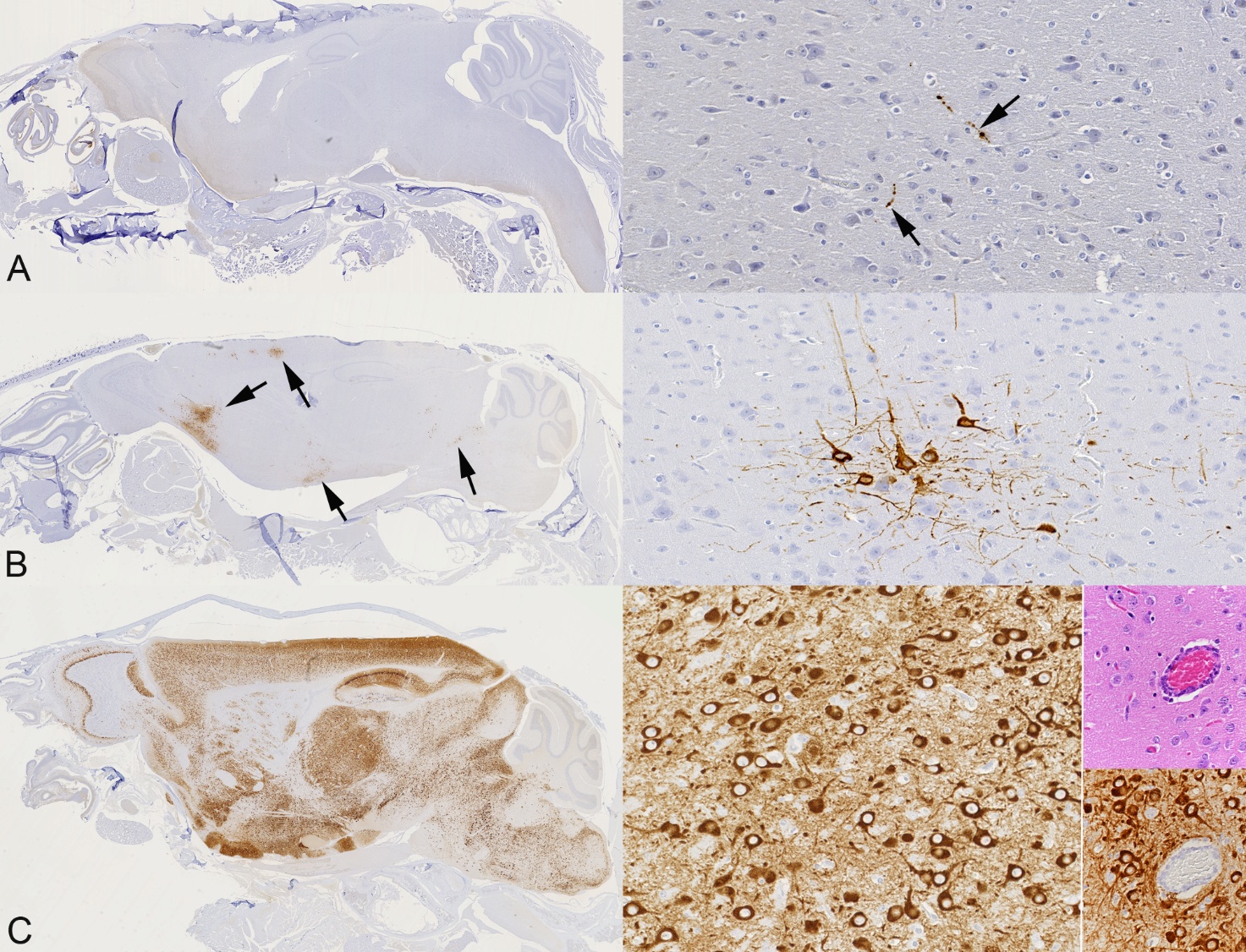

**Supplemental Figure S2. SARS-CoV-2 nucleocapsid protein (NP) expression and distribution pattern in the brains of K18-hACE2 mice after intranasal challenge with SARS-CoV-2 Liverpool strain or Delta.** Left column: overview of a midline longitudinal section of the brain; right column: illustration of positive neurons at higher magnification. **A)** ”Scattered” pattern, represented by a few individual cells or cell processes expressing viral antigen (arrows). Mouse (animal 1.2.13) infected with SARS-CoV-2 Liverpool strain (10^4^ PFU), 6 dpi. **B)** ”Clustered” pattern, represented by groups of positive neurons (arrows) with staining in cell body and processes. Mouse (animal 1.1.6) infected with SARS-CoV-2 Liverpool strain (10^3^ PFU), 6 dpi. **C)** ”Widespread” pattern, represented by large, coalescing patches of positive neurons. Mouse (animal 1.3.10) infected with SARS-CoV-2 Delta (10^3^ PFU), 6 dpi. The images on the far right show a (peri)vascular infiltrate (top: HE stain) in the brain of a mouse with widespread neuronal infection (animal 2.1.5) infected with SARS-CoV-2 Delta (10^2^ PFU), 7 dpi. The vascular structures are spared from viral antigen expression (bottom: IHC for viral NP). Immunohistochemistry (IHC), haematoxylin counterstain.

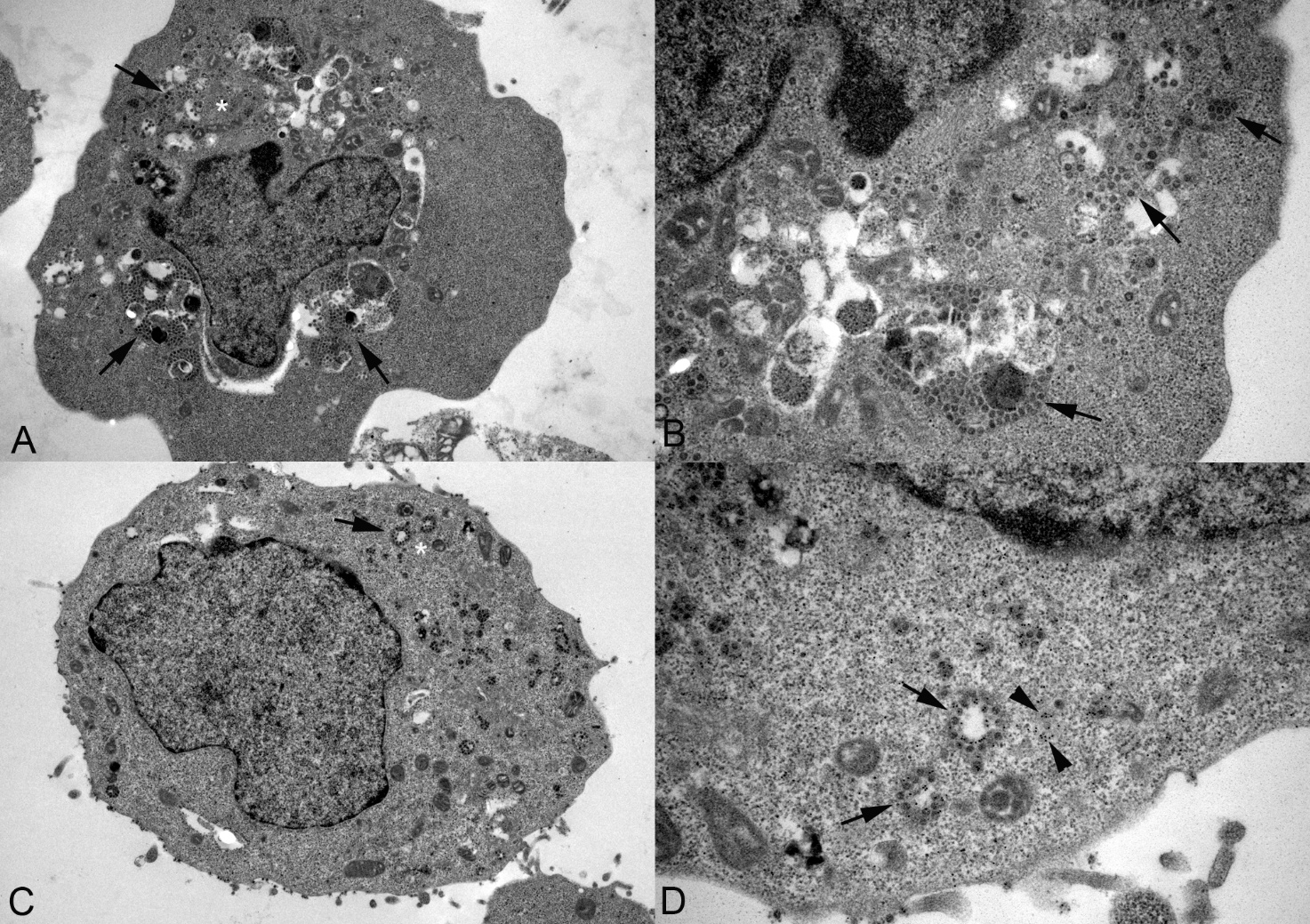

**Supplemental Figure S3. Ultrastructural evidence of SARS-CoV-2 infection in HEK-293T cells; infected with Omicron Xbb.1.5 at a dose of 5 MOI (PFU/cell) for 72 hours. A, B)** Infected cell with viral particles in the cytoplasm (arrows). The white asterisk in A highlights the area shown at higher magnification in B. **C, D)** Infected cell after immunogold labelling of viral nucleocapsid protein (NP). Viral particles are visible in the cytoplasm (arrows). The white asterisk in C highlights the area shown at higher magnification in D. Immunogold deposition (i.e. viral NP) is observed both associated with viral particles (arrows) and free in the cytoplasm (arrowheads). Transmission electron microscopy including immune-EM for SARS-CoV-2 NP (C, D).

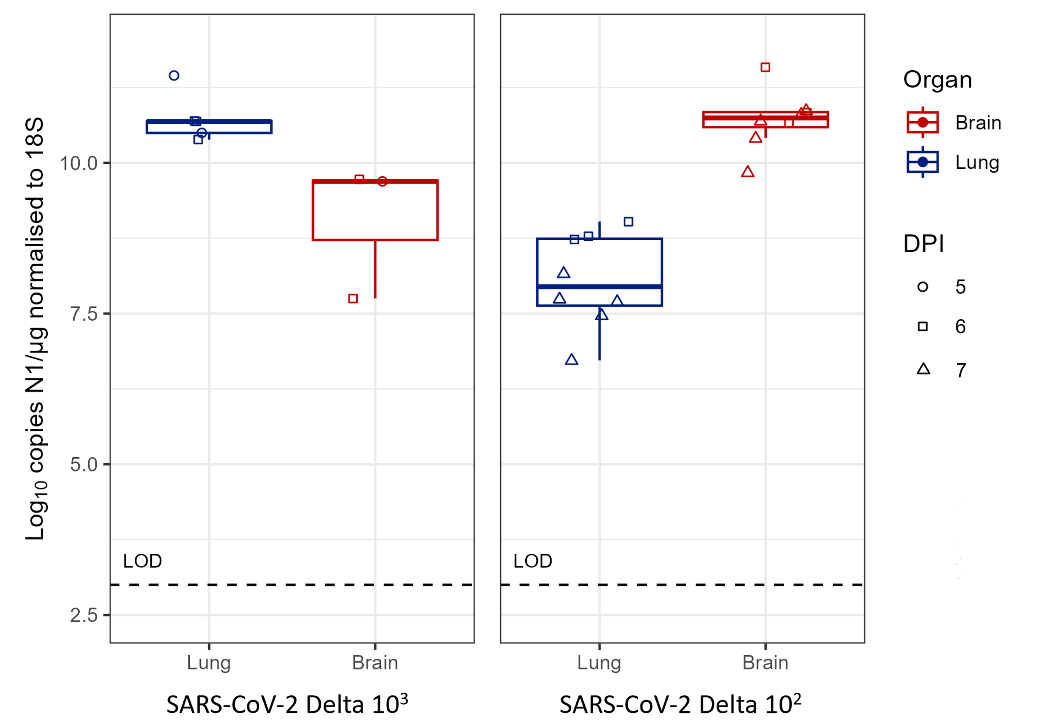

**Supplemental Figure S4. Viral RNA loads in lung and brain of K18-hACE2 mice after intranasal challenge with SARS-CoV-2 Delta at 10^2^ or 10^3^ PFU/mouse.** SARS-CoV-2 N1 RNA was detected by RT-qPCR in lungs and brains. With infection at 10^3^ PFU/mouse, viral RNA levels in the brain are lower than with infection at 10^2^ PFU/mouse at 6 and 7 dpi (cohort 2.1) whereas they are higher in the lungs. DPI: days post infection; LOD: limit of detection.

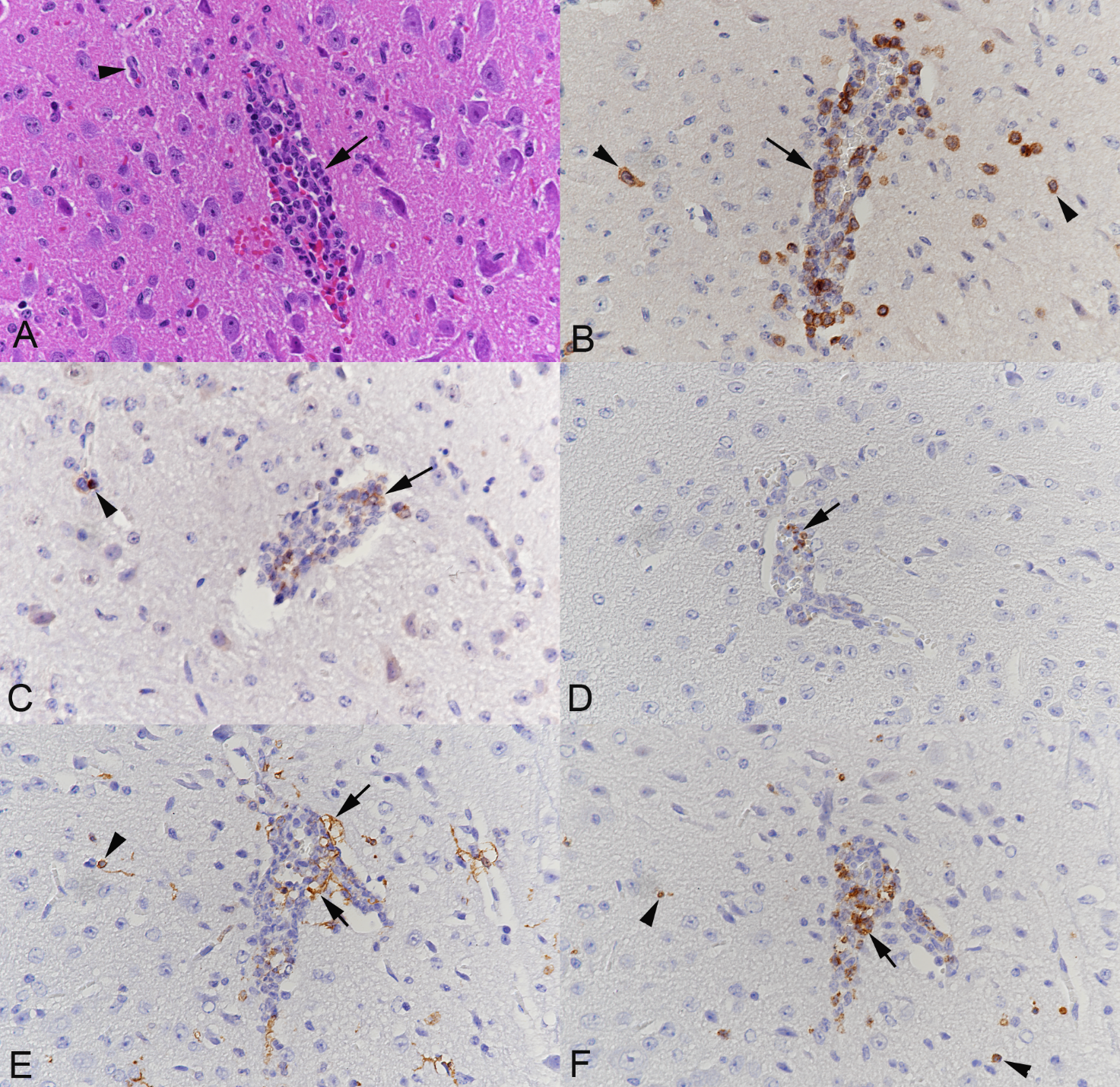

**Supplemental Figure S5. Composition of the (peri)vascular infiltrate in the brain of a K18-hACE2 mouse infected with SARS-CoV-2 Delta.** The mouse was inoculated intranasally with 10^2^ PFU and culled at 7 dpi (animal 2.1.4); it showed widespread viral antigen expression in the brain. **A)** HE stained section showing a blood vessel with a prominent (peri)vascular infiltrate (arrow) and a small unaffected capillary in the vicinity (arrowhead). **B)** The (peri)vascular infiltrate comprises numerous T cells (CD3+) (arrow); individual T cells are also found in the adjacent neuroparenchyma (arrowheads). **C)** A proportion of cells in the (peri)vascular infiltrate are CD4-positive (arrow). Individual CD4-positive T cells are also found in the adjacent neuroparenchyma (arrowhead). **D)** A proportion of cells in the (peri)vascular infiltrate are CD8-positive (arrow). **E)** Monocytes/macrophages (Iba1+) are numerous in the (peri)vascular infiltrate (arrows); individual macrophages (arrowhead) are also present in the adjacent neuroparenchyma. **F)** The (peri)vascular infiltrate also comprises neutrophilic leukocytes (Ly6G+) (arrow). Individual neutrophilic leukocytes are also found in the adjacent neuroparenchyma (arrowheads). Immunohistochemistry, haematoxylin counterstain.

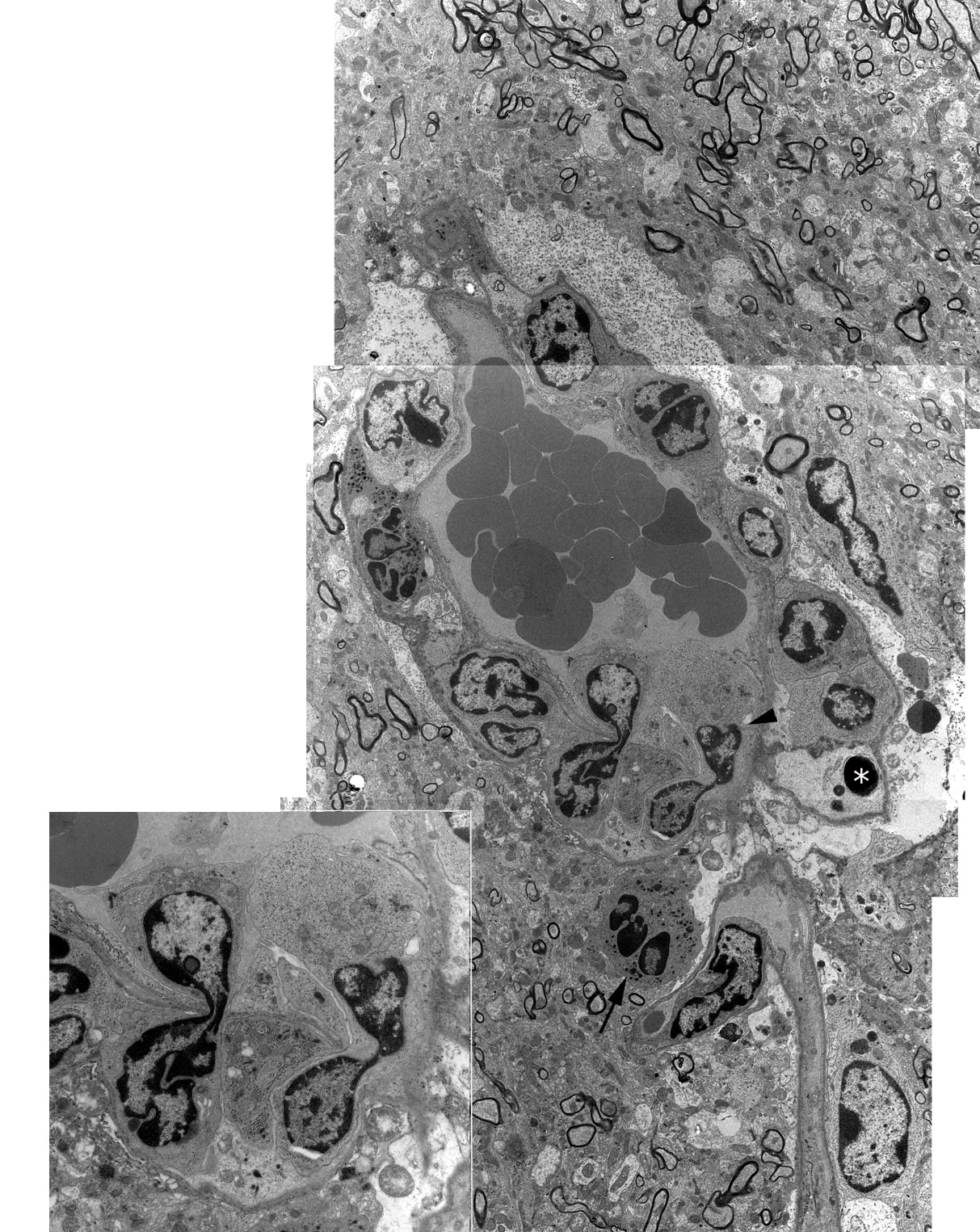

**Supplemental Figure S6. The (peri)vascular infiltrates are a consequence of leukocyte emigration without damage to the vascular wall.** Brainstem, mouse infected with 10^2^ PFU SARS-CoV-2 Delta (cohort 2.1; animal 2.1.4) and euthanised at 7 dpi. The figure is a manual montage to show the complete blood vessel and immediate surrounding illustrated in Figure 2C. It shows a second emigrating leukocyte (black arrowhead), a neutrophilic leukocyte in the neuroparenchyma (black arrow), and an apoptotic body (white asterisk) in the perivascular space. Inset (left): higher magnification of the two emigrating leukocytes.

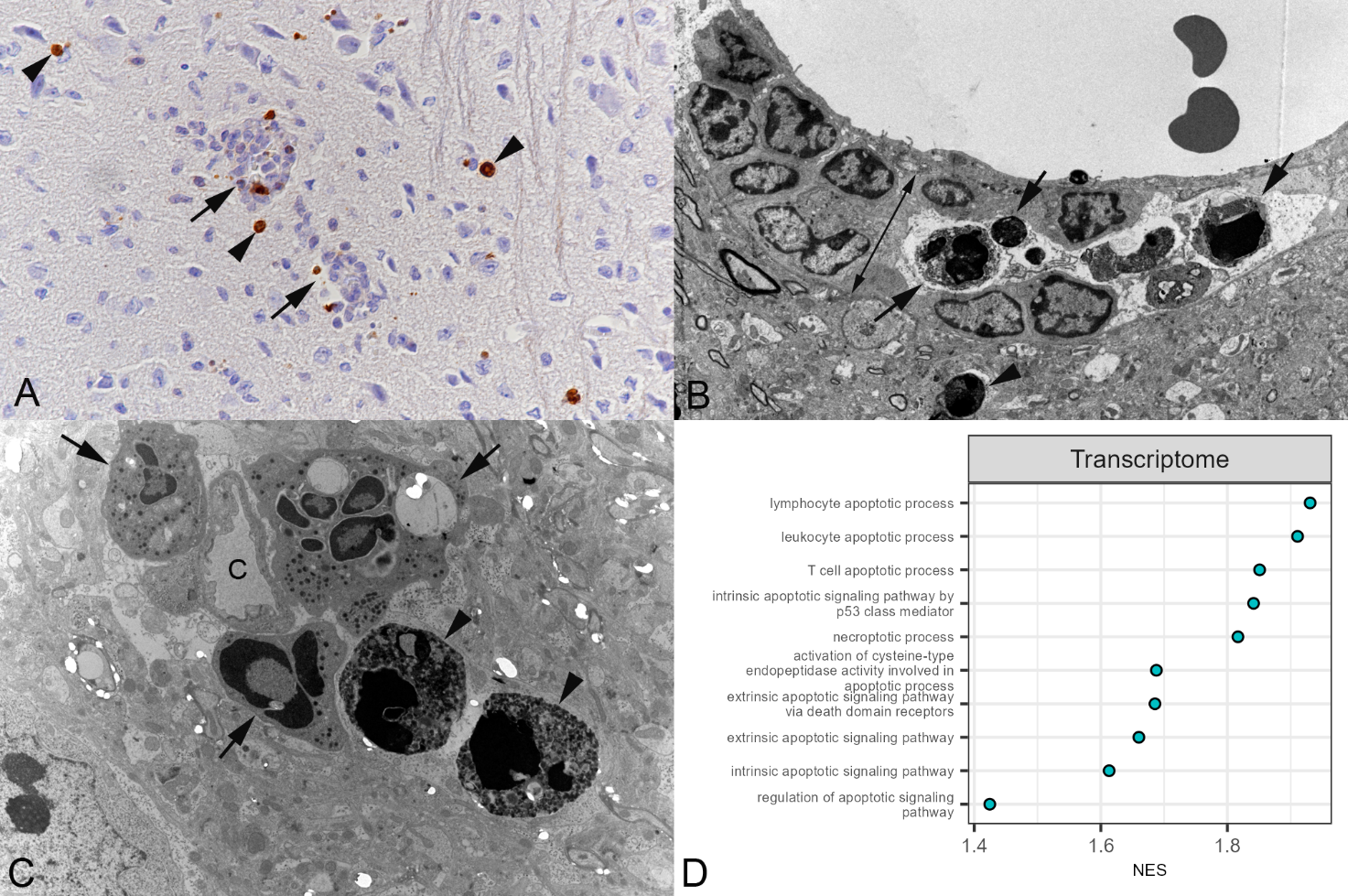

**Supplemental Figure S7. Apoptotic cell death of leukocytes in (peri)vascular infiltrates and adjacent neuroparenchyma.** Mice infected with SARS-CoV-2 Delta at 10^2^ PFU/mouse and euthanized at 6 or 7 dpi (cohort 2.1) and mock-infected mice (cohort 2.3). **A)** Animal 2.1.4. Staining for cleaved caspase 3 identifies individual apoptotic leukocytes in the (peri)vascular infiltrate (arrows) and in the adjacent neuroparenchyma (arrowheads).  Immunohistochemistry, haematoxylin counterstain. **B-C)** Transmission electron microscopy. **B)** Animal 2.1.1. Postcapillary venule; perivascular space (highlighted by the arrow with two tips) expanded by leukocytes among which are two apoptotic leukocytes and some apoptotic bodies (arrows). A further apoptotic cell is present in the adjacent neuroparenchyma (arrowhead). **C)** Animal 2.1.4. Three neutrophilic leukocytes (arrows) and two, possibly phagocytosed, apoptotic cells (arrowheads) in the neuroparenchyma adjacent to a capillary (C). **D)** Illustration of the top 10 GSEA-enriched pathways for biological process from Gene Ontology (comparing cohorts 2.1 and 2.3), sorted on q-value, associated with (leukocyte) cell death. NES: normalised enrichment score.

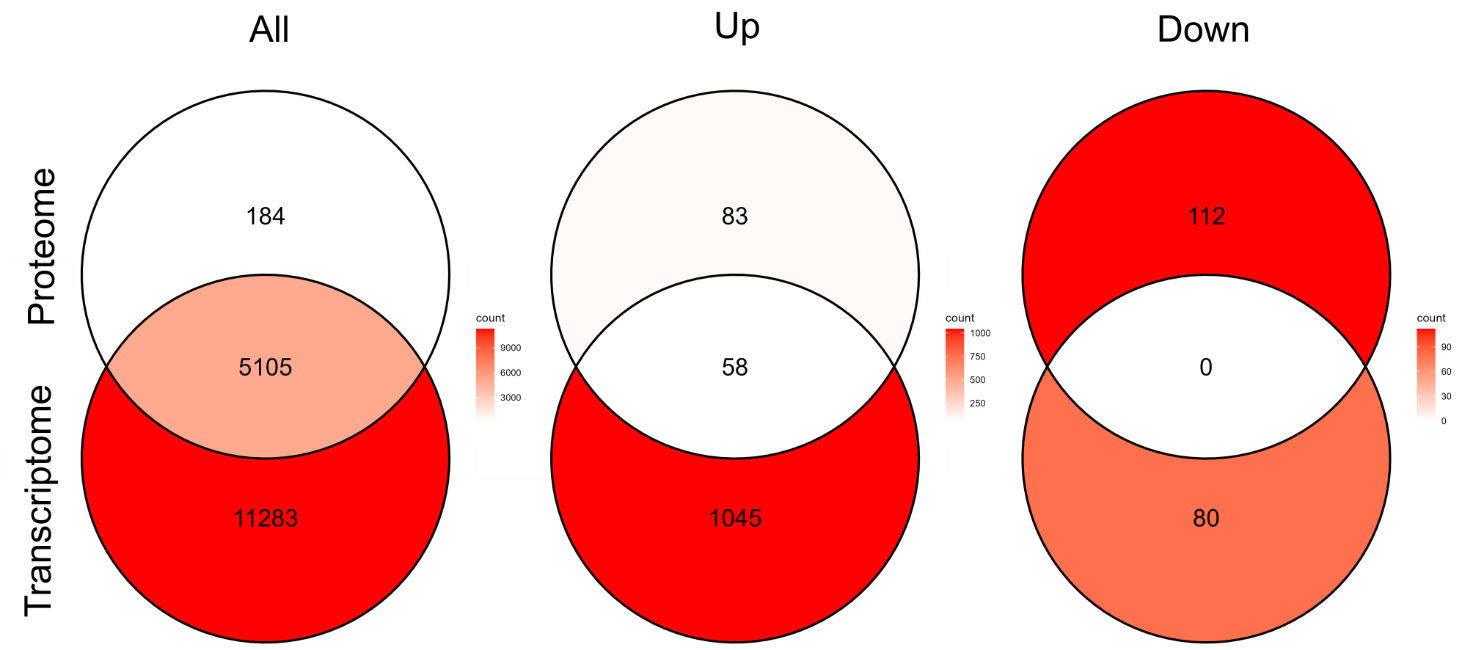

**Supplemental Figure S8.** **Results of the bulk transcriptomic and proteomic analyses in SARS-CoV-2 Delta-infected brains.** Mice infected with SARS-CoV-2 Delta at 10^2^ PFU/mouse (cohort 2.1) and mock-infected mice (cohort 2.3). Intersection of the transcriptome and proteome for all transcripts and proteins (All), the significantly up-regulated ones (Up) (log_2_FC > 2 (transcript); log_2_FC > 0 (protein), FDR < 0.05), and the significantly downregulated ones (Down) (log_2_FC < -2 (transcript); log_2_FC < 0 (protein), FDR < 0.05).

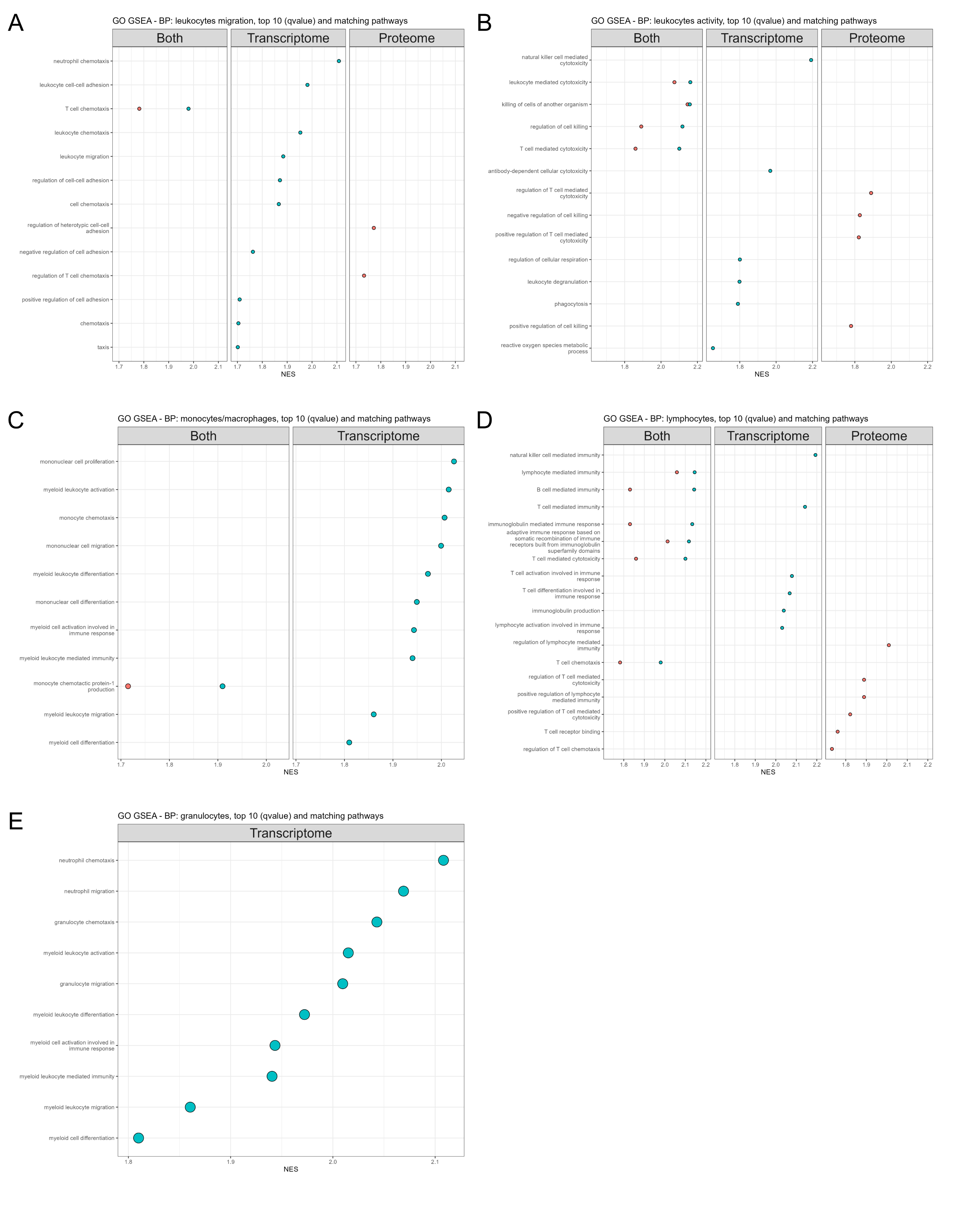

**Supplemental Figure S9. GO GSEA pathway enrichment analysis** **in SARS-CoV-2 Delta-infected brains based on leukocyte keywords.** Mice infected with SARS-CoV-2 Delta at 10^2^ PFU/mouse (cohort 2.1) and mock-infected mice (cohort 2.3). Illustration of the top 10 enriched pathways (GO GSEA) (based on q-value) of the Biological Process (BP) ontology for the transcriptome and proteome, with matching pathways, sorted on the coalesced transcriptome and proteome normalised enrichment score (NES). **A)** Leukocyte migration. **B)** Leukocyte activity. **C)** Monocytes and macrophages. **D)** Lymphocytes. **E)** Granulocytes.

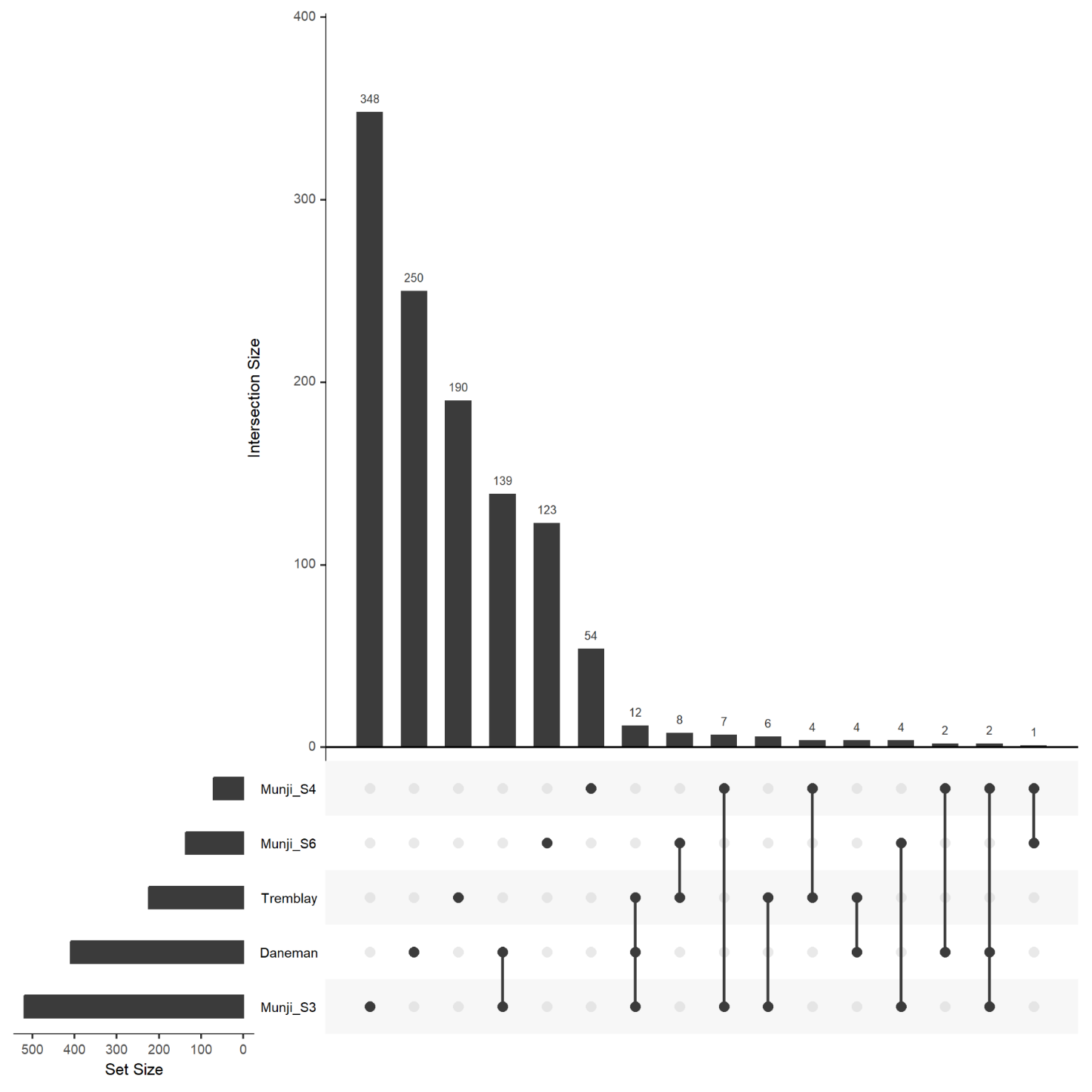

**Supplemental Figure S10. Comparisons of the public transcriptome and proteome datasets used to investigate the blood-brain barrier (BBB)/neurovascular unit. This Upset plot shows the intersection between the different datasets.** Tremblay: proteome of the BBB (Tremblay et al., 2024); Daneman: transcriptome of the BBB (Daneman et al., 2010); Munji_S3: transcriptome of BBB-enriched genes (Munji et al., 2019); Munji_S4: transcriptome of tight junction proteins expressed in brain endothelial cells (Munji et al., 2019); Munji_S6: BBB dysfunction module (Munji et al., 2019).

References

1.	Seehusen F, Clark JJ, Sharma P, Bentley EG, Kirby A, Subramaniam K, et al. Neuroinvasion and neurotropism by SARS-CoV-2 variants in the K18-hACE2 mouse. Viruses 2022. doi:10.3390/v14051020.

2.	Cabrera LE, Jokiranta ST, Mäki S, Miettinen S, Kant R, Kareinen L, et al. The assembly of neutrophil inflammasomes during COVID-19 is mediated by type I interferons. PLoS Pathog. 2024;20:e1012368. doi:10.1371/journal.ppat.1012368.

3.	Kang Y, Hepojoki J, Maldonado RS, Mito T, Terzioglu M, Manninen T, et al. Ancestral allele of DNA polymerase gamma modifies antiviral tolerance. Nature. 2024;628:844–53. doi:10.1038/s41586-024-07260-z.

4.	Clark JJ, Penrice-Randal R, Sharma P, Dong X, Pennington SH, Marriott AE, et al. Sequential infection with Influenza A virus followed by Severe Acute Respiratory Syndrome Coronavirus 2 (SARS-CoV-2) leads to more severe disease and encephalitis in a mouse model of COVID-19. Viruses 2024. doi:10.3390/v16060863.

5.	Wickham H. ggplot2: Elegant Graphics for Data Analysis. 2nd ed. Cham: Springer International Publishing; Imprint: Springer; 2016.

6.	Wickham H, Vaughan D, Girlich M. tidyr: Tidy Messy Data; 2024.

7.	Wickham H, François R, Henry L, Müller K, Vaughan D. dplyr: A Grammar of Data Manipulation; 2025.

8.	Pedersen TL. patchwork: The Composer of Plots; 2025.

9.	Soneson C, Love MI, Robinson MD. Differential analyses for RNA-seq: transcript-level estimates improve gene-level inferences. F1000Res. 2015;4:1521. doi:10.12688/f1000research.7563.2.

10.	Chen Y, Chen L, Lun ATL, Baldoni PL, Smyth GK. edgeR v4: powerful differential analysis of sequencing data with expanded functionality and improved support for small counts and larger datasets. Nucleic Acids Res 2025. doi:10.1093/nar/gkaf018.

11.	Blighe Kevin, Lun Aaron. PCAtools: PCAtools: Everything Principal Components Analysis; 2024.

12.	Blighe K. EnhancedVolcano: Publication-ready volcano plots with enhanced colouring and labeling: Bioconductor; 2024.

13.	Yu G. Thirteen years of clusterProfiler. Innovation (Camb). 2024;5:100722. doi:10.1016/j.xinn.2024.100722.

14.	Xu S, Hu E, Cai Y, Xie Z, Luo X, Zhan L, et al. Using clusterProfiler to characterize multiomics data. Nat Protoc. 2024;19:3292–320. doi:10.1038/s41596-024-01020-z.

15.	Wu T, Hu E, Xu S, Chen M, Guo P, Dai Z, et al. clusterProfiler 4.0: A universal enrichment tool for interpreting omics data. Innovation (Camb). 2021;2:100141. doi:10.1016/j.xinn.2021.100141.

16.	Yu G, Wang L-G, Han Y, He Q-Y. clusterProfiler: an R package for comparing biological themes among gene clusters. OMICS. 2012;16:284–7. doi:10.1089/omi.2011.0118.

17.	Korotkevich G, Sukhov V, Budin N, Shpak B, Artyomov MN, Sergushichev A. Fast gene set enrichment analysis. bioRxiv 2016. doi:10.1101/060012.

18.	Hervé Pagès, Marc Carlson, Seth Falcon, Nianhua Li. AnnotationDbi: Bioconductor; 2024.

19.	Wickham H. forcats: Tools for Working with Categorical Variables (Factors); 2023.

20.	Wickham H. stringr: Simple, Consistent Wrappers for Common String Operations; 2023.

21.	Guangchuang Yu. enrichplot: Bioconductor; 2024.

22.	Ritchie ME, Phipson B, Di Wu, Hu Y, Law CW, Shi W, Smyth GK. limma powers differential expression analyses for RNA-sequencing and microarray studies. Nucleic Acids Res. 2015;43:e47. doi:10.1093/nar/gkv007.

23.	Marc Carlson. org.Mm.eg.db: Bioconductor; 2024.

24.	Feng Z, Fang P, Zheng H, Zhang X. DEP2: an upgraded comprehensive analysis toolkit for quantitative proteomics data. Bioinformatics 2023. doi:10.1093/bioinformatics/btad526.

25.	Zhang X, Smits AH, van Tilburg GB, Ovaa H, Huber W, Vermeulen M. Proteome-wide identification of ubiquitin interactions using UbIA-MS. Nat Protoc. 2018;13:530–50. doi:10.1038/nprot.2017.147.

26.	Hervé Pagès, Marc Carlson, Seth Falcon, Nianhua Li. AnnotationDbi: Bioconductor; 2025.

27.	Gu Z. Complex heatmap visualization. Imeta. 2022;1:e43. doi:10.1002/imt2.43.

28.	Gu Z, Eils R, Schlesner M. Complex heatmaps reveal patterns and correlations in multidimensional genomic data. Bioinformatics. 2016;32:2847–9. doi:10.1093/bioinformatics/btw313.

29.	Wilke CO. cowplot: Streamlined Plot Theme and Plot Annotations for 'ggplot2'; 2025.

30.	Guangchuang Yu. enrichplot: Bioconductor; 2025.

31.	Wilke CO. ggridges: Ridgeline Plots in 'ggplot2'; 2025.

32.	Gao C-H, Dusa A. ggVennDiagram: A 'ggplot2' Implement of Venn Diagram; 2026.

33.	Marc Carlson. org.Mm.eg.db: Bioconductor; 2026.

34.	Wickham H, Vaughan D, Girlich M. tidyr: Tidy Messy Data; 2025.

35.	Blighe Kevin, Lun Aaron. PCAtools: PCAtools: Everything Principal Components Analysis; 2026.

36.	Wickham H. stringr: Simple, Consistent Wrappers for Common String Operations; 2025.

37.	Blighe K. EnhancedVolcano: Publication-ready volcano plots with enhanced colouring and labeling: Bioconductor; 2025.

38.	Wickham H. forcats: Tools for Working with Categorical Variables (Factors); 2025.

39.	Lex A, Gehlenborg N, Strobelt H, Vuillemot R, Pfister H. UpSet: Visualization of Intersecting Sets. IEEE Trans Vis Comput Graph. 2014;20:1983–92. doi:10.1109/TVCG.2014.2346248.

40.	Shen X, Yan H, Wang C, Gao P, Johnson CH, Snyder MP. TidyMass an object-oriented reproducible analysis framework for LC-MS data. Nat Commun. 2022;13:4365. doi:10.1038/s41467-022-32155-w.

41.	Kassambara A. rstatix: Pipe-Friendly Framework for Basic Statistical Tests; 2025.

42.	Slowikowski K. ggrepel: Automatically Position Non-Overlapping Text Labels with 'ggplot2'; 2024.

43.	Fahy E, Subramaniam S. RefMet: a reference nomenclature for metabolomics. Nat Methods. 2020;17:1173–4. doi:10.1038/s41592-020-01009-y.

44.	Kolde Raivo. pheatmap: Pretty heatmaps; 2025.

45.	Lin W-J, Shen P-C, Liu H-C, Cho Y-C, Hsu M-K, Lin I-C, et al. LipidSig: a web-based tool for lipidomic data analysis. Nucleic Acids Res. 2021;49:W336-W345. doi:10.1093/nar/gkab419.

46.	Liu C-H, Shen P-C, Lin W-J, Liu H-C, Tsai M-H, Huang T-Y, et al. LipidSig 2.0: integrating lipid characteristic insights into advanced lipidomics data analysis. Nucleic Acids Res. 2024;52:W390-W397. doi:10.1093/nar/gkae335.

47.	Liu C-H, Shen P-C, Tsai M-H, Liu H-C, Lin W-J, Lai Y-L, et al. LipidSigR: a R-based solution for integrated lipidomics data analysis and visualization. Bioinform Adv. 2025;5:vbaf047. doi:10.1093/bioadv/vbaf047.

48.	Kopczynski D, Hoffmann N, Peng B, Ahrends R. Goslin: A Grammar of Succinct Lipid Nomenclature. Anal Chem. 2020;92:10957–60. doi:10.1021/acs.analchem.0c01690.

49.	Kopczynski D, Hoffmann N, Peng B, Liebisch G, Spener F, Ahrends R. Goslin 2.0 Implements the Recent Lipid Shorthand Nomenclature for MS-Derived Lipid Structures. Anal Chem. 2022;94:6097–101. doi:10.1021/acs.analchem.1c05430.

50.	Martin Morgan, Valerie Obenchain, Jim Hester, Hervé Pagès. SummarizedExperiment: Bioconductor; 2025.

51.	Müller K, Wickham H. tibble: Simple Data Frames; 2026.

52.	Daneman R, Zhou L, Agalliu D, Cahoy JD, Kaushal A, Barres BA. The mouse blood-brain barrier transcriptome: a new resource for understanding the development and function of brain endothelial cells. PLoS One. 2010;5:e13741. doi:10.1371/journal.pone.0013741.

53.	Tremblay T-L, Alata W, Slinn J, Baumann E, Delaney CE, Moreno M, et al. The proteome of the blood-brain barrier in rat and mouse: highly specific identification of proteins on the luminal surface of brain microvessels by in vivo glycocapture. Fluids Barriers CNS. 2024;21:23. doi:10.1186/s12987-024-00523-x.

54.	Munji RN, Soung AL, Weiner GA, Sohet F, Semple BD, Trivedi A, et al. Profiling the mouse brain endothelial transcriptome in health and disease models reveals a core blood-brain barrier dysfunction module. Nat Neurosci. 2019;22:1892–902. doi:10.1038/s41593-019-0497-x.

55.	Nahirney PC, Tremblay M-E. Brain ultrastructure: putting the pieces together. Front Cell Dev Biol. 2021;9:629503. doi:10.3389/fcell.2021.629503.

56.	Krasemann S, Dittmayer C, Stillfried S von, Meinhardt J, Heinrich F, Hartmann K, et al. Assessing and improving the validity of COVID-19 autopsy studies - A multicentre approach to establish essential standards for immunohistochemical and ultrastructural analyses. EBioMedicine. 2022;83:104193. doi:10.1016/j.ebiom.2022.104193.

57.	Nahirney PC, Reeson P, Brown CE. Ultrastructural analysis of blood-brain barrier breakdown in the peri-infarct zone in young adult and aged mice. J Cereb Blood Flow Metab. 2016;36:413–25. doi:10.1177/0271678X15608396.

58.	Haley MJ, Lawrence CB. The blood-brain barrier after stroke: Structural studies and the role of transcytotic vesicles. J Cereb Blood Flow Metab. 2017;37:456–70. doi:10.1177/0271678X16629976.

59.	Kajihara H, Tsutsumi E, Kinoshita A, Nakano J, Takagi K, Takeo S. Activated astrocytes with glycogen accumulation in ischemic penumbra during the early stage of brain infarction: immunohistochemical and electron microscopic studies. Brain Res. 2001;909:92–101. doi:10.1016/s0006-8993(01)02640-3.

60.	Sá-Pereira I, Brites D, Brito MA. Neurovascular unit: a focus on pericytes. Mol Neurobiol. 2012;45:327–47. doi:10.1007/s12035-012-8244-2.

61.	Díaz-Castro B, Robel S, Mishra A. Astrocyte endfeet in brain function and pathology: Open questions. Annu Rev Neurosci. 2023;46:101–21. doi:10.1146/annurev-neuro-091922-031205.

62.	Hartmann DA, Coelho-Santos V, Shih AY. Pericyte control of blood flow across microvascular zones in the central nervous system. Annu Rev Physiol. 2022;84:331–54. doi:10.1146/annurev-physiol-061121-040127.

63.	Gouveia-Freitas K, Bastos-Leite AJ. Perivascular spaces and brain waste clearance systems: relevance for neurodegenerative and cerebrovascular pathology. Neuroradiology. 2021;63:1581–97. doi:10.1007/s00234-021-02718-7.

64.	Ojha R, Jiang A, Mäntylä E, Quirin T, Modhira N, Witte R, et al. Dynamin independent endocytosis is an alternative cell entry mechanism for multiple animal viruses. PLoS Pathog. 2024;20:e1012690. doi:10.1371/journal.ppat.1012690.

65.	R Core Team. R: A language and environment for statistical computing. Vienna, Austria: R Foundation for Statistical Computing; 2024.

66.	The Gene Ontology Consortium. The Gene Ontology knowledgebase in 2026. Nucleic Acids Res. 2026;54:D1779-D1792. doi:10.1093/nar/gkaf1292.

67.	Ashburner M, Ball CA, Blake JA, Botstein D, Butler H, Cherry JM, et al. Gene ontology: tool for the unification of biology. The Gene Ontology Consortium. Nat Genet. 2000;25:25–9. doi:10.1038/75556.

68.	R Core Team. R: A Language and Environment for Statistical Computing. Vienna, Austria: R Foundation for Statistical Computing; 2025.

69.	Bengtsson H. matrixStats: Functions that Apply to Rows and Columns of Matrices (and to vectors); 2025.

70.	Cui A, Huang T, Li S, Ma A, Pérez JL, Sander C, et al. Dictionary of immune responses to cytokines at single-cell resolution. Nature. 2024;625:377–84. doi:10.1038/s41586-023-06816-9.

71.	Hughes CE, Nibbs RJB. A guide to chemokines and their receptors. FEBS J. 2018;285:2944–71. doi:10.1111/febs.14466.

72.	Matsuoka T, Narumiya S. Prostaglandin receptor signaling in disease. ScientificWorldJournal. 2007;7:1329–47. doi:10.1100/tsw.2007.182.

73.	Nakamura M, Shimizu T. Leukotriene receptors. Chem Rev. 2011;111:6231–98. doi:10.1021/cr100392s.

74.	Sánchez-García S, Jaén RI, Fernández-Velasco M, Delgado C, Boscá L, Prieto P. Lipoxin-mediated signaling: ALX/FPR2 interaction and beyond. Pharmacol Res. 2023;197:106982. doi:10.1016/j.phrs.2023.106982.

75.	Coyne E, Nie Y, Abdurrachim D, Ong CLZ, Zhou Y, Ali AAB, et al. Leukotriene B4 receptor 1 (BLT1) does not mediate disease progression in a mouse model of liver fibrosis. Biochem J. 2023;481:177–90. doi:10.1042/BCJ20230422.

76.	Maravillas-Montero JL, Burkhardt AM, Hevezi PA, Carnevale CD, Smit MJ, Zlotnik A. Cutting edge: GPR35/CXCR8 is the receptor of the mucosal chemokine CXCL17. J Immunol. 2015;194:29–33. doi:10.4049/jimmunol.1401704.

77.	Ineichen BV, Okar SV, Proulx ST, Engelhardt B, Lassmann H, Reich DS. Perivascular spaces and their role in neuroinflammation. Neuron. 2022;110:3566–81. doi:10.1016/j.neuron.2022.10.024.

78.	Marchetti L, Engelhardt B. Immune cell trafficking across the blood-brain barrier in the absence and presence of neuroinflammation. Vasc Biol. 2020;2:H1-H18. doi:10.1530/VB-19-0033.

79.	Gautam J, Zhang X, Yao Y. The role of pericytic laminin in blood brain barrier integrity maintenance. Scientific reports. 2016;6:36450. doi:10.1038/srep36450.

80.	Alvarez JI, Kébir H, Cheslow L, Charabati M, Chabarati M, Larochelle C, Prat A. JAML mediates monocyte and CD8 T cell migration across the brain endothelium. Ann Clin Transl Neurol. 2015;2:1032–7. doi:10.1002/acn3.255.

81.	Bouwman AC, van Daalen KR, Crnko S, Broeke T ten, Bovenschen N. Intracellular and Extracellular Roles of Granzyme K. Front Immunol. 2021;12:677707. doi:10.3389/fimmu.2021.677707.

82.	Voskoboinik I, Whisstock JC, Trapani JA. Perforin and granzymes: function, dysfunction and human pathology. Nat Rev Immunol. 2015;15:388–400. doi:10.1038/nri3839.

83.	Odobasic D, Kitching AR, Yang Y, O'Sullivan KM, Muljadi RCM, Edgtton KL, et al. Neutrophil myeloperoxidase regulates T-cell-driven tissue inflammation in mice by inhibiting dendritic cell function. Blood. 2013;121:4195–204. doi:10.1182/blood-2012-09-456483.

84.	Knox EG, Aburto MR, Clarke G, Cryan JF, O'Driscoll CM. The blood-brain barrier in aging and neurodegeneration. Mol Psychiatry. 2022;27:2659–73. doi:10.1038/s41380-022-01511-z.

85.	Zhao Z, Nelson AR, Betsholtz C, Zlokovic BV. Establishment and dysfunction of the blood-brain barrier. Cell. 2015;163:1064–78. doi:10.1016/j.cell.2015.10.067.

86.	Wareham LK, Baratta RO, Del Buono BJ, Schlumpf E, Calkins DJ. Collagen in the central nervous system: contributions to neurodegeneration and promise as a therapeutic target. Mol Neurodegener. 2024;19:11. doi:10.1186/s13024-024-00704-0.

87.	Halder SK, Sapkota A, Milner R. The importance of laminin at the blood-brain barrier. Neural Regen Res. 2023;18:2557–63. doi:10.4103/1673-5374.373677.

88.	Grant RI, Hartmann DA, Underly RG, Berthiaume A-A, Bhat NR, Shih AY. Organizational hierarchy and structural diversity of microvascular pericytes in adult mouse cortex. Journal of Cerebral Blood Flow & Metabolism. 2019;39:411–25. doi:10.1177/0271678X17732229.

89.	He J, Siahaan TJ, Kuczera K. Systematic Search for Blood-Brain Barrier Modulating Peptides Based on Exhaustive E-Cadherin Domain-Domain Docking. J Chem Inf Model. 2025;65:10688–700. doi:10.1021/acs.jcim.5c01227.

90.	Li X, Ding W, Zhang X, Tang X, Zheng Y, Ma Y, et al. Caveolin-1-Mediated Blood-Brain Barrier Disruption via MMP2/9 Contributes to Postoperative Cognitive Dysfunction. Neurochem Res. 2025;50:210. doi:10.1007/s11064-025-04458-z.

91.	Badaut J, Blochet C, Obenaus A, Hirt L. Physiological and pathological roles of caveolins in the central nervous system. Trends Neurosci. 2024;47:651–64. doi:10.1016/j.tins.2024.06.003.

92.	Sowa G. Caveolae, caveolins, cavins, and endothelial cell function: new insights. Front Physiol. 2012;2:120. doi:10.3389/fphys.2011.00120.

93.	Gatseva A, Sin YY, Brezzo G, van Agtmael T. Basement membrane collagens and disease mechanisms. Essays Biochem. 2019;63:297–312. doi:10.1042/EBC20180071.

94.	Bader BL, Smyth N, Nedbal S, Miosge N, Baranowsky A, Mokkapati S, et al. Compound genetic ablation of nidogen 1 and 2 causes basement membrane defects and perinatal lethality in mice. Mol Cell Biol. 2005;25:6846–56. doi:10.1128/MCB.25.15.6846-6856.2005.

95.	Nakamura K, Ikeuchi T, Nara K, Rhodes CS, Zhang P, Chiba Y, et al. Perlecan regulates pericyte dynamics in the maintenance and repair of the blood-brain barrier. J Cell Biol. 2019;218:3506–25. doi:10.1083/jcb.201807178.

96.	Suzuki Y, Nampei M, Kawakita F, Oinaka H, Nakajima H, Suzuki H. The effect of Fibulin-5 on early brain injury after subarachnoid hemorrhage in mice. Neurochem Int. 2025;187:105989. doi:10.1016/j.neuint.2025.105989.

97.	Hoegg-Beiler MB, Sirisi S, Orozco IJ, Ferrer I, Hohensee S, Auberson M, et al. Disrupting MLC1 and GlialCAM and ClC-2 interactions in leukodystrophy entails glial chloride channel dysfunction. Nat Commun. 2014;5:3475. doi:10.1038/ncomms4475.

98.	Idkowiak J, Dehairs J, Schwarzerová J, Olešová D, Truong JXM, Kvasnička A, et al. Best practices and tools in R and Python for statistical processing and visualization of lipidomics and metabolomics data. Nat Commun. 2025;16:8714. doi:10.1038/s41467-025-63751-1.

99.	Daigle TL, Madisen L, Hage TA, Valley MT, Knoblich U, Larsen RS, et al. A Suite of Transgenic Driver and Reporter Mouse Lines with Enhanced Brain-Cell-Type Targeting and Functionality. Cell. 2018;174:465-480.e22. doi:10.1016/j.cell.2018.06.035.

100.	Lein ES, Hawrylycz MJ, Ao N, Ayres M, Bensinger A, Bernard A, et al. Genome-wide atlas of gene expression in the adult mouse brain. Nature. 2007;445:168–76. doi:10.1038/nature05453.

101.	Harris JA, Mihalas S, Hirokawa KE, Whitesell JD, Choi H, Bernard A, et al. Hierarchical organization of cortical and thalamic connectivity. Nature. 2019;575:195–202. doi:10.1038/s41586-019-1716-z.

102.	Oh SW, Harris JA, Ng L, Winslow B, Cain N, Mihalas S, et al. A mesoscale connectome of the mouse brain. Nature. 2014;508:207–14. doi:10.1038/nature13186.
